# Comparative proximity biotinylation implicates RAB18 in sterol mobilization and biosynthesis

**DOI:** 10.1101/871517

**Authors:** Robert S. Kiss, Jarred Chicoine, Youssef Khalil, Robert Sladek, He Chen, Alessandro Pisaturo, Cyril Martin, Jessica D. Dale, Tegan A. Brudenell, Archith Kamath, Emanuele Paci, Anja Kerksiek, Dieter Lütjohann, Peter Clayton, Jimi C. Wills, Alex von Kriegsheim, Tommy Nilsson, Eamonn Sheridan, Mark T. Handley

**Author notes:** CORRESPONDING AUTHORS Robert S. Kiss Mark T. Handley. authors contributed equally.

## Abstract

Loss of functional RAB18 causes the autosomal recessive condition Warburg Micro syndrome. To better understand this disease, we used proximity biotinylation to generate an inventory of potential RAB18 effectors. A restricted set of 28 RAB18-interactions were dependent on the binary RAB3GAP1-RAB3GAP2 RAB18-guanine nucleotide exchange factor (GEF) complex. 12 of these 28 interactions are supported by prior reports and we have directly validated novel interactions with SEC22A, TMCO4 and INPP5B. Consistent with a role for RAB18 in regulating membrane contact sites (MCSs), interactors included groups of microtubule/membrane-remodelling proteins, membrane-tethering and docking proteins, and lipid-modifying/transporting proteins. Two of the putative interactors, EBP and OSBPL2/ORP2, have sterol substrates. EBP (emopamil binding protein) is a Δ8-Δ7 sterol isomerase and OSBPL2/ORP2 is a lipid transport protein. This prompted us to investigate a role for RAB18 in cholesterol biosynthesis. We find that the cholesterol precursor and EBP-product lathosterol accumulates in both RAB18-null HeLa cells and RAB3GAP1-null fibroblasts derived from an affected individual. Further, *de novo* cholesterol biosynthesis is impaired in cells in which RAB18 is absent or dysregulated. Our data demonstrate that GEF-dependent Rab-interactions are highly amenable to interrogation by proximity biotinylation and may suggest that Micro syndrome is a cholesterol biosynthesis disorder.

## INTRODUCTION

Rab Proteins are a large subfamily of small GTPases with discrete roles in coordinating membrane trafficking (1). Like other small GTPases, they adopt different conformations and enter into different protein-protein interactions according to whether they are GDP-, or GTP-bound. Although they possess some intrinsic GTP-hydrolysis activity, their nucleotide-bound state in cells is tightly governed by two classes of regulatory proteins. Guanine-nucleotide exchange factors (GEFs) catalyse the exchange of bound GDP for GTP while GTPase-activating proteins (GAPs) promote the hydrolysis of bound GTP to GDP (2, 3).

Biallelic loss-of-function variants in *RAB18*, *RAB3GAP1*, *RAB3GAP2*, or *TBC1D20*, cause the autosomal recessive condition Warburg Micro syndrome (4–8)(MIMs 600118, 614222, 614225, 615663, 212720). *RAB3GAP1* and *RAB3GAP2* encode subunits of the binary RAB18-GEF complex ‘RAB3GAP’, whereas *TBC1D20* encodes a RAB18-GAP (9, 10). Thus, the same pathology is produced when functional RAB18 is absent or when its normal regulation is disrupted. However, it is unclear how RAB18 dysfunction contributes to disease pathology at a molecular level.

Rab proteins fulfil their roles by way of protein-protein interactions with interacting partners termed ‘effectors’. The identification of these proteins can therefore provide insight into these roles. However, biochemical identification of Rab effectors is challenging; Rab-effector interactions are usually GTP-dependent and are often highly transient. Immunoprecipitation, affinity purification and yeast-2-hybrid approaches have each been used, but may be more or less effective depending on the Rab isoform studied (11, 12).

One newer approach is ‘BioID’ proximity biotinylation utilizing Rab proteins fused to mutant forms of the biotin ligase BirA. The Rab fusion protein biotinylates proximal proteins which are then purified on streptavidin and identified through mass spectrometry (13–15). Biotin labelling occurs in a relatively physiological context and prospective effectors can be purified under high stringency conditions. However, a drawback of the technique is that it does not distinguish between close associations resulting from functional protein-protein interactions and those resulting from overlapping localizations.

To discriminate functional RAB18 interactions, we compared BirA*-RAB18 labelling of protein in wild-type HeLa cells to that in cells in which RAB18-GEF activity was disrupted with CRISPR. Known and novel effectors were more strongly labelled in the wild-type cells. 28 RAB18-interactions were categorized as RAB3GAP-dependent. These proteins comprised several groups. Proteins within each group were clearly interrelated through involvement in connected biological processes. Moreover, gene-disease associations within the set included multiple overlapping phenotypes.

Our data elaborate an existing model suggesting that RAB18 effectors act collectively in lipid transfer at membrane contact sites (16). We identify multiple proteins already implicated in the establishment and maintenance of membrane-contacts. We verify novel interactions with SEC22A, TMCO4 and INPP5B using immunoprecipitation of exogenously expressed fusion proteins. We also identify putative RAB18-interactors involved in sterol biosynthesis and mobilization. In particular, a putative interaction with the Δ8-Δ7 sterol isomerase enzyme EBP led us to examine sterol profiles and cholesterol biosynthesis in several cell lines. We find that a sterol product of EBP-catalysis – lathosterol - accumulates in RAB18-null HeLa cells and RAB3GAP1-null human fibroblasts. Further, that cholesterol biosynthesis is reduced in cells in which RAB18 is absent or dysregulated. Because Micro syndrome shares a number of features with known cholesterol biosynthesis disorders, these data provide a tentative indication that this deficit might partly underlie disease pathology.

## RESULTS

### An inventory of RAB18-GEF-dependent RAB18-associated proteins in HeLa cells

We first used CRISPR to generate a panel of clonal, otherwise isogenic, HeLa cell lines null for RAB18 and a number of its regulators (see Figure S1). We then carried out proximity labelling using transient expression of the same exogenous BirA*-RAB18 construct in RAB3GAP1-, RAB3GAP2- and TRAPPC9-null cell lines and in wild-type cells (Figure 1A). RAB3GAP1 and RAB3GAP2 are each essential subunits of a binary RAB18-GEF complex (9). TRAPPC9 is reported to be essential for the RAB18-GEF activity of a different GEF, the multi-subunit TRAPPII complex (17).

**Figure 1.**
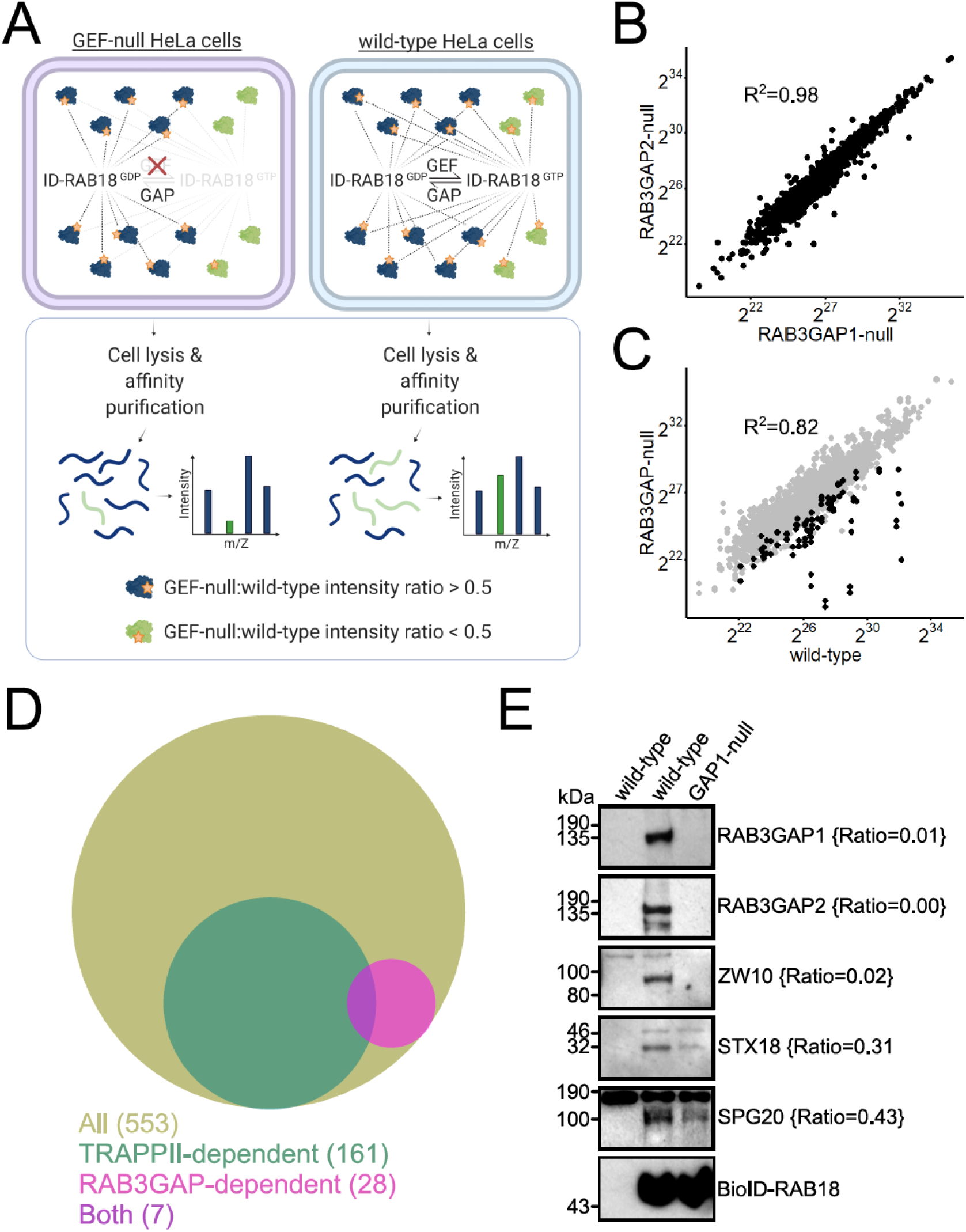
RAB3GAP-dependent RAB18-interactions in HeLa cells. (A) Schematic to show experimental approach. Proximity biotinylation of guanine nucleotide exchange factor (GEF)-dependent interactors by BirA*-RAB18 (ID-RAB18) is disrupted in GEF-null cells. GEF-independent interactors are biotinylated in both GEF-null and wild-type cells. Following affinity purification, GEF-dependent interactions are determined by LFQ intensity ratios. (B) Plot to show correlation between Log_2_ LFQ intensities of individual proteins identified in samples purified from RAB3GAP1- and RAB3GAP2-null cells. (C) Plot to show correlation between Log_2_ LFQ intensities of individual proteins identified in samples purified from wild-type and RAB3GAP-null cells. Highlighted datapoints correspond to proteins later found to have RAB3GAP-null:wild-type intensity ratios ≤0.5. (D) Venn diagram to show overlap between all RAB18-associations, TRAPPII-dependent interactions (TRAPPC9-null:wild-type intensity ratios <0.5) and RAB3GAP-dependent associations (RAB3GAP-null:wild-type intensity ratios <0.5). (E) Western blotting of samples purified from wild-type and RAB3GAP1-null cells in an independent BioID experiment. Levels of selected proteins are consistent with RAB3GAP-null:wild-type intensity ratios {braces}.

Proximity-labelling, affinity purification and mass spectrometry of biotinylated proteins were carried out essentially as previously described (15, 18). Prior to mass-spec analysis, samples from each of the streptavidin pull-downs were subjected to Western blotting to ensure comparable BirA*-RAB18 expression (Figure S2A). Label-free quantitative proteomics (LFQP) analyses were used to calculate ‘LFQ intensities’ for each RAB18-associated protein (19). These were then normalized in each experiment according to the quantity of RAB18 found in each sample. Samples from three independent experiments were analysed. Pull-downs from untransfected biotin-treated cells were used as controls. After filtering the data to remove known mass-spec contaminants and any protein identified at a high level in control samples, a total of 902, 635 and 661 RAB18-associated proteins were identified in each experiment. A total of 553 proteins were present in two or more of the replicate experiments (see Table S1).

Different Rab-GEF complexes may operate in distinct subcellular localizations and coordinate associations with different effectors (20). Therefore, we assessed whether non-zero intensities for each RAB18-associated protein correlated between samples (Figure 1B, Figure S2B). Very strong correlations between protein intensities from RAB3GAP1- and RAB3GAP2-null cells indicated that loss of either protein had a functionally equivalent effect (R^2^=0.98, Figure 1B). In contrast, intensities from RAB3GAP1/2- and TRAPPC9-null cells were much more poorly correlated (R^2^=0.74, Figure S2B). We therefore considered RAB3GAP- and TRAPPC9-dependent RAB18-interactions separately. Intensities from wild-type and RAB3GAP-null samples correlated with an R^2^=0.82, but a number of proteins showed reduced intensities in the RAB3GAP-null samples (Figure 1C).

GEF activity promotes Rab GTP-binding and this is usually necessary for effector interactions. We therefore reasoned that levels of true effector proteins would be reduced in samples from GEF-null cells as compared to those from wild-type cells (Figure 1A). We calculated GEF-null:wild-type intensity ratios for each RAB18-associated protein (Table S1). Only 28 proteins showed a RAB3GAP-null:wild-type ratio ≤0.5 (Figure 1D, Table 1, Table S1). 161 proteins showed a TRAPPII-null:wild-type intensity ratio ≤0.5 (Figure 1D, Table S1). There was only limited overlap between RAB3GAP- and TRAPPC9-dependent associations (Figure 1D).

**Table 1.**
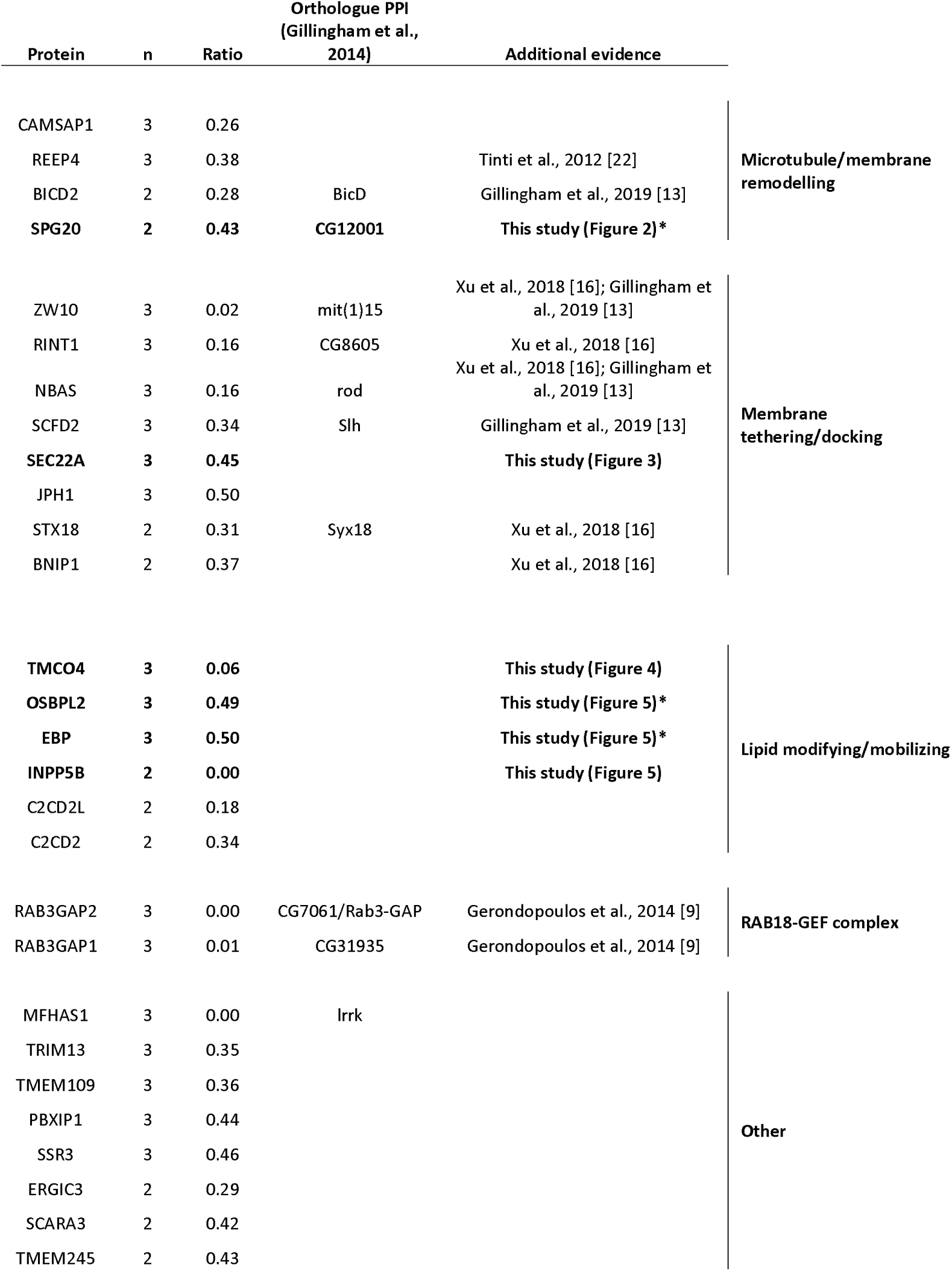
RAB3GAP-dependent RAB18-interactions in HeLa cells. 28 proteins with mean RAB3GAP-null:wild-type intensity ratios ≤0.5, identified in two or more independent proximity biotinylation experiments. Orthologous proteins identified by Gillingham et al., 2014, and other studies providing supporting evidence for interactions are shown. Proteins are grouped according to their reported functions. The full dataset is provided in Table S1. *indirect additional evidence for this interaction is presented in this study.

The most comprehensive annotation of candidate RAB18 effectors thus far was made in the 2014 paper by Gillingham et al., which utilized an affinity purification-mass spectrometry (AP-MS) approach and the *Drosophila* RAB18 orthologue (21). In that study, a total of 456 proteins were identified as interacting with RAB18. However, only 14 of these were well represented in terms of spectral counts, exhibited low non-specific binding to GST/sepharose and showed low binding to other Rab protein isoforms. We took these 14 proteins as the most plausible physiological RAB18 interactors and searched for these in our datasets.

Orthologues or paralogs of 11 of the 14 putative RAB18-interacting proteins identified by Gillingham et al. were identified as GEF-dependent in our combined dataset. 10/14 were among the 28 RAB3GAP-dependent associations (listed in Table 1). 4/14 were among the TRAPPII-dependent associations (Table S1). 9/14 interactors from the Gillingham et al. study and 12/28 of the RAB3GAP-dependent associations from our study have been described in other previously published work (9,13,16,22).

For initial validation of our dataset and the reproducibility of our results, we carried out an additional independent BioID experiment with wild-type and RAB3GAP1-null cells and subjected the resulting samples to Western blotting for selected RAB18-associated proteins (Figure 1E). As with the mass spectrometry, these proteins showed either complete (RAB3GAP2, ZW10) or partial (SPG20, STX18) dependence on RAB3GAP for their RAB18 association.

We further validated our approach with additional proximity biotinylation experiments in HEK293 cells. We used cells stably expressing BirA*-tagged RAB18 fusions incorporating wild-type RAB18, GTP-hydrolysis deficient RAB18(Gln67Leu), or nucleotide-binding deficient RAB18(Ser22Asn) mutants (Figure S3A-B). A total of 96 proteins were identified as associating with RAB18 across all samples (Table S2). Gln67Leu:wild-type intensity ratios for known RAB18-interactors ranged from 0.1-1.49 indicating that RAB18 associations were altered by the Gln67Leu variant, but not predictably so. In contrast, Ser22Asn:wild-type intensity ratios were <0.5 for the majority of these proteins. 28 nucleotide-binding-dependent RAB18 associations included 5 of the RAB3GAP-dependent associations and 7 of the TRAPPII-dependent associations seen in the HeLa cells (Figure S3C). These data confirm that the loss of GEFs has similar effects on RAB18-interactions to direct loss of nucleotide binding. In addition, they support the differing regulation of specific RAB18-interactions by different GEFs.

### Validation screening of RAB3GAP-dependent RAB18 associations reveals reduced levels of SPG20 in RAB18-null and TBC1D20-null cells

Our continued study focused on the 28 RAB3GAP-dependent RAB18 associations identified in HeLa cells. Encouragingly, these appeared to share interconnected functions and fell into discrete groups (Table 1). Furthermore, genes encoding 7 of the 28 proteins or their homologues are associated with inherited diseases that share features with Micro syndrome (Table 2).

**Table 2.**
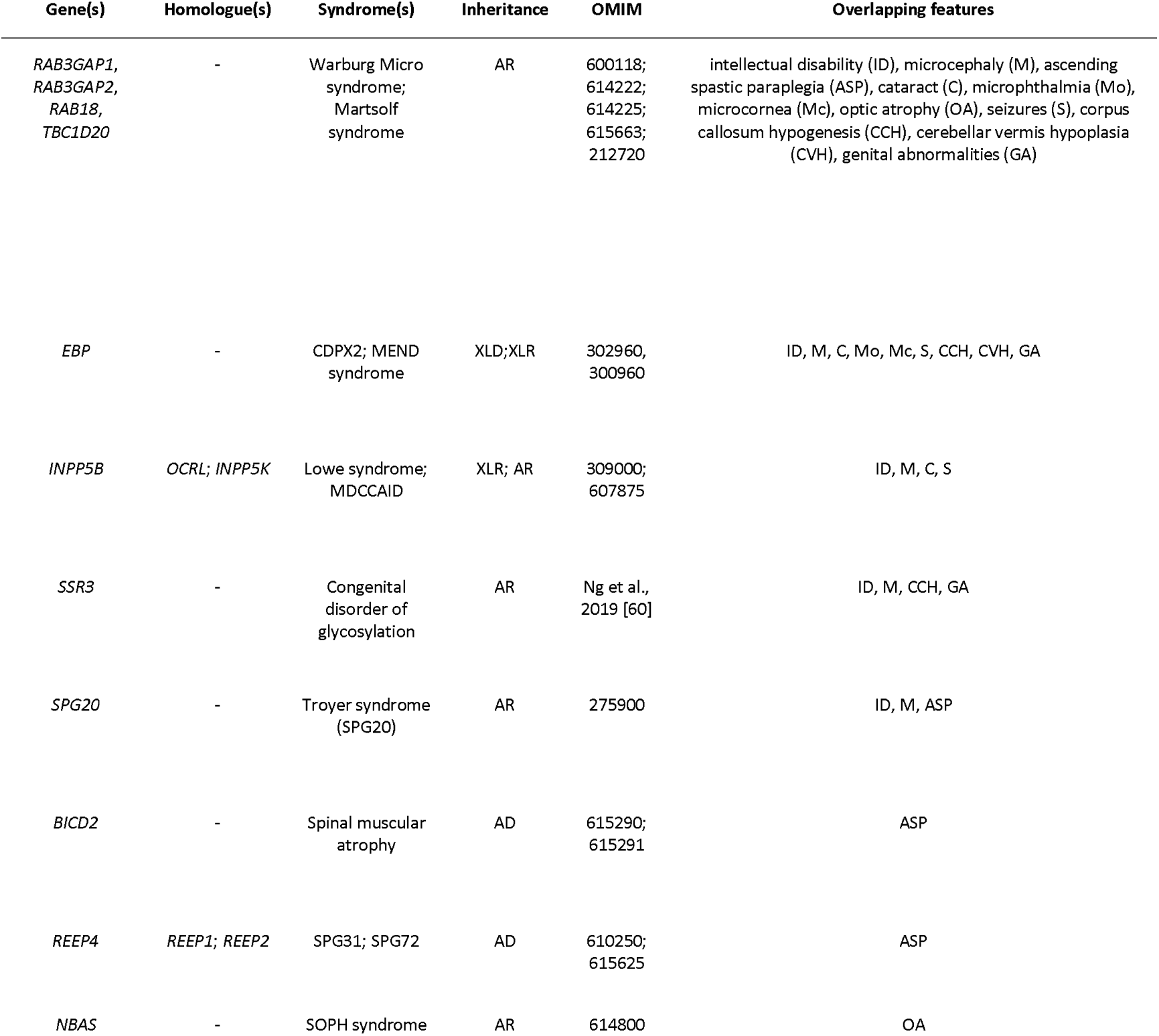
Genes encoding putative RAB18 effectors or their homologues are associated with diseases that share overlapping features with Warburg Micro syndrome. AR, autosomal recessive; XLD, X-linked dominant; XLR, X-linked recessive; AD, autosomal dominant.

Given the suggestive convergences in protein function and gene-disease-associations, we examined the subcellular localizations of 11 putative effectors for which antibodies were available (Figure 2A-B). For a rapid qualitative assessment of localizations we employed automated epifluorescence microscopy. The majority of antibodies used were validated in prior studies, or produced bands of the expected sizes when used in Western blotting (Table S6). However, antibodies for RINT1, C2CD2 and TRIM13 had only previously been manufacturer validated. To determine whether the localizations of the putative effectors were appreciably altered in cells lacking RAB18, we analysed wild type and RAB18-null lines in each case. In order to directly compare cells of different genotypes under otherwise identical conditions, we labelled them with CellTrace-Violet and CellTrace-Far Red reagents before seeding, immunostaining and imaging them together. Since RAB18 can localize to lipid droplets (LDs), we analysed both untreated cells (Figure 2A) and cells loaded with oleic acid and labelled with the LD marker BODIPY-558/568-C12 (Figure 2B).

**Figure 2.**
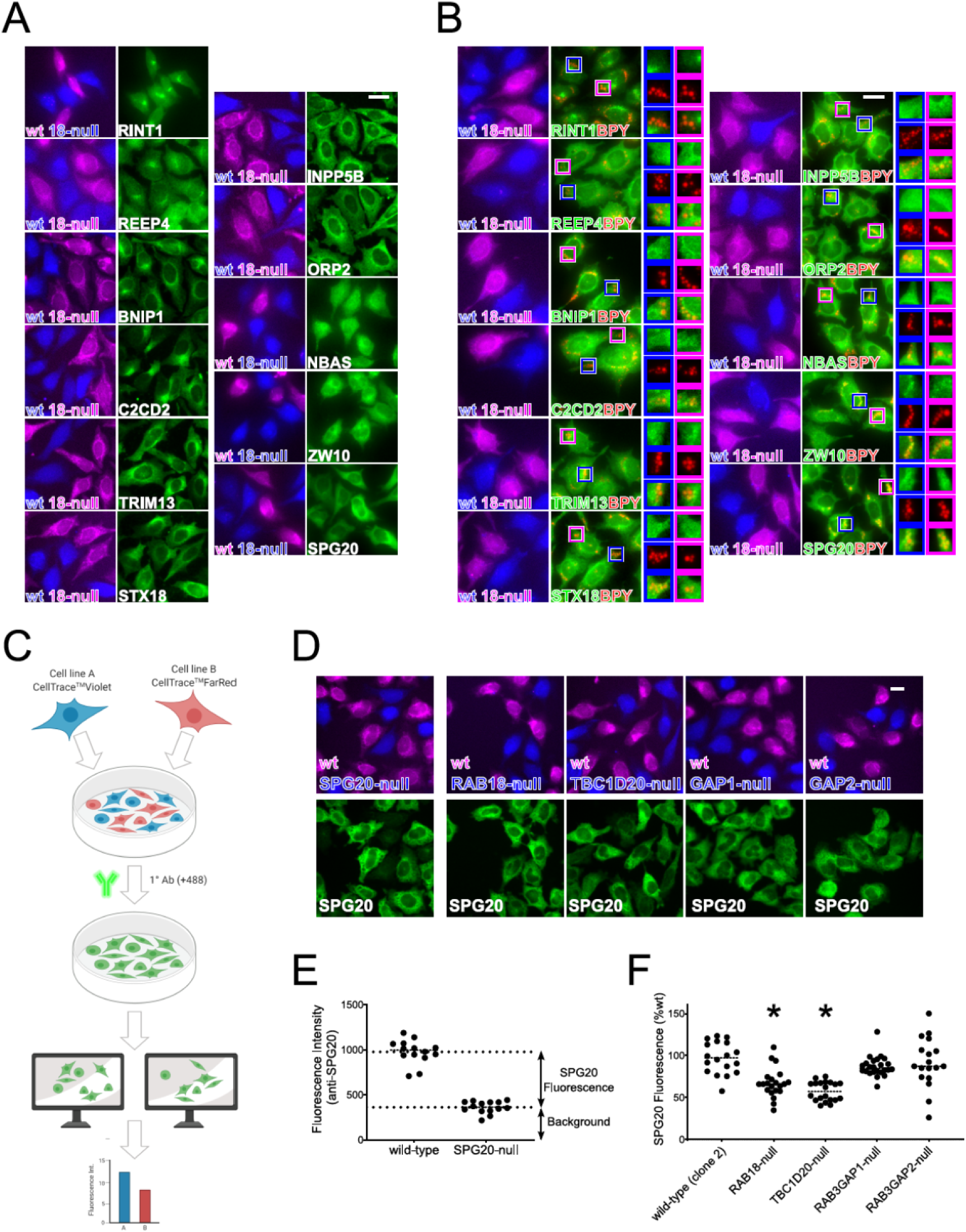
Initial screening of putative RAB18 effectors reveals that levels of SPG20 are significantly reduced in RAB18-null and TBC1D20-null cells. (A) Comparative fluorescence microscopy of selected RAB18-associated proteins in wild-type and RAB18-null HeLa cells. Cells of different genotypes were labelled with CellTrace-Violet and CellTrace-Far Red reagents, corresponding to blue and magenta channels respectively. Cells were stained with antibodies against indicated proteins in green channel panels. (B) Comparative fluorescence microscopy of selected RAB18-associated proteins in lipid-loaded wild-type and RAB18-null HeLa cells. Cells were stained as above but were treated for 15 hours with 200µM oleic acid, 1µg/ml BODIPY-558/568-C12 (Red channel; BPY) prior to fixation. (C) Schematic to show method for quantification of protein levels by fluorescence intensity. In each frame, cell areas for each genotype are generated by thresholding CellTrace channels, intensity of antibody-staining is measured for each cell in multiple frames. (D) Example frames showing wild-type and mutant cells of the indicated genotypes, labelled with CellTrace-Far Red and CellTrace-Violet and reagents respectively, then stained for SPG20. (E) Quantification of SPG20-specific fluorescence in wild-type cells by direct comparison with SPG20-null cells. (F) Quantification of SPG20 fluorescence (%wt) in cells of different genotypes. Data were derived from analysis of at least 18 frames – each containing >5 wild-type and >5 mutant cells – per genotype. *p<0.001. Bars 20µm.

The putative effector proteins showed various staining patterns. These ranged from staining that was enriched at the perinuclear region of cells, to staining that appeared reticular, to staining that appeared more diffuse. Staining patterns were similar in the HeLa cells and also in RPE1 cells generated to provide biological replicates (Figure S4A). Each pattern was compatible with the known localization of RAB18, which is distributed between *cis*-Golgi, ER, and cytosolic compartments (10). In lipid-loaded cells, localizations of proteins with reticular staining patterns overlapped with LDs but they did not obviously shift to adopt a predominantly LD localization. Two proteins that showed diffuse staining patterns in untreated cells - ZW10 and SPG20 - appeared enriched in the vicinity of LDs (Figure 2B, bottom right panels).

We saw no evidence for dramatic changes in protein localizations in RAB18-null cells as compared to their wild-type counterparts. Fluorescence intensities in RAB18-null and wild-type cells were also generally similar, except in the case of staining for SPG20, which appeared lower in RAB18-null HeLa cells than in wild-type cells (Figure 2A, bottom right panels).

To confirm the reduction in SPG20 fluorescence we observed in the RAB18-null HeLa cells, and to determine the effects of other genotypes, we used quantitative fluorescence microscopy (Figure 2C). To establish SPG20 antibody specificity we first analysed SPG20-null cells (Figure 2D, left panels). Measured background fluorescence intensity of these SPG20-null cells also provided a baseline level, above which fluorescence levels reflect the presence of SPG20 protein (Figure 2E). In RAB18-null cells, SPG20 fluorescence was reduced to 67.16±17.3%SD (p<0.001) of that in wild-type cells (Figure 2F). Loss of the RAB18-GEF subunits RAB3GAP1 or RAB3GAP2 had no significant effect, whereas loss of the RAB18-GAP TBC1D20 led to a reduction comparable to that seen in RAB18-null cells (57.48±11.48%SD, p<0.001)(Figure 2F).

We analysed levels of SPG20 in the corresponding panel of RPE1 cell lines using LFQP analysis of whole cell lysates (Figure S4B, Table S3). Levels of SPG20 were significantly reduced in RAB18- and TBC1D20-null RPE1 cells compared to wild-type controls (p<0.05 following FDR correction), but not in the other genotypes tested. These data suggest that these genotypes cause reduced SPG20 levels and that this is not the result of clonal variation. A comparison between LFQP data from wild-type and TBC1D20-null RPE1 and HeLa cells (Tables S3 and S4) showed limited overlap between differentially expressed proteins. This indicates that reduced SPG20 levels are unlikely to have resulted from widespread dysregulation of proteostasis. The RAB18-SPG20 interaction has been previously reported and validated (21), and our findings (above) provide further support for a physiological relationship between these proteins.

### SEC22A associates with RAB18 and its knockdown causes altered LD morphology

The most studied group of RAB18 effector proteins to date are the tethering factors that comprise the NRZ/Dsl complex (ZW10, NBAS and RINT1), and the ER SNARE proteins that comprise the Syntaxin18 complex (STX18, BNIP1, USE1 and SEC22B) (16,21,23,24). Although SNARE complexes typically mediate membrane fusion, it has been proposed that RAB18 interacts with these proteins to mediate the close apposition of membranes to facilitate lipid transfer (16). It has also been suggested that SEC22B is dispensable for this function (16).

Our screen for RAB3GAP-dependent RAB18-interactors identified all of the NRZ complex components, as well as the SNARE proteins STX18 and BNIP1 (Table 1). Interestingly, we did not identify SEC22B but did identify SEC22A among these proteins. SEC22A is one of the two SEC22B homologues in humans that lack the central coiled-coil SNARE domain through which SEC22B mediates membrane fusion (25). Since it had not been previously described as a RAB18-interacting protein, we investigated this further.

In the absence of appropriate commercially available antibodies for SEC22A, we examined its localization through expression of a mEmerald-SEC22A fusion protein (Figure 3A). mEmerald-SEC22A produced a characteristic reticular staining pattern and colocalized with an exogenous ER marker suggesting that SEC22A localizes to the ER. To verify the RAB18-SEC22A interaction, we carried out immunoprecipitation experiments after exogenous expression of mEmerald-SEC22A and/or HA-RAB18 fusion proteins (Figure 3B). mEmerald-SEC22A copurified together with HA-RAB18 in precipitates from wild-type but not RAB3GAP1-null cells. These data are consistent with a RAB3GAP-dependent interaction between RAB18 and SEC22A. However, we found that coexpression of mEmerald-SEC22A and mCherry-RAB18 disrupted normal ER morphology and produced vesicular structures and/or inclusions positive for both proteins in both wild-type and RAB3GAP-null cells (Figure 3C). Although not inconsistent with a functional protein-protein interaction, this precluded the use of coexpressed exogenous proteins in continued testing.

**Figure 3.**
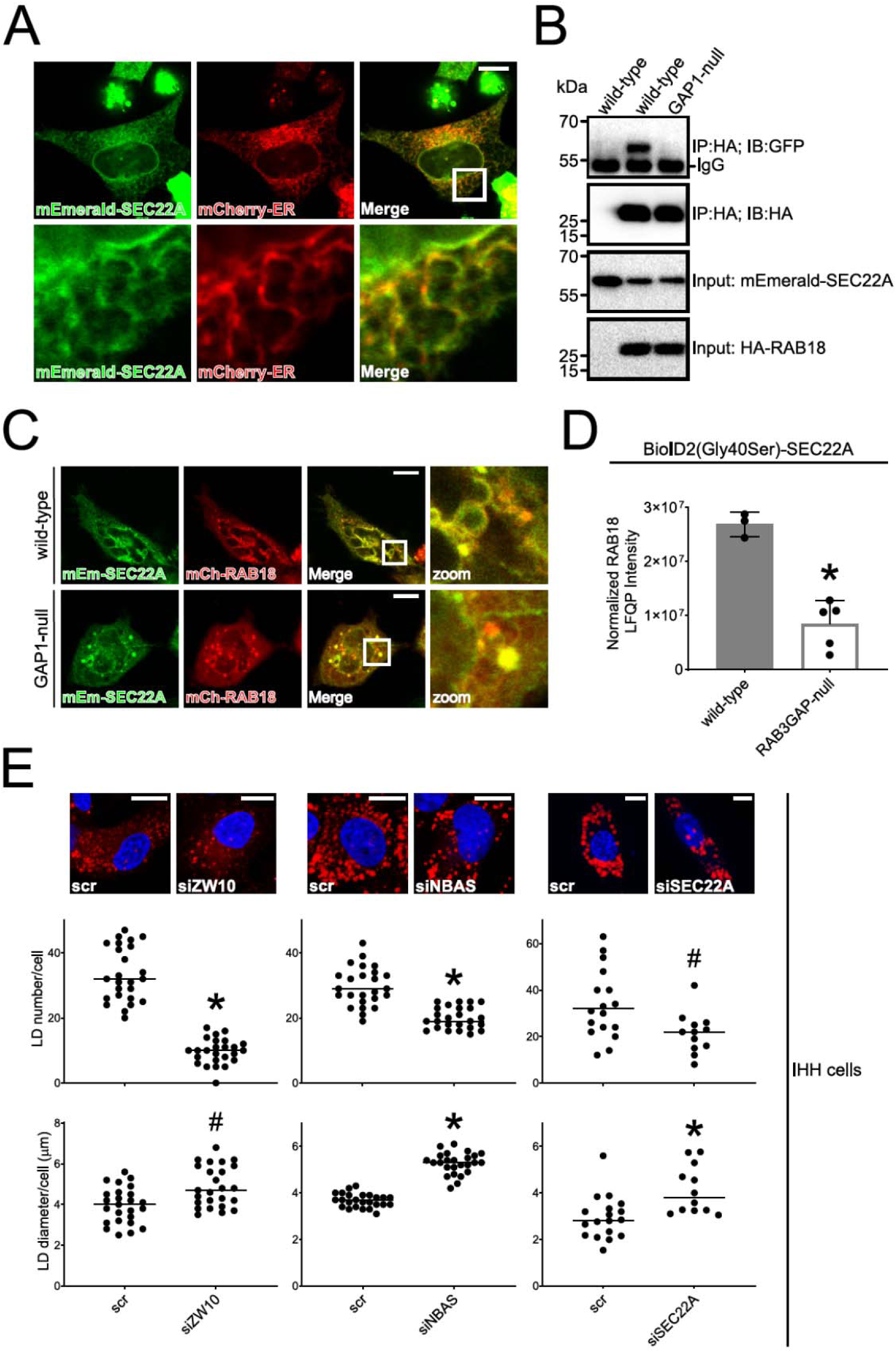
SEC22A associates with RAB18 and influences LD morphology. (A) Confocal micrograph to show overlapping localization of exogenous mEmerald-SEC22A (Green) and mCherry-ER (Red) in HeLa cells. (B) Immunoprecipitation of exogenous HA-RAB18 from wild-type and RAB3GAP1-null HeLa cells. Cells were transfected with HA-RAB18 and/or mEmerald-SEC22A and lysed 24 hours post-transfection. Anti-HA immunoprecipitates and input samples were subjected to SDS-PAGE and immunostaining for HA and GFP (mEmerald). (C) Confocal micrographs showing altered morphology in wild-type and RAB3GAP1-null HeLa cells coexpressing mEmerald-SEC22A and mCherry-RAB18; zoom shows co-labelled vesicular structures. (D) RAB18 LFQ intensities from a reciprocal BioID experiment showing a reduced association between BioID2(Gly40Ser)-SEC22A and endogenous RAB18 in RAB3GAP-null compared to wild-type HeLa cells. Data were adjusted to account for non-specific binding of RAB18 to beads and normalized by SEC22A LFQ intensities in each replicate experiment. Error bars represent SD. Data for other BioID2(Gly40Ser)-SEC22A-associated proteins are provided in table S5. (E) Example confocal micrographs and scatter plots to show effects of ZW10, NBAS and SEC22A knockdowns on lipid droplet number and diameter. siRNA-treated IHH cells were loaded with 200nM BSA-conjugated oleate, fixed and stained with BODIPY and DAPI, and imaged. Images were analysed using ImageJ. Data are representative of three independent experiments. ^#^p<0.05, *p<0.005. Bars 10µm.

As another means of assessing SEC22A-interactions, we used proximity biotinylation with a BirA*-SEC22A fusion protein in the HeLa cell panel. To minimize potential toxicity while increasing biotin-ligase activity, we used BioID2 (26) with a p.Gly40Ser active site modification (27) and reduced biotin incubation time. Despite a low level of BioID2(Gly40Ser)-SEC22A expression, the construct appeared to label RAB18 in a RAB3GAP-dependent manner (the labelling was reduced in RAB3GAP-null cells)(Figure 3D). 55 SEC22A-associated proteins were present in samples from wild-type cells in >2 replicate experiments and represented by >3 unique peptides (Table S5). Further, a subset of 9 SEC22A-associations were attenuated (intensity ratios <0.5) in samples from both RAB18-null and RAB3GAP-null cells.

A phenotype of altered LD morphology in lipid-loaded cells has been widely reported in cells deficient in RAB18 (8,9,16,17,28,29). Similar observations have been made in cells deficient in some components of the NRZ or Syntaxin18 complexes, but not in cells deficient in SEC22B (16). To test whether SEC22A expression influences LD morphology, we examined the effects of its silencing in oleic acid-loaded immortalized human hepatocyte (IHH) cells (Figure 3E). ZW10 and NBAS silencing provided positive controls in our experiments. ZW10 and NBAS silencing each led to a significant reduction in LD number (p<0.005) compared to controls, and a significant increase in LD size (p<0.05 and p<0.005 respectively). The effects of SEC22A silencing mirrored these findings, producing a significant reduction in LD number (p<0.05) and a significant increase in LD size (p<0.005). Together, these data implicate SEC22A as involved in the same RAB18-mediated process(es) as the NRZ and SNARE proteins.

### RAB18 recruits the orphan lipase TMCO4 to the ER membrane in a RAB3GAP-dependent manner

The most novel group of putative RAB18 effectors identified in our study were the lipid modifying/mobilizing proteins, none of which had been reported to associate with RAB18 previously. Among these, TMCO4 was identified in all three replicate experiments and its association with RAB18 was highly RAB3GAP-dependent (intensity ratio 0.06). Although annotated as containing transmembrane and coiled-coil domains, it is orthologous to the Yeast protein Mil1/ Yfl034w, and likely to be a partly soluble lipase (30). Consistently, TMCO4-EGFP expressed in HeLa cells showed a diffuse localization. In contrast, EGFP-RAB18 partly localizes to the ER, as shown by its colocalization with an ER marker (Figure 4A).

**Figure 4.**
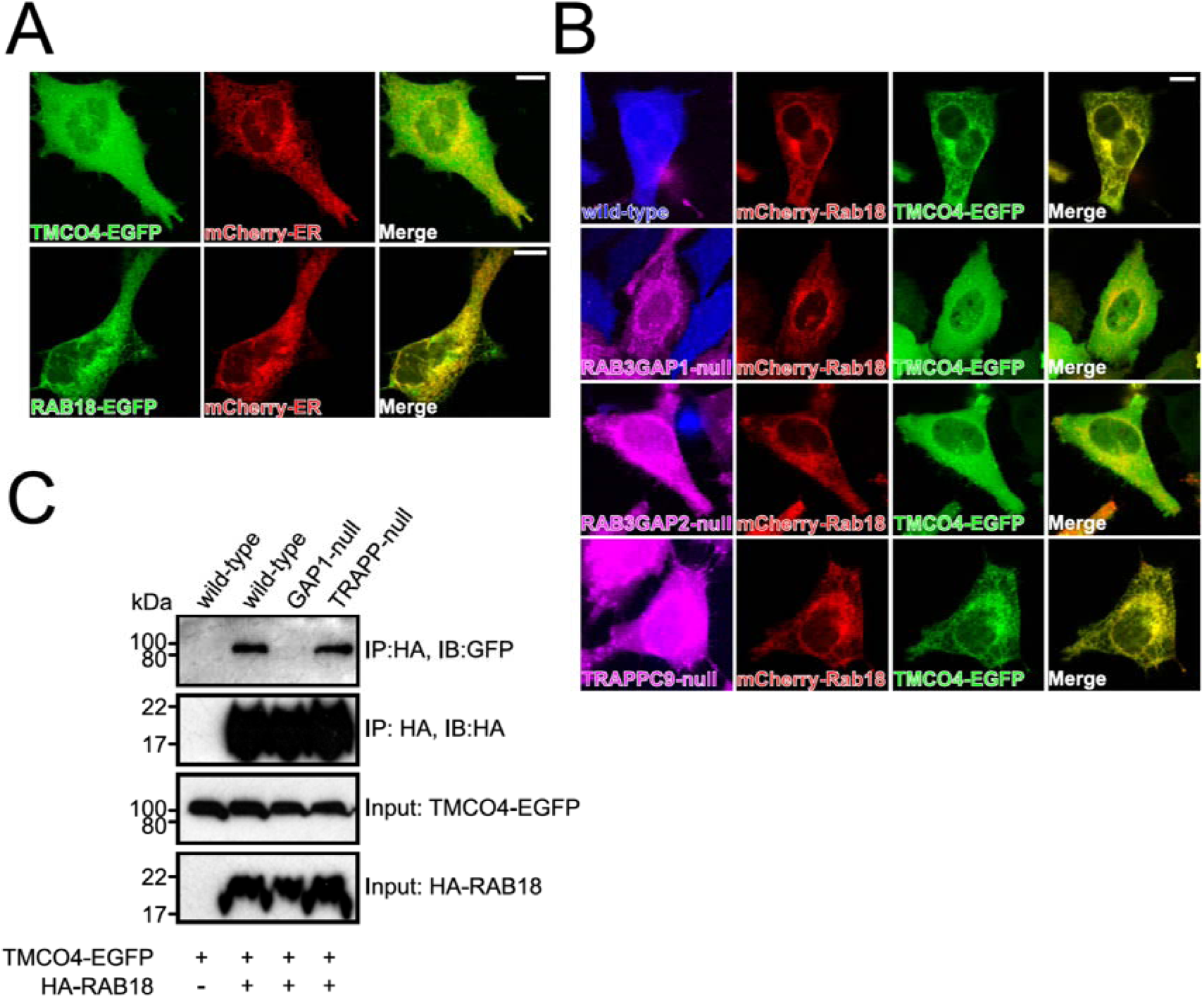
mCherry-RAB18 recruits TMCO4-EGFP to the ER membrane in a RAB3GAP-dependent manner. (A) Confocal micrographs to show diffuse localization of exogenous TMCO4-EGFP (Green) compared to mCherry-ER (Red) and overlapping localization of exogenous EGFP-RAB18 (Green) and mCherry-ER in HeLa cells. (B) Confocal micrographs to show localization of exogenous mCherry-RAB18 and TMCO4-EGFP in wild-type cells and in mutant cells of different genotypes. Wild-type and mutant cells of the indicated genotypes were labelled with CellTrace-Violet and CellTrace-Far Red reagents respectively (magenta and blue channels). (C) Immunoprecipitation of exogenous HA-RAB18 from HeLa cells of different genotypes. Cells were transfected with the indicated constructs and lysed 24 hours post-transfection. Anti-HA immunoprecipitates and input samples were subjected to SDS-PAGE and immunostaining for HA and GFP. Bars 10µm.

To assess the potential RAB18-TMCO4 interaction, we coexpressed mCherry-RAB18 and TMCO4-EGFP (Figure 4B). As in our previous experiments, Celltrace reagents were used to distinguish cells of wild-type and mutant genotypes. In wild-type HeLa cells, coexpression of mCherry-RAB18 led to a dramatic redistribution of TMCO4-EGFP to the ER membrane suggesting that RAB18 mediates recruitment of TMCO4 to this compartment. Redistribution was completely absent in RAB3GAP1- and RAB3GAP2-null cells but unaffected in TRAPPC9-null cells, consistent with the BioID data.

As a means of verifying the RAB18-TMCO4 interaction, we carried out immunoprecipitation experiments using exogenous HA-RAB18 and TMCO4-EGFP (Figure 4C). As expected, TMCO4-EGFP copurified with HA-RAB18 when expressed in wild-type or TRAPPC9-null cells, but not when expressed in RAB3GAP1-null cells. These data indicate that RAB18 and TMCO4 interact directly or indirectly as part of a protein complex in a RAB3GAP-dependent manner. Further, both the microscopy and the immunoprecipitation data support the suggestion that different GEFs can promote different RAB18-interactions.

### RAB18 is involved in cholesterol mobilization and biosynthesis

Other putative RAB18 effectors with lipid-related functions included ORP2/OSBPL2, INPP5B and EBP. ORP2 and INPP5B are robustly linked to a role in cholesterol mobilization. ORP2 is thought to function as a lipid transfer protein that exchanges cholesterol and PI(4, 5)P_2_ (31). INPP5B is implicated in the hydrolysis of PI(4, 5)P_2_, presumably driving the exchange process (31). EBP is involved in *de novo* cholesterol biosynthesis (32). In the Bloch pathway, it catalyses the conversion of 5α-cholesta-8, 24-dien-3β-ol (zymosterol) to 5α-cholesta-7, 24-dien-3β-ol (24-dehydrolathosterol). In the Kandutsch-Russel pathway, it catalyses the conversion of 5α-cholest-8(9)-en-3β-ol to 5α-cholest-7-en-3β-ol (lathosterol)(33). On the basis of these findings, we investigated the potential role of RAB18 in cholesterol uptake, efflux and biosynthesis.

We first performed loading and efflux experiments to measure the flux of cholesterol/cholesteryl ester (CE) while modifying the activity of RAB18. Chinese hamster ovary (CHO) cells were generated to stably express RAB18(WT), RAB18(Gln67Leu), or RAB18(Ser22Asn) (Figure S5A). In cells labelled with [^14^C]-oleate, but cholesterol-depleted with lipoprotein-depleted serum (LPDS), levels of CE were comparable in RAB18(Ser22Asn) and RAB18(WT) cells, whereas RAB18(Gln67Leu) cells stored significantly more (Figure 5A, left panel). In cells labelled with [^14^C]-oleate and cholesterol-loaded with FBS, levels of CE in RAB18(Ser22Asn) remained unchanged, whereas its storage was elevated in RAB18(WT) cells and RAB18(Gln67Leu) cells (Figure 5A, right panel). Interestingly, in both [^14^C]-oleate/LPDS and [^14^C]-oleate/FBS cells, the addition of high density lipoprotein (HDL, a vehicle mediating removal of cellular cholesterol) led to rapid depletion of CE in RAB18(Gln67Leu) cells, but not in RAB18(Ser22Asn) or RAB18(WT) cells (Figure 5A). Consistently, RAB18(Gln67Leu) cells also effluxed significantly more [^3^H]-sterol upon their incubation with apolipoprotein (apo) A-I than the other cell types (Figure 5B). These observations were not explained by altered expression of ABCA1, the transporter responsible for the rate-limiting step of cholesterol efflux (Figure S5B). These data suggest that ‘activated’ GTP-bound RAB18 strongly promotes the storage, turnover and mobilization of CE stored in LDs. A plausible explanation for this is that active RAB18 promotes cholesterol mobilization via ORP2 and INPP5B.

**Figure 5.**
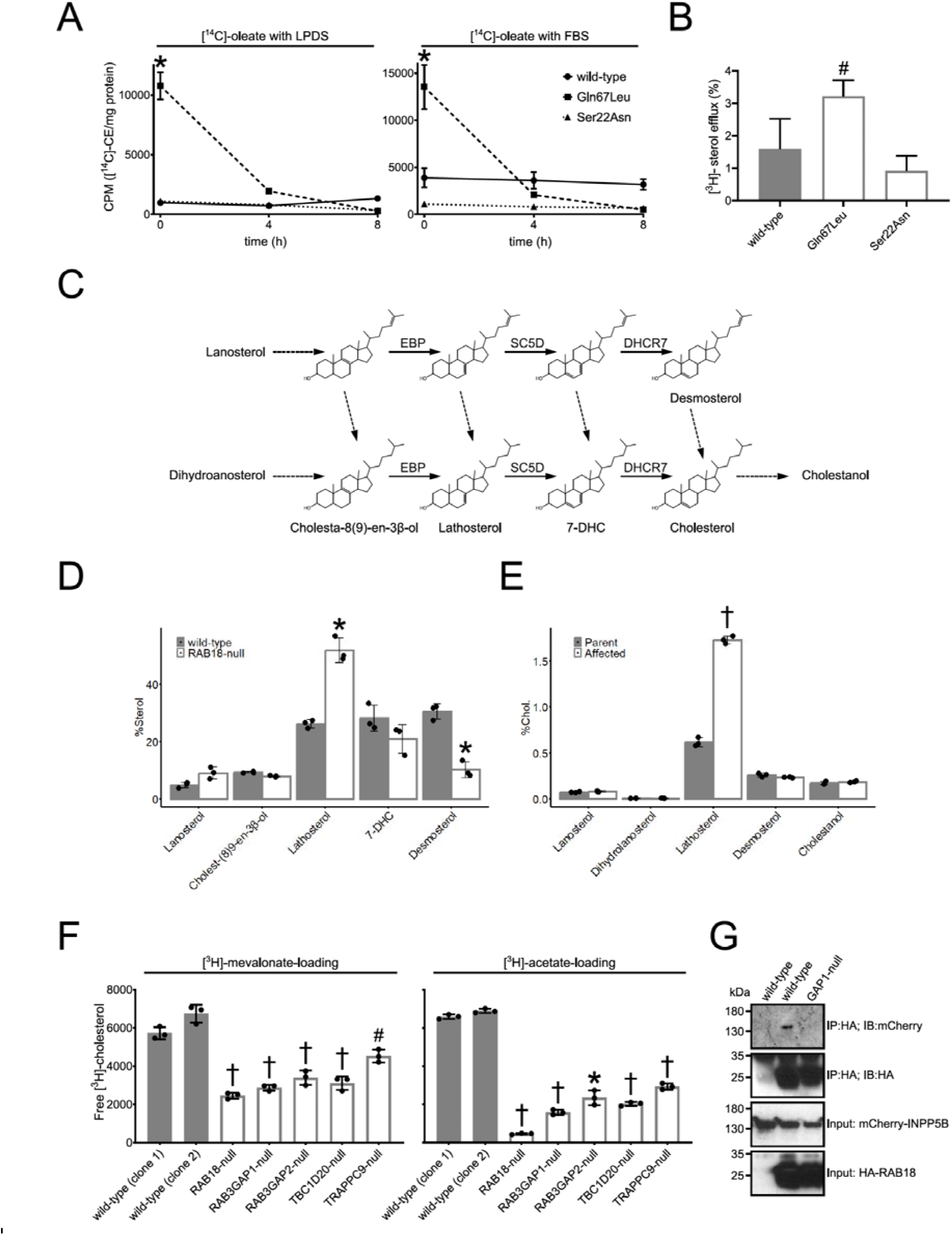
RAB18 is involved in the mobilization and biosynthesis of cholesterol. (A) Plots to show cholesteryl ester (CE) loading and efflux. CHO cells, stably expressing RAB18(WT), RAB18(Gln67Leu) and RAB18(Ser22Asn), were incubated with [^14^C]-oleate, for 24 hours, in the presence of lipoprotein depleted serum (LPDS)(Left panel) or FBS (Right panel). Following lipid extraction, thin layer chromatography (TLC) was used to separate CE, and radioactivity was measured by scintillation counting. Measurements were made at t=0 and at 4 and 8 hours following the addition of 50µg/ml high density lipoprotein (HDL) to the cells. (B) Bar graph to show cholesterol efflux. CHO cells were incubated with [^3^H]-cholesterol, for 24 hours, in the presence of FBS. After washing, they were incubated with 25µg/ml apolipoprotein A-I for 5 hours. The quantity of [^3^H]-sterol in the media is shown as a percentage of the total cellular radioactivity (mean±SD). (C) Schematic of post-squalene cholesterol biosynthesis pathway with the sterols quantified by gas chromatography-mass spectrometry-selected ion monitoring (GC-MS-SIM) named. Solid arrows indicate biosynthetic steps catalysed by EBP, SC5D and DHCR7. (D) Bar graph of sterols profile in wild-type and RAB18-null HeLa cells. Cells were grown in media supplemented with LPDS for 48 hours. Extracted sterols were analysed by GC-MS-SIM. % Sterol was calculated as a proportion of total quantified sterols, excluding cholesterol, following normalization to a 5α-cholestane internal standard. n=3; ±SD. (E) Bar graph of sterols profile in parental control fibroblasts and RAB3GAP1-deficient fibroblasts from an individual with Micro syndrome. Cells were grown in media supplemented with LPDS for 48 hours. Extracted sterols were analysed by GC-MS-SIM. % Cholesterol was calculated to express each quantified sterol as a proportion of total quantified cholesterol. n=3; ±SD. (F) Bar graphs to show incorporation of [^3^H]-mevalonate and [^3^H]-acetate into cholesterol in a panel of HeLa cell lines. Cells were grown in media supplemented with LPDS for 24 hours, then incubated with 5µCi/well [^3^H]-mevalonate or 10µCi/well [^3^H]-acetate for 24 hours. TLC was used to separate free cholesterol and radioactivity was quantified by scintillation counting (n=3; mean±SD). (G) Immunoprecipitation of HA-RAB18 from HeLa cells of different genotypes. Cells were transfected with the indicated constructs and lysed 24 hours post-transfection. Anti-HA immunoprecipitates and input samples were subjected to SDS-PAGE and immunostaining for HA and mCherry. ^#^p<0.05, *p<0.01, †p<0.0005.

Given that EBP functions in cholesterol biosynthesis, we next explored whether the absence of putative RAB18-reglulation of EBP might produce abnormal sterol profiles. A schematic of the cholesterol biosynthetic pathway is shown in Figure 5C. We incubated wild-type and RAB18-null HeLa cells for 48 hours in media supplemented with LPDS, then subjected samples to analysis by GC-MS-SIM (Figure 5D). In RAB18-null cells, we found that levels of the EBP-substrate cholest-8(9)-en-3β-ol were not significantly different from those in wild-type cells. In contrast, levels of EBP-product lathosterol were significantly higher (p<0.01). Moreover, levels of desmosterol - downstream of 24-dehydrolathosterol in the Bloch pathway - were significantly lower in RAB18-null cells (p<0.01). We extended our sterol profiling with additional experiments using RAB3GAP1-deficient fibroblasts from an individual with Micro syndrome together with control cells derived from a parent (Figure 5E). Following culturing with LPDS, as in the HeLa cells, we found that lathosterol levels were significantly higher in the RAB3GAP1-deficient fibroblasts than in the control cells (p<0.0005). Unlike in the HeLa cells, levels of desmosterol were not significantly different the between genotypes.

The finding that a product of EBP-catalysis, lathosterol, is significantly elevated in both HeLa cells and fibroblasts in which RAB18 function is impaired, provided good evidence that the RAB18-EBP interaction identified in our screening is meaningful. We reasoned that the elevated levels of a cholesterol precursor we observed in the RAB18-null and RAB3GAP1-deficient cells might be reflected in altered cholesterol biosynthesis when RAB18 is absent or dysregulated. In particular, the elevated lathosterol levels might reflect its accumulation due to perturbed transit through the biosynthetic pathway. To explore this possibility, we cultured the panel of HeLa cell lines for 24 hours in media supplemented with LPDS, treated them for 24 hours with [^3^H]-mevalonate or [^3^H]-acetate, then quantified labelled cholesterol (Figure 5F). Under both conditions, cholesterol synthesis in two clonal wild-type controls was comparable, but it was reduced in RAB18-, RAB3GAP1-, RAB3GAP2-, TBC1D20- and TRAPPC9-null cells. Levels of newly-produced cholesterol were lowest in the RAB18-null cells (39.5±2.5%SD and 6.8±0.5%SD of controls for [^3^H]-mevalonate and [^3^H]-acetate respectively). Levels in the cells of other genotypes were between 46±2.5%-73±5%SD for [^3^H]-mevalonate and 23±2%-43±2%SD for [^3^H]-acetate. These data strongly suggest that RAB18 and its regulators are required for normal cholesterol biosynthesis.

We attempted more direct verification of the putative interactions between RAB18 and ORP2, EBP and INPP5B using exogenously expressed fusion proteins and immunoprecipitation and GFP-Trap experiments. We did not detect the interactions with ORP2 or EBP, perhaps indicating that these are transient, weak, or disrupted by the lysis conditions or tags used. We were able to verify a RAB3GAP-dependent interaction between HA-RAB18 and mCherry-INPP5B using immunoprecipitation (Figure 5G).

## DISCUSSION

In this study, we have complemented previous work showing that proximity biotinylation is a powerful means of identifying candidate Rab effectors (13). Further – at least in the case of RAB18 - we have found that comparing biotin-labelling produced by a BirA*-Rab in wild-type and GEF-deficient cells can be particularly informative. We found that marked reductions in RAB18-association in RAB3GAP-null cells were restricted to only 28 proteins and that these comprised known and/or plausible effectors. We were able to exclude ∼94% of RAB18-associations from consideration as more likely to represent ‘noise’ from bystander proteins.

Prior evidence supported 12 of the 28 interactions we detected. Independent experiments with a mutant RAB18 fusion protein confirmed nucleotide-binding-dependence of several interactors, and immunofluorescence confirmed compatible localizations of several more. The known functions of the proteins were consistent with previous work implicating RAB18 in coordination of lipid exchange between apposed membranes (16). Further, gene-disease associations showed substantial overlap with RAB18-deficiency/Warburg Micro syndrome. We have presented additional validation by immunoprecipitation of novel interactions with SEC22A, TMCO4 and INPP5B (Figures 3B, 4C and 5G). Our more indirect data are consistent with functional interactions between RAB18 and its known interactor SPG20 (Figure 2D-F, Figure S4B), and novel putative interactors ORP2 and EBP (Figure 5A-F).

Together, our protein-interaction data implicate RAB18 in regulation of a stepwise process in which membrane/cytoskeletal remodelling precedes the engagement of tethering proteins and then SNAREs to establish membrane contact sites. The possible substitution of SEC22B for SEC22A in a RAB18-regulated Syntaxin18 SNARE complex, and a possible role for this complex in promoting membrane contacts rather than membrane fusion, is consistent with previous data (16). Further, it would be compatible with roles for the NRZ/Dsl1 complex and SCFD2/Sly1 in dynamically orchestrating SNARE complex assembly (34–36). More ambiguously, the RAB18-interacting microtubule binding proteins have not previously been reported to work together but do function in compatible locations. SPG20 and CAMSAP1 each associate with mitotic spindle poles, REEP4 participates in spindle-dependent ER clearance from metaphase chromatin and BICD2 is a component of the minus-end-directed dynein-dynactin motor complex (37–43). Our TRAPPII-dependent RAB18 interaction data indicate that different GEF complexes affect largely distinct subsets of interactions. However, more work will be required to determine whether these regulators mediate independent or interdependent functions.

Proteins implicated in lipid biology – particularly sterol biology – were prominent in our dataset. Consistent with the putative interaction between RAB18 and novel effectors ORP2 and INPP5B, which are reported to function in cholesterol mobilization (31), altered RAB18 activity was associated with altered cholesterol/cholesteryl ester mobilization in our experiments (Figure 5A-B). Other research implicates interactions between other Rab and ORP/OSBP isoforms in cholesterol mobilization at discrete sites (44–45), and several INPPs including INPP5B have a broad Rab-binding specificity (12-13,46). Thus, there may be a conserved relationship between these protein families, functioning in an analogous manner to the ARF1 GTPase, OSBP, and the phosphatase SACM1L (SAC1), in mediating sterol exchange (47).

Consistent with a functional interaction between RAB18 and EBP, levels of the EBP-product lathosterol were elevated in RAB18-null HeLa cells and in RAB3GAP1-deficient fibroblasts (Figure 5D-E). Cholesterol biosynthesis in HeLa cells was impaired when RAB18 was absent or dysregulated (Figure 5F). Given that ORP2 and INPP5B function in sterol mobilization whereas EBP functions in sterol biosynthesis, an attractive hypothesis is that RAB18 might coordinate their activities; that ORP2 might act as an exchanger for the products of EBP-catalysis as well as for cholesterol (Figure S6). In this case, defective mobilization of lathosterol would explain its accumulation in RAB18-null cells. Impaired delivery of substrates to downstream biosynthetic enzymes would explain the reduced cholesterol biosynthesis observed in these and the other model cell lines. The mobilization of cholesterol precursors by ORP proteins would not be unprecedented since the mobilization of cholesterol and its metabolites by these proteins is well established. Nevertheless, future work should aim to test this hypothesis more definitively. Among the other lipid-related proteins, TMCO4 may potentially be directly or indirectly associated with sterol metabolism. Although its substrate(s) are unknown, its expression is found to be upregulated in hypercholesterolemia (48) and it is present on lipid rafts (49). C2CD2L/TMEM24 and C2CD2, might potentially function in concert with ORP2 and/or INPP5B, since C2CD2L is found to mediate PI transport and to facilitate generation of PI(4, 5)P_2_ (50).

Our objective in studying RAB18 was to better understand the molecular pathology of Warburg Micro syndrome. Though our protein-interaction data are relatively preliminary, our functional findings represent good progress towards this goal. One key finding is that levels of lathosterol are significantly elevated in fibroblasts from an affected individual when these cells are cultured under LPDS (Figure 5E). In future work, we aim to determine whether this is reproducible in fibroblasts of other genotypes from other Micro syndrome individuals. If so, this could form the basis for a biochemical test for Micro syndrome which would complement genetic testing.

Another key finding is that disrupted *de novo* cholesterol biosynthesis may contribute to disease pathogenesis. Strongly supporting this suggestion, genes encoding multiple cholesterol biosynthesis enzymes are linked to similar disorders (33). For example, pathogenic variants in the lathosterol oxidase gene, *SC5D*, cause lathosterolosis, which is associated with microcephaly, intellectual disability, micrognathia, high arched palate and cataract (51–55). Pathogenic variants in the 7-dehydrocholesterol reductase gene, *DHCR7*, cause Smith-Lemli-Opitz syndrome (SLOS), which has a similar spectrum of features and is among the top differential diagnoses for Micro syndrome (7, 56). Indeed, the similarities with SLOS were noted in the report first identifying RAB18 as a disease-associated gene more than a decade ago (5).

## EXPERIMENTAL PROCEDURES

### Plasmids

The EGFP-RAB18 construct has been described previously (9). The RAB18 sequence was excised from this construct using BamHI and HindIII restriction enzymes (New England Biolabs, Hitchin, UK), and used to generate constructs encoding mEmerald-RAB18 and mCherry-RAB18 by ligation into mEmerald-C1 and mCherry-C1 vectors (Addgene, Watertown, MA) using HC T4 Ligase and rapid ligation buffer (Promega, Southampton, UK). Constructs encoding BirA*-RAB18, BioID2(Gly40Ser)-SEC22A and mEmerald-SEC22A were generated following PCR amplification from template and subcloning into an intermediate pCR-Blunt II-TOPO vector using a Zero Blunt TOPO PCR Cloning Kit (ThermoFisher Scientific, Waltham, MA) according to manufacturer’s instructions. Fragments were excised from intermediate vectors and then subcloned into target vectors using restriction-ligation, as above. A construct encoding mCherry-ER was obtained from Addgene, and a construct encoding TMCO4-EGFP was synthesised and cloned by GeneWiz (Leipzig, Germany). Generation of recombinant pX461 and pX462 plasmids for CRISPR gene-editing, and recombinant pCMV vectors for preparation of stable CHO cell lines are described below. Generation of recombinant pcDNA5 FRT/TO FLAG-BirA(Arg118Gly) vectors for preparation of stable T-Rex-293 cell lines is described in supplementary methods. Details of other plasmids, PCR templates, primers and target vectors are listed in Table S6.

### Antibodies and reagents

A custom polyclonal antibody to RAB18 generated by Eurogentec (Southampton, UK) has been described previously (10). An antibody to RAB3GAP1 was obtained from Bethyl Labs (Montgomery, TX), an antibody to GFP was obtained from Takara Bio (Saint-Germain-en-Laye, France), an antibody to β-Tubulin was obtained from Abcam (Cambridge, UK) and an antibody to β-Actin was obtained from ThermoFisher. Antibodies to hemagglutinin (HA), RAB3GAP2 and TBC1D20 were obtained from Merck (Gillingham, UK). Antibodies to ZW10, STX18, SPG20, RINT1, REEP4, BNIP1, C2CD2, TRIM13, WFS1, INPP5B, OSBPL2 and NBAS were obtained from Proteintech (Manchester, UK). Antibody catalogue numbers and the dilutions used in this study are listed in Table S6.

### Cell culture

HeLa, T-REx-293, IHH cells and human fibroblasts were maintained in DMEM media, RPE1 cells in DMEM/F12 media and CHO cells in alpha-MEM media (ThermoFisher). In each case, media was supplemented with 10% foetal calf serum (FCS) and 1% penicillin-streptomycin (PS). Cells were maintained at 37°C and 5% CO_2_.

### Generation of clonal ‘knockout’ HeLa and RPE1 cell lines

CRISPR/Cas9 gene-editing was carried out essentially as described in Ran et al., 2013 (57). Guide RNA (gRNA) sequences are shown in Table S6. A list of the clonal cell lines generated for this study, together with the loss-of-function variants they carry is shown in Figure S1A. Western blot validation is shown in Figure S1B-E. Briefly, for each targeted exon, pairs of gRNA sequences were selected using the online CRISPR design tool (http://crispr.mit.edu/). Oligonucleotide pairs incorporating these sequences (Sigma) were annealed (at 50mM ea.) in 10mM Tris pH8, 50mM NaCl and 1mM EDTA by incubation at 95⁰C for 10 minutes followed by cooling to room temperature. Annealed oligonucleotides were diluted and ligated into BbsI-digested pX461 and pX462 plasmids (Addgene) using HC T4 Ligase and rapid ligation buffer (Promega). Sequences of all recombinant plasmids were verified by direct sequencing. Pairs of plasmids were contransfected into cells using Lipofectamine 2000 reagent according to manufacturer’s instructions. Cells were selected for puromycin resistance (conferred by pX462) using 24 hours puromycin-treatment. Following 12 hours recovery, they were selected for GFP fluorescence (conferred by pX461) and cloned using FACSAria2 SORP, Influx or FACSMelody instruments (BD, Wokingham, UK). After sufficient growth, clones were analysed by PCR of the targeted exons (Primers are listed in Table S6). In order to sequence individual gene-edited alleles, PCR products from each clone were first cloned into ZeroBlunt TOPO vector (ThermoFisher) and then subjected to colony PCR. These PCR products were then analysed by direct sequencing. Sequencing data was assessed using BioEdit software (http://www.mbio.ncsu.edu/BioEdit/bioedit.html).

### BirA*/BioID proximity labelling (HeLa cells)

Proximity-labelling in HeLa cells was carried out largely as described by Roux et al. (15), but with minor modifications. HeLa cells were grown to 80% confluence in T75 flasks and then each flask was transfected with 1-1.5µg of the BirA*-RAB18 construct or 1µg of the BioID2(Gly40Ser)-SEC22A construct using Lipofectamine 2000 reagent in Optimem serum-free medium (ThermoFisher) for 4 hours, according to manufacturer’s instructions. 24 hours post-transfection, media was replaced with fresh media containing 50µM Biotin (Merck) and the cells were incubated for a further 24 or 6 hours (for BirA*-RAB18 and BioID2(Gly40Ser)-SEC22A experiments respectively). Cells were then trypsinised and washed twice in PBS before pellets were transferred to 2ml microcentrifuge tubes and snap-frozen. For each pellet, lysis was carried out in 420µl of a buffer containing 0.2% SDS, 6% Triton-X-100, 500mM NaCl, 1mM DTT, EDTA-free protease-inhibitor solution (Expedeon, Cambridge, UK), 50mM Tris pH7.4. Lysates were sonicated for 10 minutes using a Bioruptor device together with protein extraction beads (Diagenode, Denville, NJ). Each lysate was diluted with 1080µl 50mM Tris pH7.4, and they were then clarified by centrifugation at 20 000xg for 30 minutes at 4⁰C. Affinity purification of biotinylated proteins was carried out by incubation of clarified lysates with streptavidin-coated magnetic Dynabeads (ThermoFisher) for 24 hours at 4⁰C. Note that a mixture of Dynabeads - MyOne C1, MyOne T1, M270 and M280 – was used to overcome a problem with bead-clumping observed when MyOne C1 beads were used alone. Successive washes were carried out at room temperature with 2% SDS, a buffer containing 1% Triton-X-100, 1mM EDTA, 500mM NaCl, 50mM HEPES pH7.5, a buffer containing 0.5% NP40, 1mM EDTA, 250mM LiCl, 10mM Tris pH7.4, 50mM Tris pH7.4, and 50mM ammonium bicarbonate.

### Mass spectrometry

Washed beads from BioID experiments with HeLa cells were subjected to limited proteolysis by trypsin (0.3 µg) at 27°C for 6.5hours in 2mM urea, 1mM DTT, 75mM Tris, pH=8.5, then supernatants were incubated overnight at 37°C. Samples were alkylated with 50mM iodoacetamide (IAA) in the dark for 20minutes, then acidified by addition of 8µl 10% trifluoroacetic acid (TFA). Peptides were generated using trypsin. Trypsin cleaves on the C-terminal side of lysine and arginine residues unless the C-terminal residue is proline. Hydrolysis is slower where the C-terminal residue is acidic. Peptides were loaded on to activated (methanol), equilibrated (0.1% TFA) C18 stage tips before being washed with 0.1% TFA and eluted with 0.1% TFA/80 acetonitrile. The organic was dried off, 0.1% TFA added to 15 µl and 5 µl injected onto LC-MS. Peptides were separated on an Ultimate nano HPLC instrument (ThermoFisher), and analysed on either an Orbitrap Lumos or a Q Exactive Plus instrument (ThermoFisher).

Three sets of replicate samples were used for the BioID-RAB18 experiment with HeLa cells (Figure 1, Table S1). Two different wild-type clones and two different null-genotypes of each of the RAB3GAP1-, RAB3GAP2- and TRAPPC9-null cells were used (see Figure S1). Three sets of replicate samples were used in the BioID2(Gly40Ser)-SEC22A experiment (Figure 3D, Table S5), though preparation of one ‘RAB3GAP1-null’ replicate failed. In each experiment, each set of samples was prepared independently and so these can be considered biological replicates.

### Analysis

After data-dependent acquisition of HCD fragmentation spectra, data were analysed using MaxQuant (version 2.2.0.0 for the BioID-RAB18 experiment and version 1.6.7.0 for the BioID2(Gly40Ser)-SEC22A experiment). For the BioID-RAB18 experiment, the Uniprot Human 2022_05 database with 20594 entries was searched. For the BioID2(Gly40Ser)-SEC22A experiment, the Uniprot Human 2019_07 database with 20667 entries was searched. 2 missed/non-specific cleavages were permitted. Fixed modification by carbamidomethylation of cysteine residues was considered. Variable modification by oxidation of methionine residues and N-terminal acetylation were considered. Mass error was set at 20 ppm for the first search tolerance and 4.5 ppm main search tolerance. Thresholds for accepting individual spectra were set at p<0.05. Single-peptide identifications of proteins were used in analysis of the BioID-RAB18 experiment with single peptide identifications made ‘by modification site only’ excluded. %FDR for these single-peptide identifications, and that for the combined dataset, was estimated at <5% using the decoy search method. Additional parameters and gradients used for separation are provided in Table S6. Annotated spectra for single-peptide identifications are provided in the ‘single_peptide_identifications’ document in the supporting information.

Quantification data were produced with MaxLFQ [19]. For the BioID-RAB18 experiment, data were first processed to remove any protein identified in samples from control (untransfected, biotin-treated) samples at high levels (>25% wild-type LFQ value) in any replicate from all replicates. Next, proteins identified in only one replicate sample-set were removed. For each sample set, LFQ values were normalized according to the quantity of RAB18 detected in each sample. GEF-null:wild-type ratios for each protein were calculated for each replicate sample set and then their means calculated for the experiment (Table S1, columns ‘Mean RAB3GAP Ratio’, ‘Mean TRAPII Ratio’). The GEF-null:wild-type <0.5 criterion for selection of putative effectors (Fig. 1A,D, Table 1) is an arbitrary cutoff rather than a measure of statistical validity. A similar approach was taken to analyses of BioID2-SEC22A data (Table S5), except that ‘Mean RAB3GAP Ratio’ and ‘Mean RAB18 Ratio’ were calculated for each protein.

### Cell labelling

In order to distinguish cells of different genotypes within the same well/on the same coverslip, CellTrace Violet and CellTrace Far Red reagents (ThermoFisher) were used to label cells before they were seeded. Cells of different genotypes were first trypsinised and washed with PBS separately. They were then stained in suspension by incubation with either 1µM CellTrace Violet or 200nM CellTrace Far Red for 20 minutes at 37°C. Remaining dye was removed by addition of a ten-fold excess of full media, incubation for a further 5 minutes, and then by centrifugation and resuspension of the resulting pellets in fresh media. Differently-labelled cells were combined prior to seeding.

### Immunofluorescence microscopy

Cells were seeded in 96-well glass-bottom plates (PerkinElmer, Waltham, MA) coated with Matrigel (Corning, Amsterdam, Netherlands) according to manufacturer’s instructions, and allowed to adhere for 48 hours prior to fixation. In lipid-loading experiments, cells were treated with 200μM oleic acid complexed to albumin (Merck) and 1μg/ml BODIPY-558/568-C12 (ThermoFisher) for 15 hours prior to fixation. Cells were fixed using a solution of 3% deionised Glyoxal, 20% EtOH, 0.75% acetic acid, pH=5 (58), for 20 minutes at room temperature. They were then washed with PBS containing 0.9mM CaCl_2_ and 0.5mM MgCl_2_ and blocked with a sterile-filtered buffer containing 1% Milk, 2% donkey serum (Merck), 0.05% Triton-X-100 (Merck), 0.9mM CaCl_2_ and 0.5mM MgCl_2_ in PBS pH=7.4 for at least 1 hour prior to incubation with primary antibody. Primary antibodies were added in blocking buffer without Triton-X-100, and plates were incubated overnight at 4⁰C. Antibody dilutions are listed in Table S6. Following washing in PBS, cells were incubated with 1:2000 Alexa 488-conjugated secondary antibody (ThermoFisher) in blocking buffer at room temperature for 1-2 hours. Following further washing in PBS, cells were imaged using an Operetta High Content Imaging System (PerkinElmer) equipped with Harmony software. In comparative fluorescence quantitation experiments, at least 18 frames – each containing >5 wild-type and >5 mutant cells – were analysed per genotype. ImageJ software was used to produce regions of interest (ROIs) corresponding to each cell using thresholding tools and images from the 405nm and 645nm channels. Median 490nm fluorescence intensity was measured for each cell and mutant fluorescence intensity (as %wild-type) was calculated for each frame and combined for each genotype.

### Confocal microscopy – Live cell imaging

HeLa or RPE1 cells were seeded on glass-bottom dishes (World Precision Instruments, Hitchin, UK) coated with Matrigel (Corning) and allowed to adhere for 24 hours prior to transfection. Transfections and cotransfections were carried out with 0.5µg of each of the indicated constructs using Lipofectamine 2000 reagent in Optimem serum-free medium for 4 hours, according to manufacturer’s instructions. Media were replaced and cells were allowed to recover for at least 18 hours prior to imaging. Imaging was carried out on a Nikon A1R confocal microscope equipped with the Nikon Perfect Focus System using a 60× oil immersion objective with a 1.4 numerical aperture. The pinhole was set to airy1. CellTrace Violet was excited using a 403.5nm laser, and emitted light was collected at 425–475nm. EGFP and mEmerald were excited using a 488 nm laser, and emitted light was collected at 500–550 nm. mCherry was excited using a 561.3 nm laser, and emitted light was collected at 570–620 nm. CellTrace Far Red was excited using a 638nm laser, and emitted light was collected at 663-738nm.

### Immunoprecipitation

HeLa cells were seeded onto 10cm dishes or 6-well plates and allowed to adhere for 24 hours prior to transfection. Transfections and cotransfections were carried out with 0.5µg of each of the indicated constructs using Lipofectamine 2000 reagent in Optimem serum-free medium for 4 hours, according to manufacturer’s instructions. 24 hours post-transfection cells were trypsinised, washed with PBS, then lysed in a buffer containing 150mM NaCl, 0.5% Triton-X-100 and EDTA-free protease-inhibitor solution (Expedeon), 10mM Tris, pH=7.4. Lysates were clarified by centrifugation, input samples taken, and the remaining supernatants then added to 4µg rabbit anti-HA antibody (Merck). After 30 minutes incubation at 4⁰C on a rotator, 100µl washed protein G-coupled Dynabeads (ThermoFisher) were added and samples were incubated for a further 1 hour. The Dynabeads were washed x3 with buffer containing 150mM NaCl, 0.1% Triton-X-100, 10mM Tris, pH=7.4, then combined with a reducing loading buffer and subjected to SDS–PAGE.

### Generation of stable CHO cell lines

A PCR product encoding mouse RAB18 was subcloned into an intermediate TOPO vector using a TOPO PCR Cloning Kit (ThermoFisher) according to manufacturer’s instructions. The RAB18 fragment was then excised and subcloned into the pCMV vector. PCR-based site-directed mutagenesis using a GeneArt kit (ThermoFisher) was then used to generate pCMV-RAB18(Gln67Leu) and pCMV-RAB18(Ser22Asn) constructs. CHO cells were transfected using Lipofectamine 2000 reagent (ThermoFisher) and cells stably-expressing each construct were selected-for with blasticidin. Under continued selection, clonal cell-lines were grown from single cells and then RAB18 protein expression was assessed. Cell lines comparably expressing RAB18 constructs at levels 2.5-5x higher than those wild-type cells were used in subsequent experiments.

### Lipid loading experiments

For LD number and diameter measurements, IHH cells were seeded onto glass coverslips. siRNA transfections were carried out using FuGene reagent (Promega) according to manufacturer’s instructions. siRNAs targeting ZW10 and NBAS were obtained from IDT, Coralville, IA; siRNA targeting SEC22A was obtained from Horizon Discovery, Cambridge, UK. 48 hours following transfection, cells were treated with 200nM BSA conjugated oleate for 24 hours. Coverslips were washed, fixed with 3% paraformaldehyde and stained with 1µg/mL BODIPY and 300nM DAPI. Fluorescence images were captured on a Zeiss LSM 780 confocal microscope equipped with a 100x objective. Images were analysed using ImageJ software. Data are representative of three independent experiments.

For cholesterol storage and efflux experiments with [^14^C]-oleate, CHO cell lines (described above) were seeded onto 12-well plates and then grown to 60-75% confluence in Alpha media supplemented with 10% LPDS. Cells were grown in the presence of 10% LPDS for at least 24 hours prior to the addition of oleate. 1 µCi/ml [^14^C]-oleate (Perkin Elmer) was added in the presence of 10% LPDS or 10% FBS for 24 hours. Cells were then washed and incubated with 50µg/ml HDL for 0, 4 or 8 hours. Cellular lipids were extracted with hexane. Lipids were then dried-down and separated by thin layer chromatography (TLC) in a hexane:diethyl ether:acetic acid (80:20:2) solvent system. TLC plates were obtained from Analtech, Newark, NJ. Bands corresponding to cholesteryl ester (CE) were scraped from the TLC plate, and radioactivity was determined by scintillation counting in a Beckman Coulter LS6500 Scintillation Counter using BetaMax ES Liquid Scintillation Cocktail (ThermoFisher). Three independent experiments were carried out, each with four replicates of each condition. Data from a representative experiment are shown.

For cholesterol efflux experiments with [^3^H]-cholesterol, CHO cells were seeded onto 12-well plates and then grown to 60% confluence in Alpha media supplemented with 10% FBS. 5 µCi/ml [^3^H]-cholesterol (PerkinElmer) was added in the presence of 10% FBS. After 3x PBS washes, cells were incubated with serum-free media containing 25µg/ml of human apolipoprotein A-I (ApoA-I) for 5 hours. ApoA-I was a kind gift of Dr. Paul Weers (California State University, Long Beach). Radioactivity in aliquots of media were determined by scintillation counting in a Beckman Coulter LS6500 Scintillation Counter using LSC Cocktail (PerkinElmer). Cell lysates were produced by addition of 0.1N NaOH for 1 hour, and their radioactivity was determined as above. Cholesterol efflux was calculated as an average (+/-SD) of the % cholesterol efflux (as a ratio of the media cpm/(media + cellular cpm) x 100%).

For the cholesterol biosynthesis experiments, HeLa cells were seeded onto 12-well plates and then grown to 80% confluence in DMEM supplemented with 10% LPDS for 24 hours. Following incubation with 5µCi/well [3H]-mevalonate or 10µCi/well [3H]-acetate for 24 hours, TLC was used to separate free cholesterol and radioactivity was quantified by scintillation counting as above.

### Sterol analysis (HeLa cells)

HeLa cells were grown to 80% confluence in T75 flasks, washed twice in PBS and then grown for a further 48 hours in DMEM supplemented with 10% LPDS. They were then trypsinised and washed twice in PBS before pellets were transferred to microcentrifuge tubes and snap-frozen. Pellets were resuspended in 200µl deionised water, sonicated for 20 seconds using an ultrasonic processor (Sonics & Materials Inc., CT, USA), then placed on ice. 750 µl of isopropanol containing 4 µmol/L 5α-cholestane as an internal standard was added to each sample, and then each was sonicated for a further 10 seconds. Lysates were transferred to 7ml glass vials and mixed with 250 µl tetramethylammonium hydroxide for alkaline saponification at 80°C for 15 min, then cooled down for 10 minutes at room temperature. Sterols were extracted by addition of 500 µl tetrachloroethylene/methyl butyrate (1:3) and 2 ml deionised water, then thorough mixing. Samples were centrifuged for 10 minutes at 3000 rpm, and the organic phase containing the sterols was transferred to 300 µl GC vials. Extracts were dried under a stream of nitrogen, then sterols were silylated with 50 µL Tri-Sil HTP (HDMS:TMCS:Pyridine) Reagent (ThermoFisher) at 60°C for 1 hour.

Chromatography separation was performed on an Agilent gas chromatography-mass spectrometry (GC-MS) system (6890A GC and 5973 MS) (Agilent Technologies, Inc., CA, USA) with an HP-1MS capillary column (30 m length. x 250 µm diameter x 0.25 µm film thickness). The GC temperature gradient was as follows: Initial temperature of 120°C increased to 200°C at a rate of 20°C/min, then increased to 300°C at a rate of 2°C/min with a 15 minute solvent delay. Injection was at 250°C in splitless mode with ultrapurified helium as the carrier gas and the transfer line was 280°C. The mass spectra were acquired by electron impact at 70 eV using selected ion monitoring as follows: Lathosterol-TMS, cholesterol-TMS, and cholest8(9)-enol-TMS: m/z 458; 5α-cholestane and desmosterol-TMS: m/z 372; Lanosterol-TMS: m/z 393; and 7-dehydrocholesterol-TMS: m/z 325. The data were analysed using MassHunter Workstation Quantitative Analysis Software (Agilent Technologies, Inc.) and OriginPro 2017 (OriginLab Corp, MA, USA).

### GC-FID and GC-MS-SIM Sterol and stanol analysis (Fibroblasts)

Gas chromatographic separation and detection of cholesterol and 5α-cholestane (internal standard, ISTD) was performed on a DB-XLB 30m x 0.25 mm i.d. x 0.25 µm film thickness (J&W Scientific Alltech, Folsom, CA, U.S.A.) in an Hewlett-Packard (HP) 6890 Series GC-system (Agilent Technologies, Palo Alto, CA, U.S.A), equipped with an flame ionization detector (FID).

Non-cholesterol sterols such as the cholesterol precursors lanosterol, 24.25-dihydrolanosterol, desmosterol, lathosterol and the cholesterol metabolite 5α-cholestanol together with epicoprostanol (ISTD) were separated on another DB-XLB column (30m x 0.25 mm i.d. x 0.25 µm film thickness, J&W Scientific Alltech, Folsom, CA, U.S.A.) in a HP 6890N Network GC system (Agilent Technologies, Waldbronn, Germany) connected with a direct capillary inlet system to a quadrupole mass selective detector HP5975B inert MSD (Agilent Technologies, Waldbronn, Germany). Both GC systems were equipped with HP 7687 series auto samplers and HP 7683 series injectors (Agilent Technologies, Waldbronn, Germany).

To determine the concentrations of cholesterol, non-cholesterol precursor sterols and 5a-cholestanol, 50 μg 5α-cholestane (Serva, Heidelberg, Germany) (50 μl from a stock solution of 5α-cholestane in cyclohexane [Merck KGaA, Darmstadt, Germany]; 1 mg/ml), and one μg epicoprostanol (Sigma, Deisenhofen, Germany) (10 μl from a stock solution epicoprostanol in cyclohexane; 100 μg/ml) were added as internal standards to 100 µl of an chloroform/methanol cell extract (5 ml chloroform/methanol, [2:1, v/v] per 10 mg dried cells). The cell pellet were dried for 12 hours at room temperature in a Savant DNA120, Speed Vac® Concentrator system (Thermo Fisher Scientific, Kandel, Germany).

After saponification with 2 mL 1M 95% ethanolic sodium hydroxide solution (Merck KGaA, Darmstadt, Germany) at 60°C for one hour, the free sterols were extracted three times with 3 mL cyclohexane each from dried tissues (speedvac). The organic solvent was evaporated by a gentle stream of nitrogen at 60°C on a heating block. The residue was dissolved in 80 µL n-decane (Merck KGaA, Darmstadt, Germany). An aliquot of 40 µL was incubated (1h at 70°C on a heating block) by addition of 20 µL of the trimethylsilylating (TMSi) reagent (chlortrimethylsilane [Merck KGaA, Darmstadt, Germany]/1.1.1.3.3.3-Hexamethyldisilasane [Sigma Aldrich, Co., St. Louis, MO, U.S.A]/pyridine [Merck KGaA, Darmstadt, Germany], 9:3:1) in a GC vial for GC-MS-SIM non-cholesterol analysis. Another aliquot of 40µL was incubated by addition of 40 µL of the TMSi-reagent and dilution with 300 µL n-decane in a GC vial for GC-FID cholesterol analysis.

An aliquot of 2 µl was injected by automated injection in a splitless mode using helium (1mL/min) as carrier gas for GC-MS-SIM and hydrogen (1ml/min) for GC-FID analysis at an injection temperature of 280°C. The temperature program for GC was as follows: 150°C for three minutes, followed by 20°C/min up to 290°C keeping for 34 minutes. For MSD electron impact ionization was applied with 70 eV. SIM was performed by cycling the quadrupole mass filter between different *m/z* at a rate of 3.7 cycles/sec. Non-cholesterol sterols were monitored as their TMSi-, the oxysterols as their di-TMSi-derivatives using the following masses: epicoprostanol m/z 370, lathosterol at m/z 458, desmosterol at m/z 441, lanosterol at m/z 393, 24.25-dihydrolanosterol at m/z 395 (M^+^-90-15, M^+^-OTMSi-CH_3_), and 5α-cholestanol at m/z 306. Peak integration was performed manually. Cholesterol was directly quantified by multiplying the ratios of the area under the curve of cholesterol to 5α-cholestane by 50 µg (ISTD amount). Non-cholesterol sterols and 5α-cholestanol were quantified from the ratios of the areas under the curve of the respective non-cholesterol sterols and 5α-cholestanol after SIM analyses against epicoprostanol using standard curves for the listed sterols/stanol. Identity of all sterols and 5α-cholestanol was proven by comparison with the full-scan mass spectra of authentic compounds. Additional qualifiers (characteristic fragment ions) were used for structural identification (m/z values not shown).

### Western blotting

Cell lysates were made with a buffer containing 150mM NaCl, 0.5% Triton-X-100 and EDTA-free protease-inhibitor solution (Expedeon), 50mM Tris, pH=7.4. Cell lysates and input samples from BioID and immunoprecipitation experiments were combined 1:1 with a 2x reducing loading buffer; a reducing loading buffer containing 10mM EDTA was added directly to Dynabead samples. SDS–PAGE and Western blotting were carried out according to standard methods.

## DATA AVAILABILITY

The mass spectrometry proteomics data have been deposited to the ProteomeXchange Consortium via the PRIDE (59) partner repository with the dataset identifiers PXD016631, PXD016336, PXD016326, PXD016233 and PXD016404.

## SUPPORTING INFORMATION

This article contains supporting information.

## Supporting information

single peptide identifications

## ACKNOWLEDGEMENTS

We thank the Warburg Micro syndrome children and their families. We thank Professor C. A. Johnson, Dr J. A. Poulter and Dr I Tsagakis for a critical reading of the manuscript.

## FUNDING AND ADDITIONAL INFORMATION

MH was supported by the University of Leeds and by a Programme Grant from the Newlife Foundation for Disabled Children (Grant Reference Number: 17-18/23). AvK and JW were supported by the Wellcome Trust (Multiuser Equipment Grant, 208402/Z/17/Z).

## CONFLICT OF INTEREST

The authors declare that they have no conflicts of interest with the contents of this article.

## Notes

### Competing Interest Statement

The authors have declared no competing interest.

### Summary of Updates

This version contains updated analysis of the original BioID data, and additional sterol-profiling and immunoprecipitation data.

