## Supporting Information for "Comparative proximity biotinylation implicates RAB18 in sterol mobilization and biosynthesis"

A

| ID# | Clone name | Cell type | Targeted gene | Reference sequence | Targeted exon | gRNA sequences | cDNA alleles | protein alleles | Further validation |
|---|---|---|---|---|---|---|---|---|---|
| 1 | RAB18_x7c14 | HeLa | *RAB18* | NM_021252.5 | 7 | ggcacattgtacaccatcac<br>gcattcagacccctggactgt | c.457_512del<br>c.455_523delins49<br>c.446-12_532delins2 | p.Lys153Aspfs*7<br>p.Ala152Glufs*20 | Fig. S1B |
| 2 | RAB18_x7c21 | HeLa | *RAB18* | NM_021252.5 | 7 | ggcacattgtacaccatcac<br>gcattcagacccctggactgt | c.456_523del<br>c.455_527del<br>c.472_490del | p.Lys153Valfs*3<br>p.Ala152Glyfs*44<br>p.Val158Asnfs*56 | Fig. S1B |
| 3 | RAB18_x7c31 | RPE1 | *RAB18* | NM_021252.5 | 7 | ggcacattgtacaccatcac<br>gcattcagacccctggactgt | c.477_485delins5<br>c.477_521delins28 | p.Cys160Glnfs*59<br>p.Glu164Lysfs*9 | Fig. S1B |
| 4 | RAB18_x7c32 | RPE1 | *RAB18* | NM_021252.5 | 7 | ggcacattgtacaccatcac<br>gcattcagacccctggactgt | c.477_485delins5<br>c.477_521delins28 | p.Cys160Glnfs*59<br>p.Glu164Lysfs*9 | Fig. S1B |
| 5 | RAB3GAP1_x15c3 | HeLa | *RAB3GAP1* | NM_012233.3 | 15 | gccactcctttcaaccctcca<br>gtcttgaaatgcgtttccgat | c.1417_1423del<br>c.1416_1432del | p.Gly473Lysfs*22<br>p.Gly473Thrfs*8 | Fig. S1C |
| 6 | RAB3GAP1x15c5 | HeLa | *RAB3GAP1* | NM_012233.3 | 15 | gccactcctttcaaccctcca<br>gtcttgaaatgcgtttccgat | c.1471_1472ins148<br>c.1453_1499+16del | p.Arg491Thrfs*8 | Fig. S1C |
| 7 | RAB3GAP1_x15c23 | RPE1 | *RAB3GAP1* | NM_012233.3 | 15 | gccactcctttcaaccctcca<br>gtcttgaaatgcgtttccgat | c.1429_1448del<br>c.1444_1445ins79 | p.Val477Ilefs*3<br>p.Gln482Leu*6 | Fig. S1C |
| 8 | RAB3GAP1_x15c24 | RPE1 | *RAB3GAP1* | NM_012233.3 | 15 | gccactcctttcaaccctcca<br>gtcttgaaatgcgtttccgat | c.1419_1432delins28<br>c.1452_1453ins26 | p.Leu474Valfs*28<br>p.Val485Profs*21 | |
| 9 | RAB3GAP2_x14c2 | HeLa | *RAB3GAP2* | NM_012414.4 | 14 | gaggaattgagctactcgact<br>gtgatctatgcgccaagaag | c.1385_1424del<br>c.1382_1397delins3<br>c.1381_1382insC | p.Ala462Glufs*11<br>p.Val461Glufs*10<br>p.Val461Alafs*24 | Fig. S1D |
| 10 | RAB3GAP2_x14c26 | RPE1 | *RAB3GAP2* | NM_012414.4 | 14 | gaggaattgagctactcgact<br>gtgatctatgcgccaagaag | c.1409_1410ins75<br>c.1408_1409ins16 | p.Arg471*<br>p.Pro470Ilefs*3 | Fig. S1D |
| 11 | RAB3GAP2_x20c12 | HeLa | *RAB3GAP2* | NM_012414.4 | 20 | gtatatttctctagtaatgcc<br>gcgattttctgatgataaaga | c.2056_2116del<br>c.2109_2110ins133<br>c.2058_2059ins6; c.2087_2097del | p.Leu686Valfs*4<br>p.Lys704Tyrfs*1<br>p.Leu686_Glu687insLeuLeu; p.Thr696Ilefs*2 | Fig. S1D |
| 12 | RAB3GAP2_x20c32 | RPE1 | *RAB3GAP2* | NM_012414.4 | 20 | gtatatttctctagtaatgcc<br>gcgattttctgatgataaaga | c.2100_2101ins88<br>c.2100_2101ins49 | p.Ser701Lysfs*7<br>p.Ser701Ilefs*46 | Fig. S1D |
| 13 | SPG20_x3c19 | HeLa | *SPART* | NM_015087.5 | 3 | gaatgtctgaccttcggctcc<br>gaagagctctctatcatcctc | c.962_977del<br>c.957_1006delins48 | p.Glu321Glyfs*1<br>p.Asp319Glufs*27 | Fig. 2C |
| 14 | TBC1D20_5H2 | HeLa | *TBC1D20* | NM_144628.4 | 5 | gtgcttggtgttgtccattgt<br>gtctgatgcccatcattgacc | c.558_586del<br>c.551_594del | p.Lys186Asnfs*1<br>p.Asn184Serfs*6 | Fig. S1E |
| 15 | TBC1D20_x5c31 | RPE1 | *TBC1D20* | NM_144628.4 | 5 | gtgcttggtgttgtccattgt<br>gtctgatgcccatcattgacc | c.547_583dup<br>c.567_568ins39 | p.Ile195Lysfs*13<br>p.Asn190_Glu197insProThrGluThrSerCysSer* | |
| 16 | TBC1D20_x5c33 | RPE1 | *TBC1D20* | NM_144628.4 | 5 | gtgcttggtgttgtccattgt<br>gtctgatgcccatcattgacc | c.577_578ins85<br>c.583_584ins83 | p.Met193Thrfs*6<br>p.Ile195Thrfs*5 | Fig. S1E |
| 17 | TBC1D20_7H5 | HeLa | *TBC1D20* | NM_144628.4 | 7 | gctgctgatcagtgtctcata<br>gcccatccgaacttgctcggg | c.890_929del<br>c.860_923del | p.Asp297Glyfs*49<br>p.Pro287Leufs*51 | Fig. S1E |
| 18 | TRAPPC9_x13c1 | HeLa | *TRAPPC9* | NM_001160372.4 | 13 | ggagttcgagtctctccctg<br>gcgacgcagcatcgtaagcc | c.1872_1897del<br>c.1855-46_1865del | p.Val625Alafs*5 | |
| 19 | TRAPPC9_x13c3 | RPE1 | *TRAPPC9* | NM_001160372.4 | 13 | ggagttcgagtctctccctg<br>gcgacgcagcatcgtaagcc | c.1855-25_1880del<br>c.1855-13_1896del | | |
| 20 | TRAPPC9_x14c6 | HeLa | *TRAPPC9* | NM_001160372.4 | 14 | gcgcgggaatgacttccactg<br>gtgcccaggtagtgtgcacg | c.2108_2114+4delins28<br>c.2058_2114+24del | | |

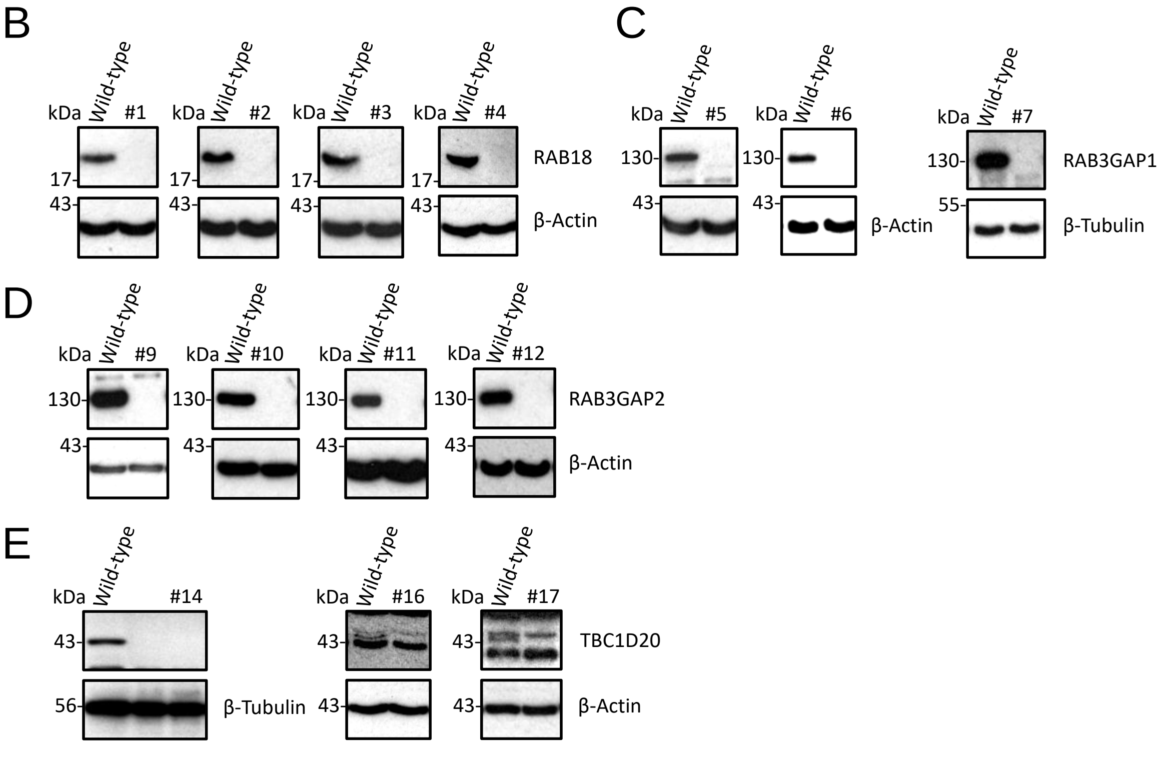


**Figure S1. Genetically-validated gene-edited clonal cell lines used in this study.** (A) Table to show clonal cell lines generated by transient expression of

Cas9n together with the indicated guide RNAs. Mutant alleles of the targeted genes were sequenced following their cloning into ZeroBlunt TOPO vector. (B) Western blotting for RAB18 shows that RAB18 is undetectable in RAB18-null cell lines. (C) Western blotting for RAB3GAP1 shows that RAB3GAP1 is undetectable in RAB3GAP1-null cell lines. (D) Western blotting for RAB3GAP2 shows that RAB3GAP2 is undetectable in RAB3GAP2-null cell lines. (E) Western blotting for TBC1D20 shows that TBC1D20 is undetectable in TBC1D20-null cell lines. Numbering in S1A corresponds to lane numbering in S1B-E. Loading controls are β-Actin or β-Tubulin. Note that different staining patterns for anti-TBC1D20 antibody in S1E are the result of antibody batch-variability.

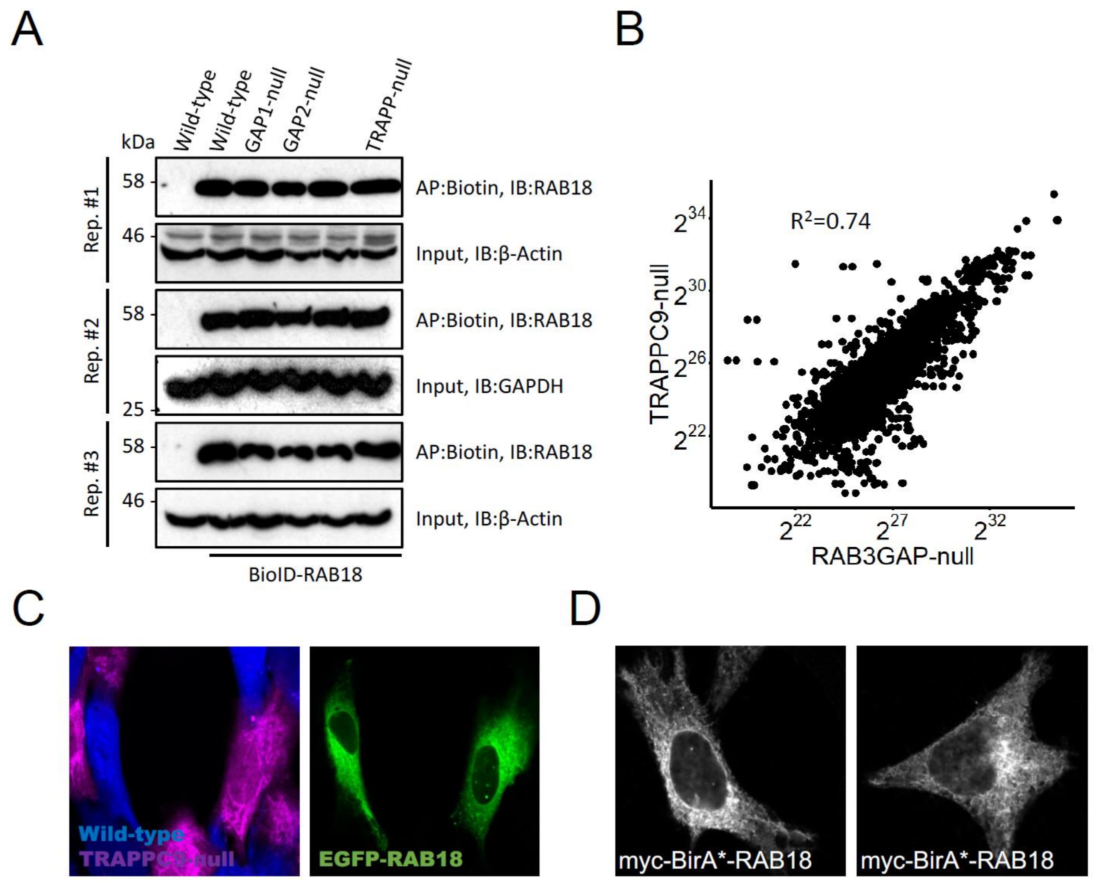


**Figure S2. Controls for HeLa BioID experiments.** (A) Western blotting to show comparable levels of BirA*-RAB18 in BioID samples from HeLa cells of wild-type and different mutant genotypes. Loading controls are β-Actin or GAPDH. (B) Plots to show correlations between non-zero LFQP intensities of individual proteins identified in samples purified from TRAPPC9-null cells and those purified from RAB3GAP1/2-null cells. (C) Confocal micrograph to show comparable localization of exogenous EGFP-RAB18 (Green) in wild-type and TRAPPC9-null HeLa cells. Cells were labelled with CellTrace-Violet and CellTrace-Far Red reagents respectively (magenta and blue channels). (D) Confocal micrographs showing the localization of BirA*-RAB18 in HeLa cells. Transfected HeLa cells were fixed with 3% deionized glyoxal, then permeabilized and stained with a mouse monoclonal anti-myc antibody and an Alexa-488-conjugated anti-mouse secondary antibody.

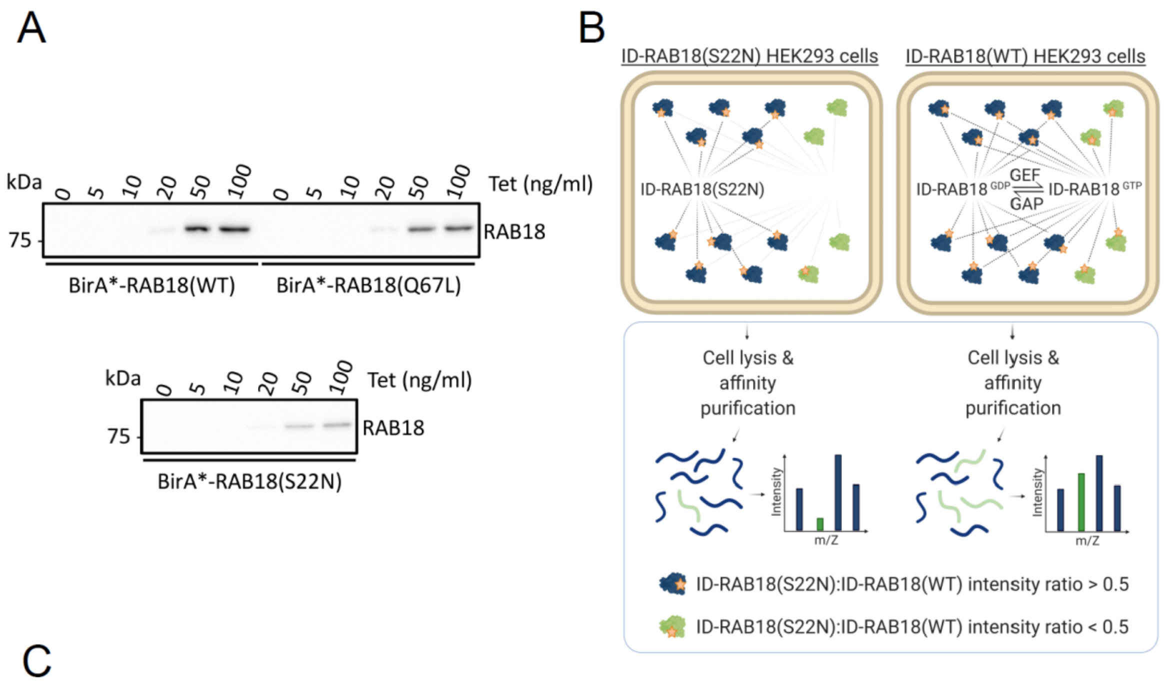


C

| Protein | Ratio | Orthologue PPI (Gillingham et al., 2014) | Additional evidence | |
|---|---|---|---|---|
| NBAS | 0.00 | rod | Xu et al., 2018; Gillingham et al., 2019 | **RAB3GAP-dependent association in HeLa cells** |
| SCFD2 | 0.00 | Slh | Gillingham et al., 2019 | |
| ZW10 | 0.00 | mit(1)15 | Xu et al., 2018; Gillingham et al., 2019 | |
| CAMSAP1 | 0.04 | | | |
| RINT1 | 0.41 | CG8605 | Xu et al., 2018 | |
| BNIP1 | 0.65 | | Xu et al., 2018 | |
| RAB3GAP2 | 1.58 | rab3-GAP | Gerondopoulos et al., 2014 | |
| RAB3GAP1 | 1.59 | CG31935 | Gerondopoulos et al., 2014 | |
| ARPIN | 0.00 | | | **TRAPPII-dependent association in HeLa cells** |
| SEC23IP | 0.00 | | | |
| TBC1D5 | 0.00 | CG8449 | Gillingham et al., 2019 | |
| ATP6AP2 | 0.29 | | | |
| GORASP2 | 0.36 | | | |
| GIGYF2 | 0.41 | | | |
| EFHD2 | 0.50 | | | |
| BET1 | 0.54 | | | |
| RABL3 | 0.58 | | | |
| CEP170 | 0.63 | | | |
| ARFGAP1 | 0.65 | | | |
| EPS15L1 | 0.65 | | | |
| MKL2 | 0.77 | | | |
| GOLGA5 | 0.79 | | | |
| CLCC1 | 0.89 | | | |
| ABCD3 | 0.93 | | | |
| SEC22B | 0.96 | | | |
| AKAP1 | 1.47 | | | |
| HDLBP | 1.50 | | | |
| DNML1 | 1.59 | | | |

**Figure S3. Nucleotide-binding-dependent RAB18-associations in HEK293 cells.** (A) Tetracycline-induced expression of BirA*-RAB18 constructs in stable HEK293 cell lines. PCR products encoding mouse RAB18, RAB18(Gln67Leu) and RAB18(Ser22Asn) were subcloned into a pcDNA5 FRT/TO FLAG-BirA(Arg118Gly) vector. Recombinant vectors were used together with pOG44 in cotrasfections of T-

REx-293 cells. Stable cell lines were selected with Blasticidin and Hygromycin B. Expression of recombinant RAB18 constructs in response to tetracycline was determined by Western blotting and densitometry. (B) Schematic to show experimental approach. Proximity biotinylation of nucleotide-binding-dependent RAB18 interactors is disrupted for the BirA*-RAB18(Ser22Asn) mutant. In contrast, BirA*-RAB18(WT) engages in both GDP-dependent and GTP-dependent interactions. Following affinity purification, nucleotide-binding-dependent interactions are determined by LFQ intensity ratios. (C) Table to show nucleotide-binding-dependent RAB18-associations with BirA*-RAB18(Ser22Asn):BirA*-RAB18(WT) association ratios <0.5.  Orthologous proteins identified by Gillingham et al., 2014, and other studies providing supporting evidence for interactions are shown. Proteins are grouped according to their attributes in the HeLa cell dataset (Figure 1 and Table S1). Instensity ratios were derived individually following normalization by total spectral counts per condition. The full dataset is provided in Table S2.

A

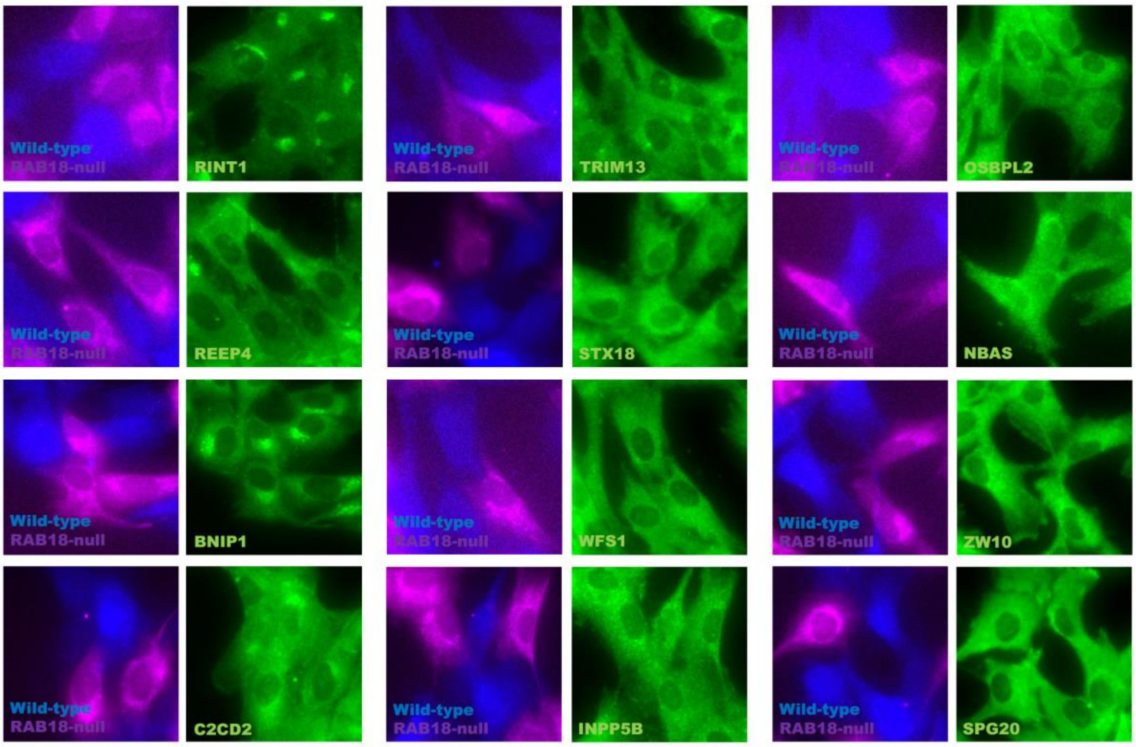


B

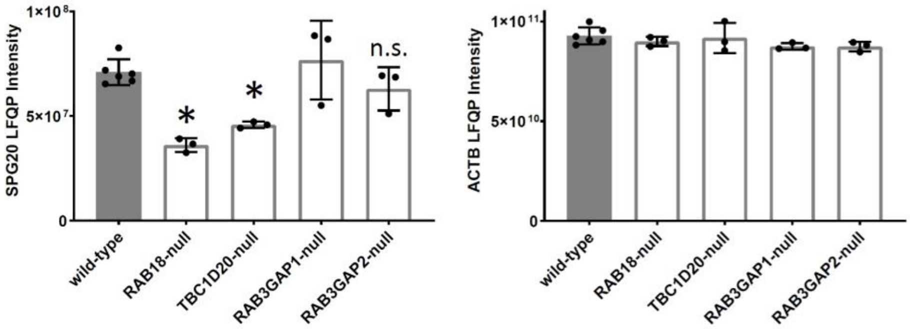


**Figure S4. Comparative fluorescence microscopy of selected RAB18-associated proteins in wild-type and RAB18-null RPE1 cells and quantification of SPG20 levels in RPE1 cells of different genotypes.** Cells of different genotypes were labelled with CellTrace-Violet and CellTrace-Far Red reagents corresponding to blue and magenta channels respectively. Cells were stained with antibodies against indicated proteins in green channel panels. (B) LFQ intensities for SPG20 (Q8N0X7) and β-Actin (P60709) in whole-cell lysates of RPE1 cells of the indicated genotypes. n=3; *p<0.05 following FDR correction. Full dataset provided in Table S3. Error bars represent s.e.m.

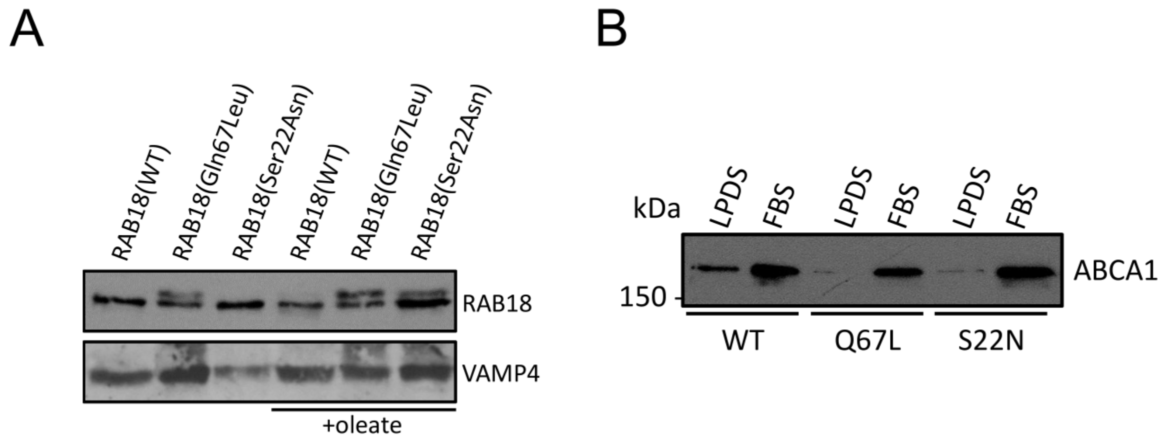


**Figure S5. Characterization of stable CHO cell lines expressing RAB18 constructs.** (A) Comparable expression of RAB18 constructs in stable CHO cell lines. Lysates were prepared from cells cultured in the absence and presence of oleate. Total protein in cell lysates was quantified by Bradford assay. Western blotting shows that levels of RAB18 and VAMP4 are comparable between cell lines. (B) Comparable levels of ABCA1 in in stable CHO cell lines. Lysates were prepared from cells grown in media supplemented with lipoprotein-deficient serum (LPDS) or FBS. Total protein in cell lysates was quantified by Bradford assay. Western blotting shows that levels of ABCA1 are comparable between cell lines under each condition.

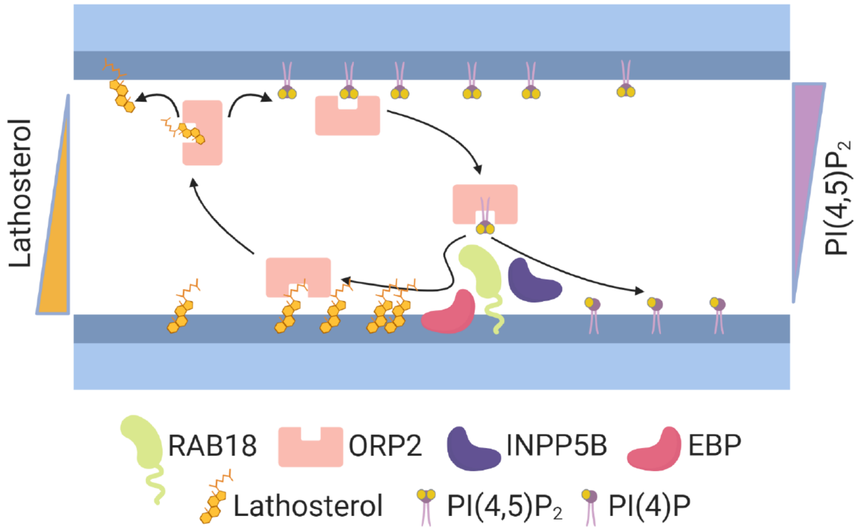


**Figure S6. Model for lathosterol mobilization mediated by RAB18.** ORP2 binds PI(4,5)P2 on an apposed membrane. RAB18 interacts with ORP2 and INPP5B promoting the hydrolysis of PI(4,5)P2 to PI(4)P and maintaining a PI(4,5)P2 concentration gradient. RAB18 coordinates the biosynthesis of lathosterol by EBP and subsequent lathosterol mobilization by ORP2.

SUPPLEMENTARY MATERIALS AND METHODS

Generation of stable T-Rex-293 cell lines

PCR products encoding mouse RAB18, RAB18(Gln67Leu) and RAB18(Ser22Asn) were subcloned into NotI-linearized pcDNA5 FRT/TO FLAG-BirA(Arg118Gly) vector using the In-Fusion HD EcoDry Cloning Plus kit (Takara Bio) according to manufacturer's instructions. Details of PCR templates, primers and target vectors are listed in Table S6. 1.5ug of each recombinant vector together with 13.5ug of pOG44 plasmid (ThermoFisher) were used in cotransfections of T-REx-293 cells, in 10cm dishes, with TransIT-LT1 Transfection Reagent (Mirus Bio, Madison, WI).16 hours following transfection, media was replaced and cells were allowed to recover for 24 hours. Each dish was then split to 4x 10cm dishes in selection media containing 10 ug/ml Blasticidin and 50 ug/ml Hygromycin B. Resistant clones were pooled and passaged once prior to use.

BirA/BioID proximity labelling (T-REx-293 cells)

The T-REx-293 Cell Lines (described above) were seeded onto 3x 15cm plates each and allowed to adhere. Expression of BirA*-RAB18 fusion proteins was induced by treatment with 20ng/ml Tetracycline for 16 hours. Media was then replaced with media containing 20% FBS, 20 ng/ml Tetracycline and 50 uM Biotin and the cells were incubated for a further 8 hours, washed with warmed PBS and pelleted in ice-cold PBS. Cell pellets were snap-frozen and stored at -80ºC prior to lysis. Lysis was carried out in 3ml of ice-cold RIPA buffer (150 mM NaCl, 1% NP40, 0.5% Sodium Deoxycholate, 0.1% SDS, 1mM EDTA, 50mM Tris, pH 7.4) supplemented with complete-mini protease inhibitor cocktail (Roche, Basel, Switzerland), 1mM PMSF, and 62.5 U/ml Benzonase (Merck). Lysates were incubated for 1 hour at 4ºC then sonicated in an ice bath (four 10 second bursts on low power). They were then clarified by centrifugation, and the supernatants transferred to tubes containing pre-washed streptavidin-sepharose (30µl bed-volume)(Merck). The beads were incubated for 3 hours at 4ºC, then washed five times in RIPA buffer and four times in buffer containing 100mM NaCl, 0.025% SDS and 25 mM Tris, pH7.4.

Preparation of cell lysates for label-free quantitative proteomics

RPE1 and HeLa cells were grown to confluence in T75 flasks. They were then trypsinised, and cell pellets were washed with PBS and snap-frozen prior to use. RPE1 pellets were resuspended in 300µl 6M GnHCl, 75mM Tris, pH=8.5. HeLa pellets were resuspended in 300µl 8M urea, 75mM NaCl, 50mM Tris, pH=8.4. In each case, samples were sonicated for 10 minutes using a Bioruptor device together with protein extraction beads (Diagenode). RPE1 samples were heated for 5 minutes at 95ºC. Samples were clarified by centrifugation.

Mass spectrometry

Washed beads from BioID experiments with T-Rex-293 cell lines were resuspended in 50µl 6M urea, 2M thiourea, 10mM Tris, pH=8.5 and DTT was added to 1mM. After 30 minutes incubation at 37ºC, samples were alkylated with 5mM iodoacetamide (IAA) in the dark for 20minutes. DTT was increased to 5mM and 1µg lysC was added, then samples were incubated at 37ºC for 6 hours. Samples were diluted to 1.4M urea, then digested with trypsin (Promega), overnight at 37°C, according to manufacturer's instructions. Samples were acidified by the addition of 0.9% formic acid and 5% acetonitrile. LC-MS was carried out as previously described (Brunet et al., 2016). Briefly, peptides in an aqueous solution containing 5% acetonitrile and 0.1% formic acid were loaded onto a 3 µm PepMap100, 2 cm, 75 µm diameter sample column using an Easy nLC 1000 ultrahigh pressure liquid chromatography system (ThermoFisher). They were eluted with acetonitrile/formic acid into an in-line 50 cm separating column (2 µm PepMap C18, 75 µm diameter) at 40°C. Separated peptides were ionized using an Easy Spray nano source and subjected to MS/MS analysis using a Velos Orbitrap instrument (ThermoFisher). One set of samples was used for the BioID-RAB18 experiment in T-Rex-293 cells (Figure S3, Table S2).

Following acquisition, data were analysed using SEAQUEST software. A NeXprot Human database with 20379 entries was searched. No missed cleavages were permitted. Fixed modification by carbamidomethylation of cysteine residues was considered. Variable modification by oxidation or hydroxylation of methionine residues was considered. Mass tolerance for precursor ions was ±2m/z and that for fragment ions was ±1m/z. Thresholds for accepting individual spectra were set at $p<0.05$. A %FDR of 0.25% was calculated using the PeptideProphet package (http://peptideprophet.sourceforge.net/). Single-peptide identifications of proteins were removed.

RPE1 lysates were reduced and alkylated through addition of tris(2-carboxyethyl)phosphine (TCEP) and 2-chloroacetamide (CAA) to 5mM and 10mM respectively and then incubated at 95°C for 5 minutes. After cooling, samples were diluted to 3M guanidine and 0.5µg lysC added with incubation overnight at 37°C. A further dilution to 1M guanidine was followed by digest with 0.3µg trypsin at 37°C for 4 hours. Samples were acidified with TFA. HeLa lysates were reduced and alkylated by addition of DTT to 10mM, then by addition of IAA to 25mM, then further addition of DTT to 25mM, with incubation at room temperature for 30-60 minutes following each step. Samples were digested with lysC, overnight at 37°C. They were then diluted to 2M urea, and further digested, overnight at 37°C. Samples were acidified with TFA. Trypsin cleaves on the C-terminal side of lysine and arginine residues unless the C-terminal residue is proline. Hydrolysis is slower where the C-terminal residue is acidic. Lys-C cleaves on the C-terminal side of lysine residues. Peptides were loaded on to activated (methanol), equilibrated (0.1% TFA) C18 stage tips before being washed with 0.1% TFA and eluted with 0.1% TFA/80 acetonitrile. The organic was dried off, 0.1% TFA added to 15 µl and 5 µl injected onto LC-MS. Peptides were separated on an Ultimate nano HPLC instrument (ThermoFisher), and analysed on either an Orbitrap Lumos or a Q Exactive Plus instrument (ThermoFisher).

Three sets of replicate samples were used to generate the RPE1 quantitative proteomics dataset (Figure S4, Table S3). Each set of samples was grown and harvested independently. Six sets of replicate samples were used to generate the HeLa cell quantitative proteomics dataset (Table S4). Two different wild-type clones and two different TBC1D20-null genotypes were used (three replicates each). These can be considered biological replicates.

After data-dependent acquisition of HCD fragmentation spectra, data were analysed using MaxQuant (version 1.6.2.10 for the RPE1 experiment and version 1.5.7.4 for the HeLa cell experiment). For the RPE1 experiment, the Uniprot Human 2018_07 database with 21050 entries was searched. For the HeLa cell experiment, the

Uniprot Human 2017_01 database with 21031 entries was searched. 2 missed/non-specific cleavages were permitted. Fixed modification by carbamidomethylation of cysteine residues was considered. Variable modification by oxidation of methionine residues and N-terminal acetylation were considered. Mass error was set at 20 ppm for the first search tolerance and 4.5 ppm main search tolerance. Thresholds for accepting individual spectra were set at $p<0.05$. Single-peptide identifications of proteins were removed. %FDR was estimated at <5% using the decoy search method. Additional parameters and gradients used for separation are provided in Table S6. Quantification data were produced with MaxLFQ [19]. p values for comparisons between LFQ intensities of each protein in samples from test and wild-type genotypes were first calculated by Student's t-tests. p values were then adjusted for multiple testing using an online calculator (https://www.sdmproject.com/utilities/?show=FDR).
