## Supplementary material for "Comparative proximity biotinylation implicates RAB18 in sterol mobilization and biosynthesis": single peptide identifications

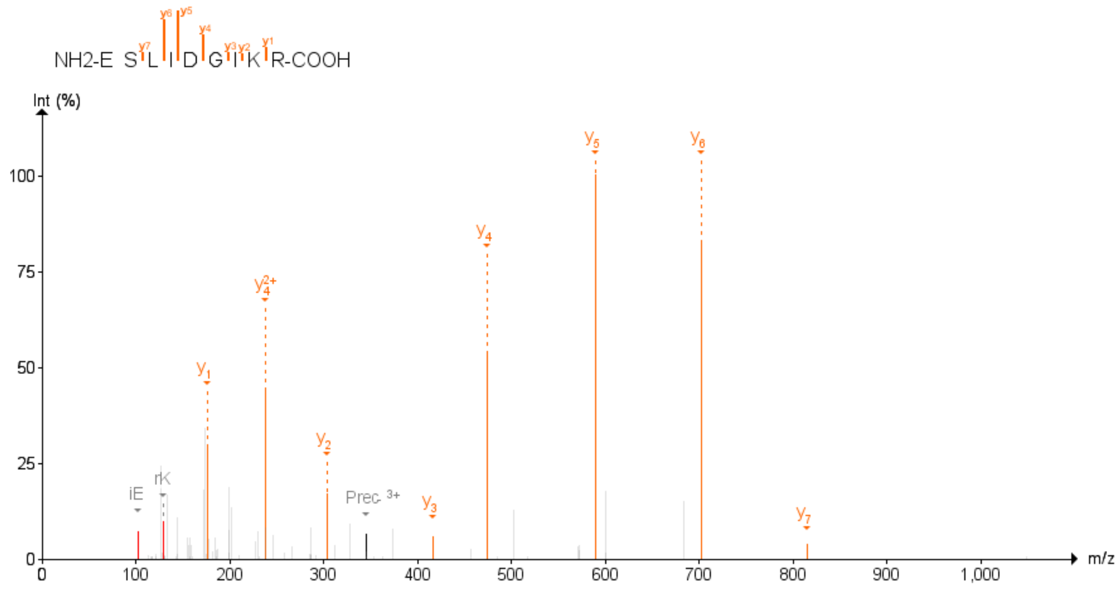

Figure 1: AHCY First experiment

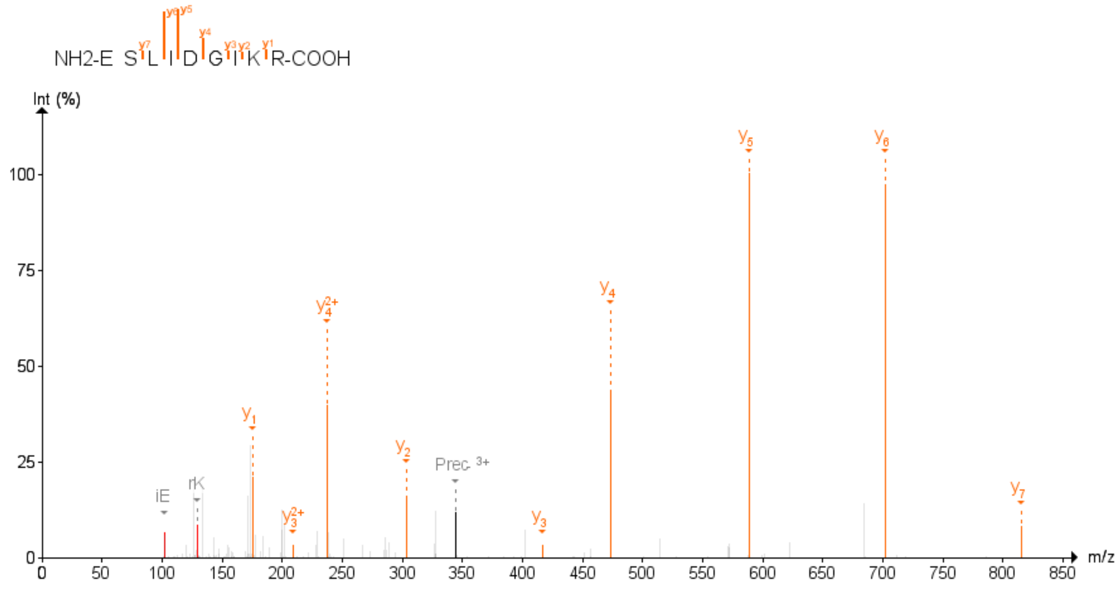

Figure 2: AHCY Second experiment

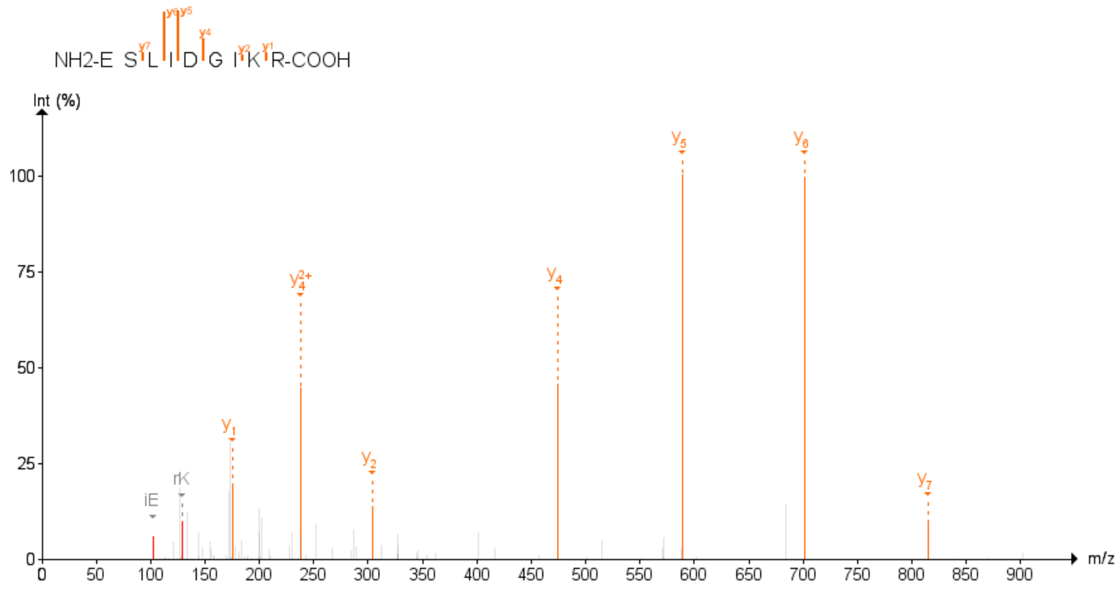

Figure 3: AHCY Third experiment

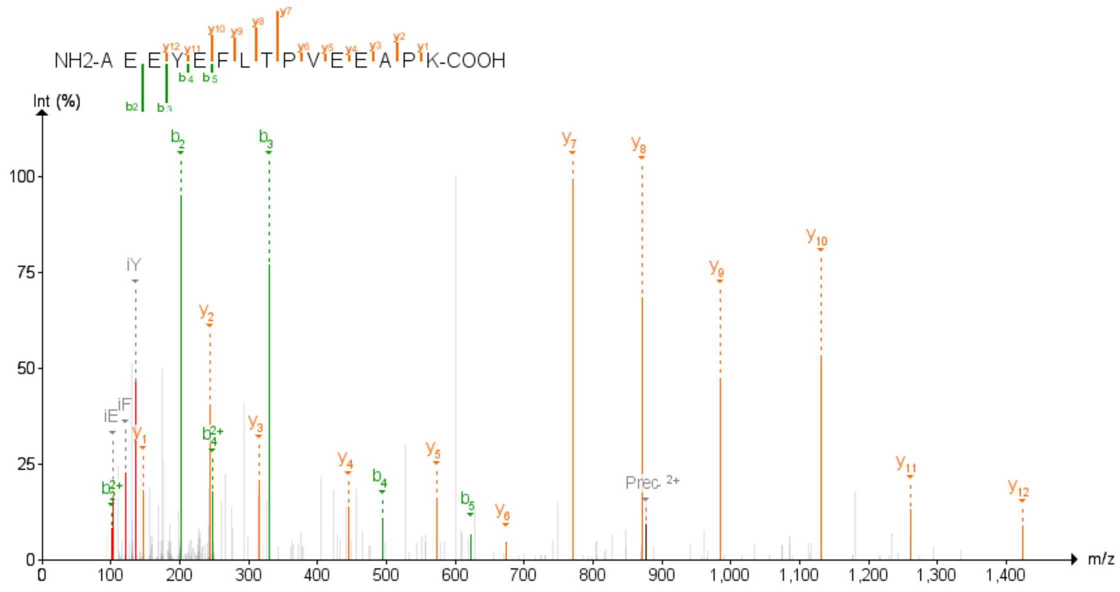

Figure 4: ARHGDIA Second experiment

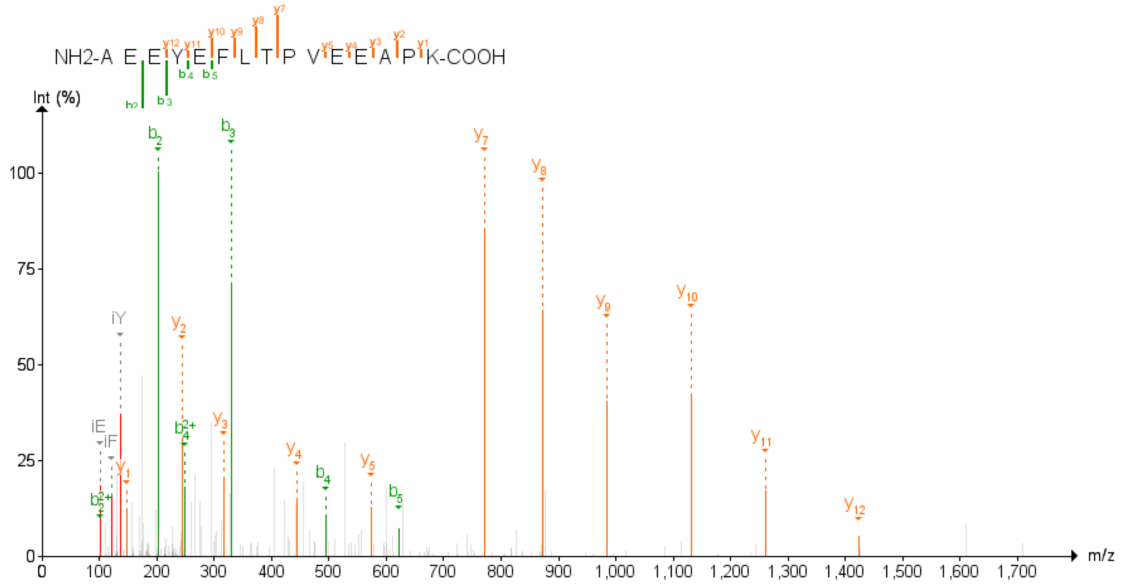

Figure 5: ARHGDIA Third experiment

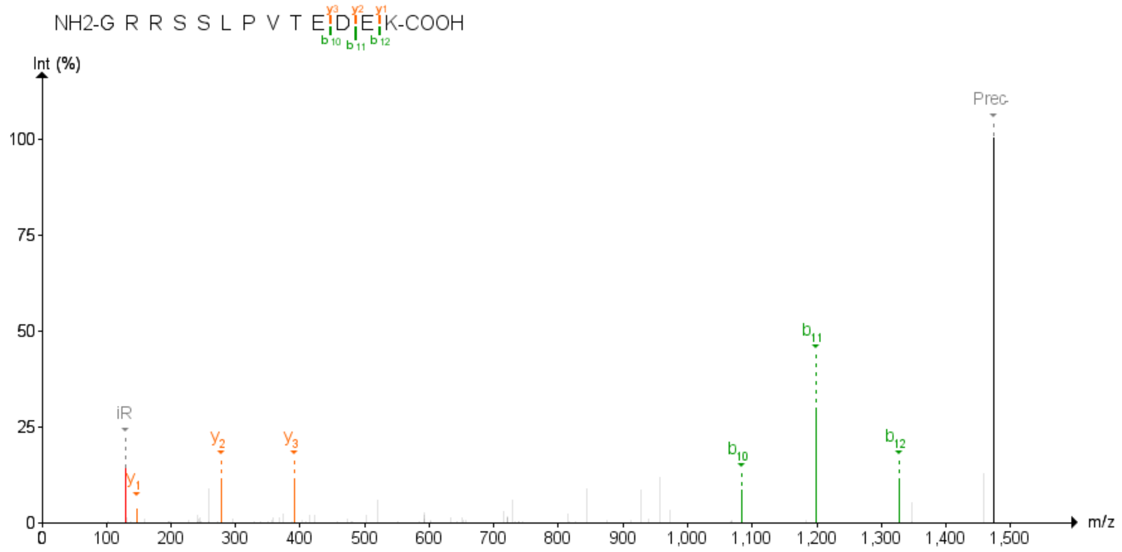

Figure 6: ARID4A Second experiment

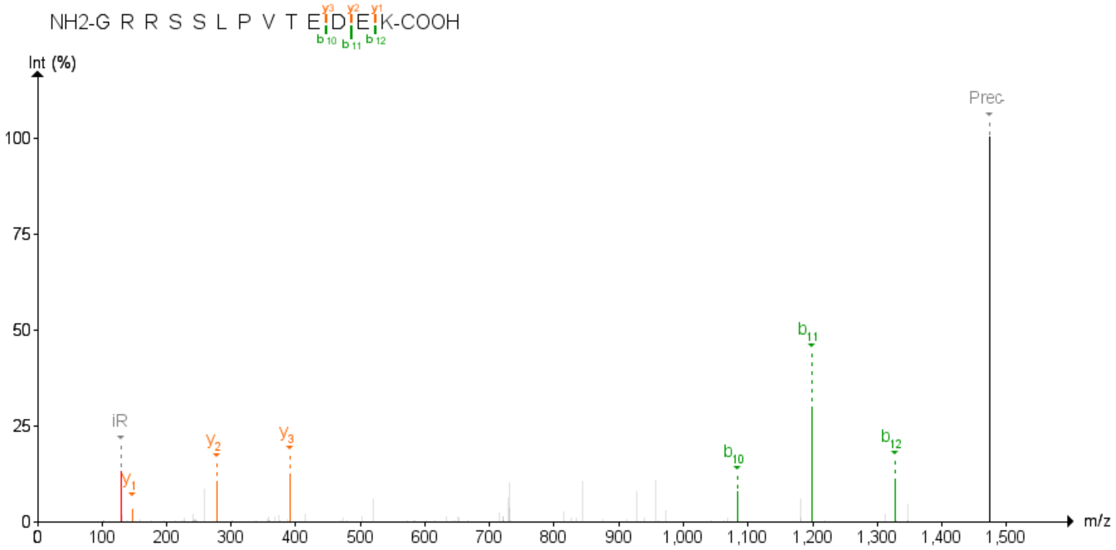

Figure 7: ARID4A Third experiment

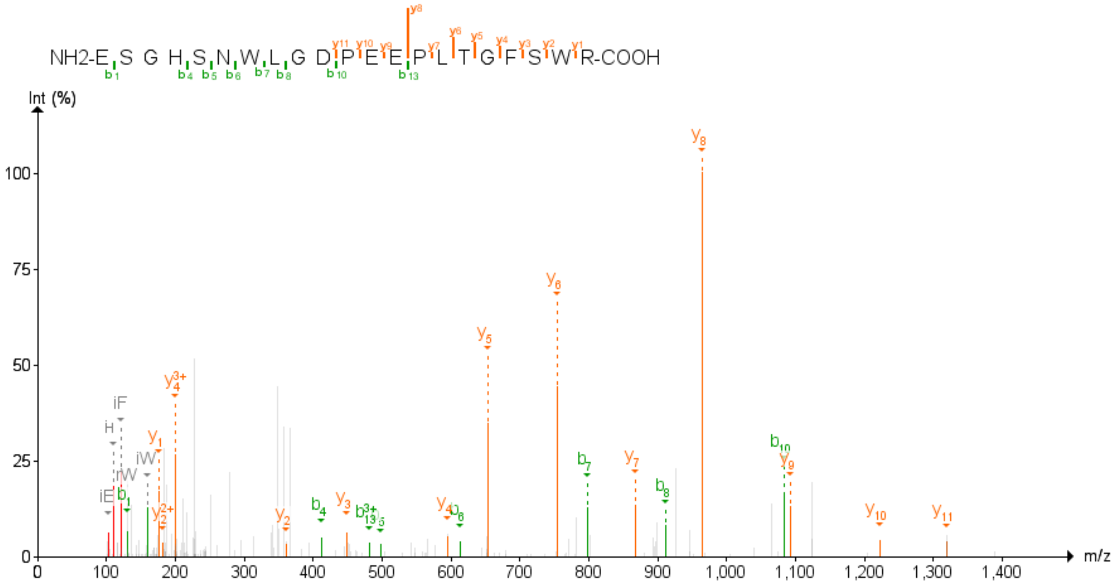

Figure 8: ATL3 First experiment

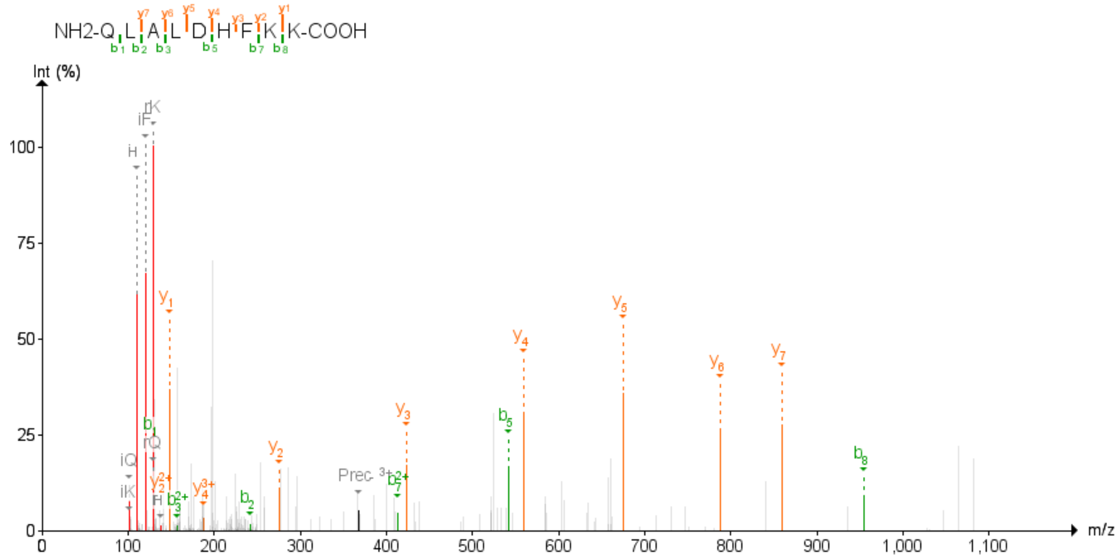

Figure 9: ATL3 Second experiment

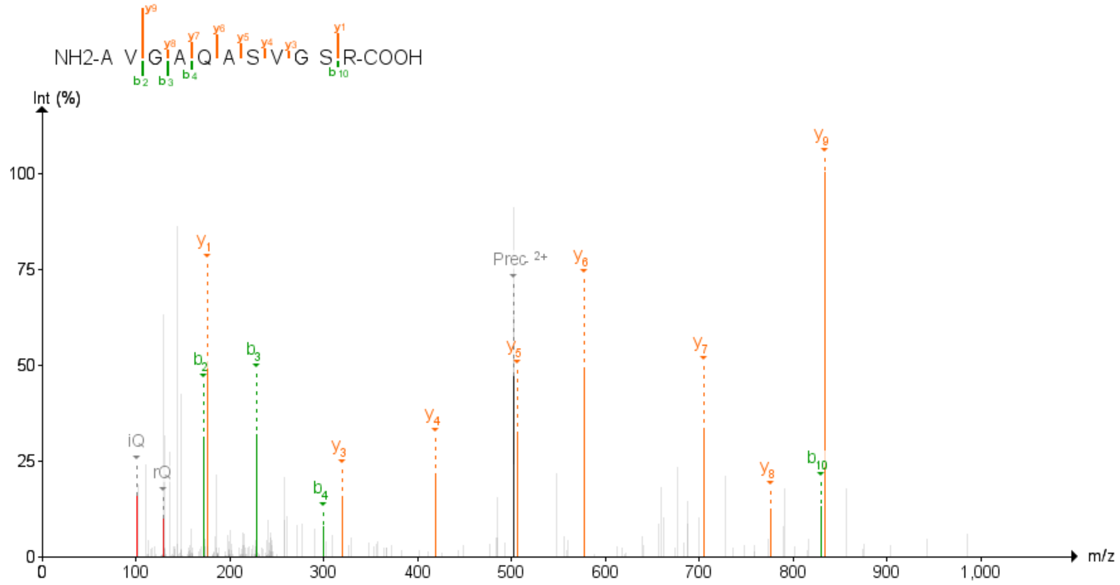

Figure 10: BABAM1 Second experiment

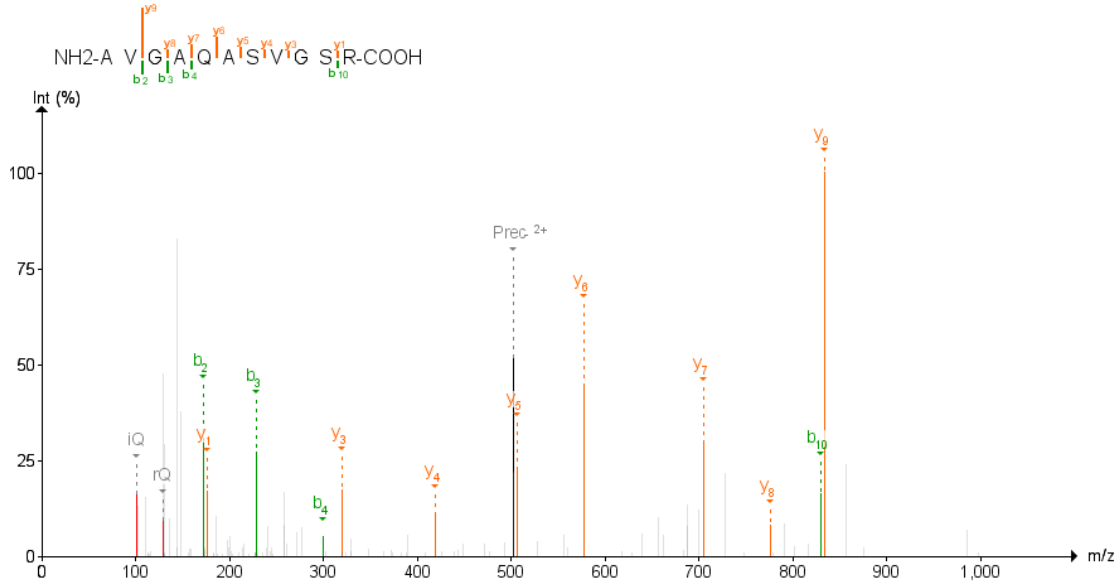

Figure 11: BABAM1 Third experiment

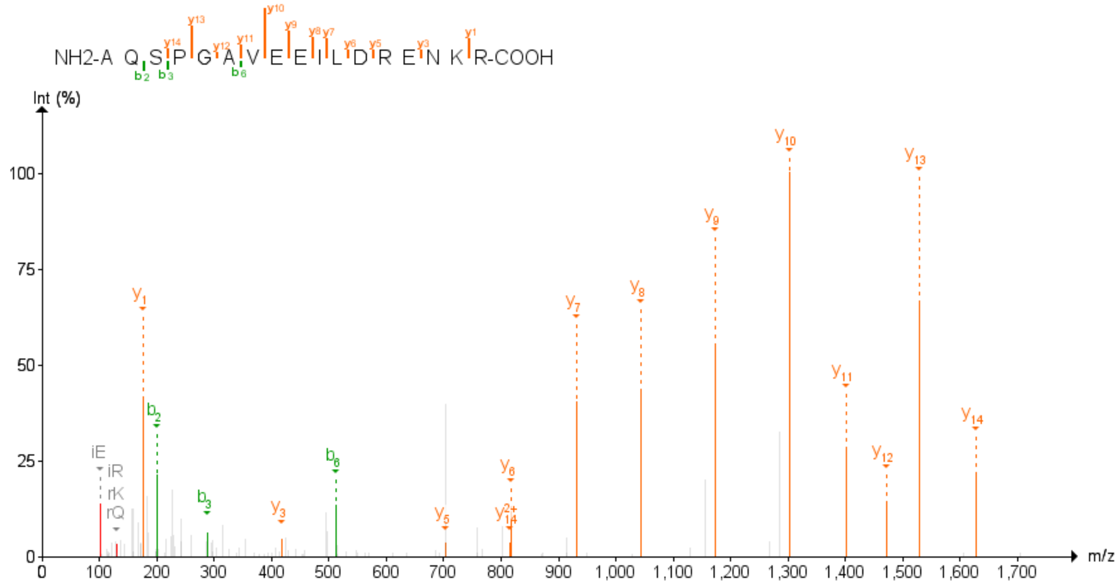

Figure 12: BET1L First experiment

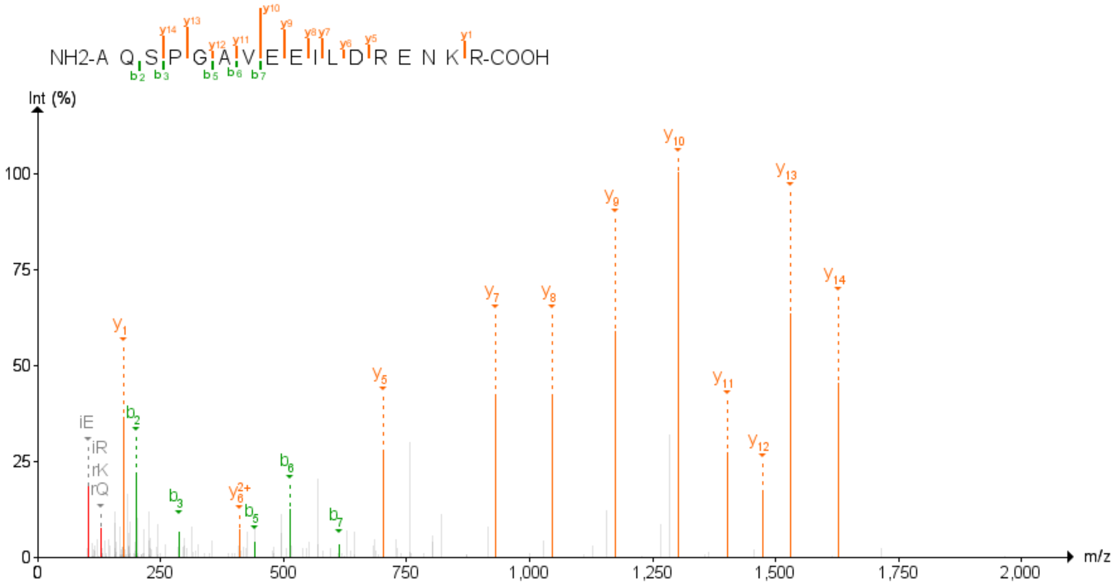

Figure 13: BET1L Second experiment

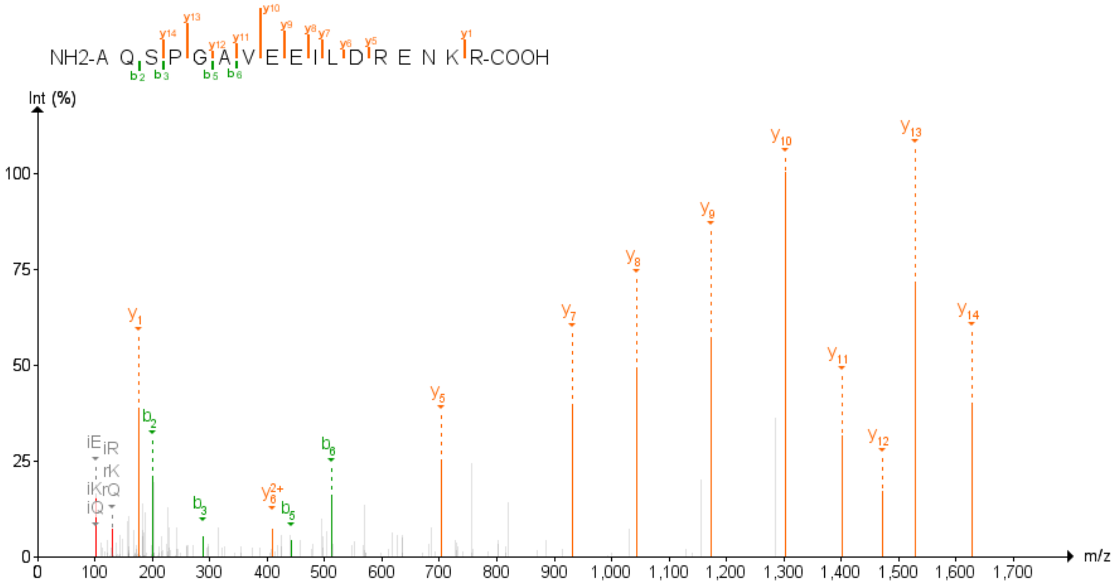

Figure 14: BET1L Third experiment

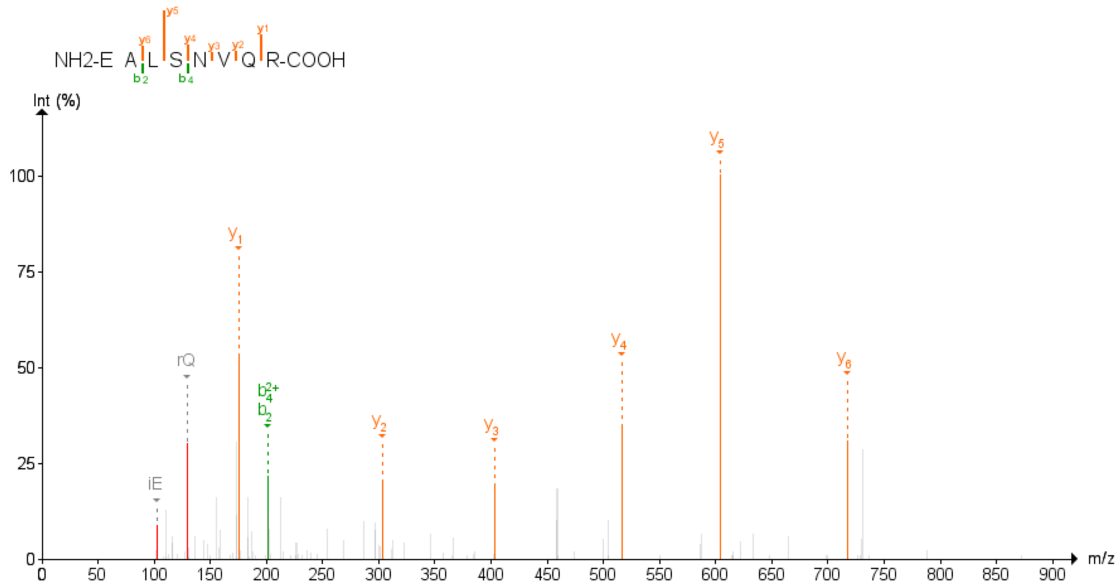

Figure 15: c11orf49 First experiment

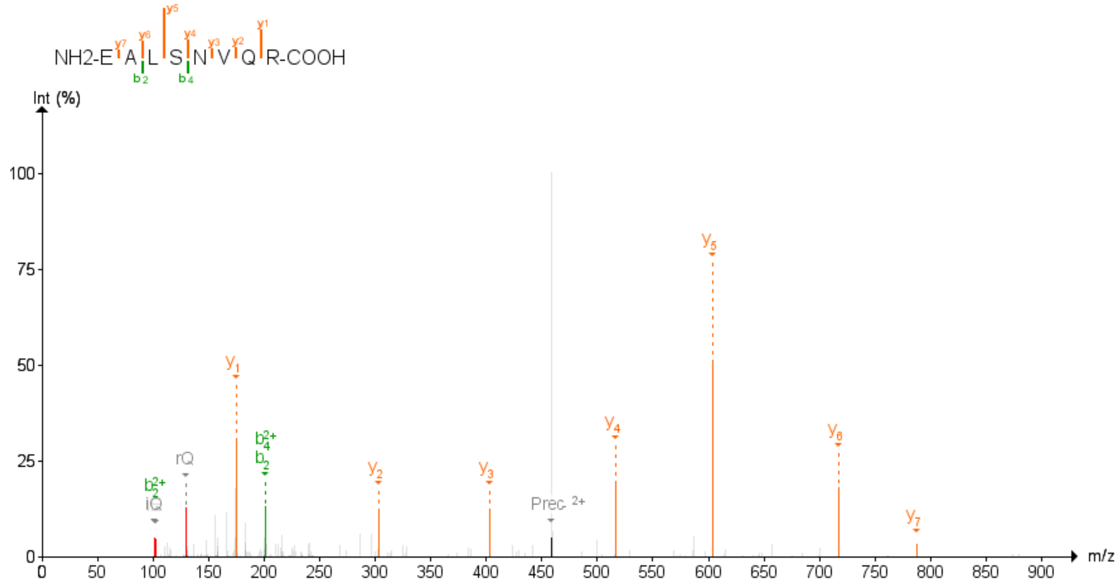

Figure 16: c11orf49 Second experiment

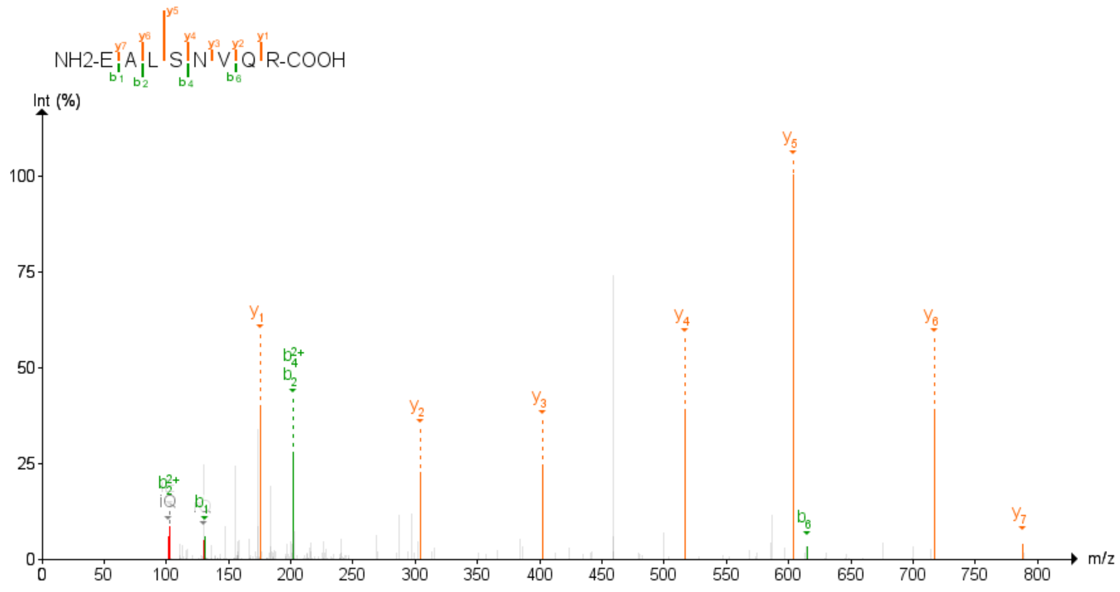

Figure 17: c11orf49 Third experiment

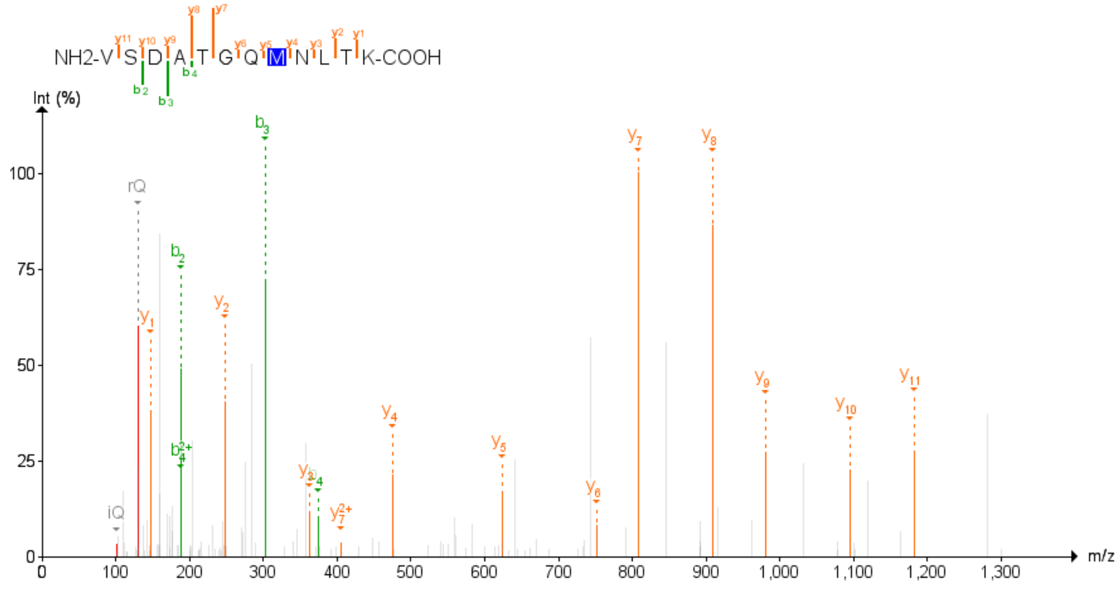

Figure 18: CAPG Second experiment

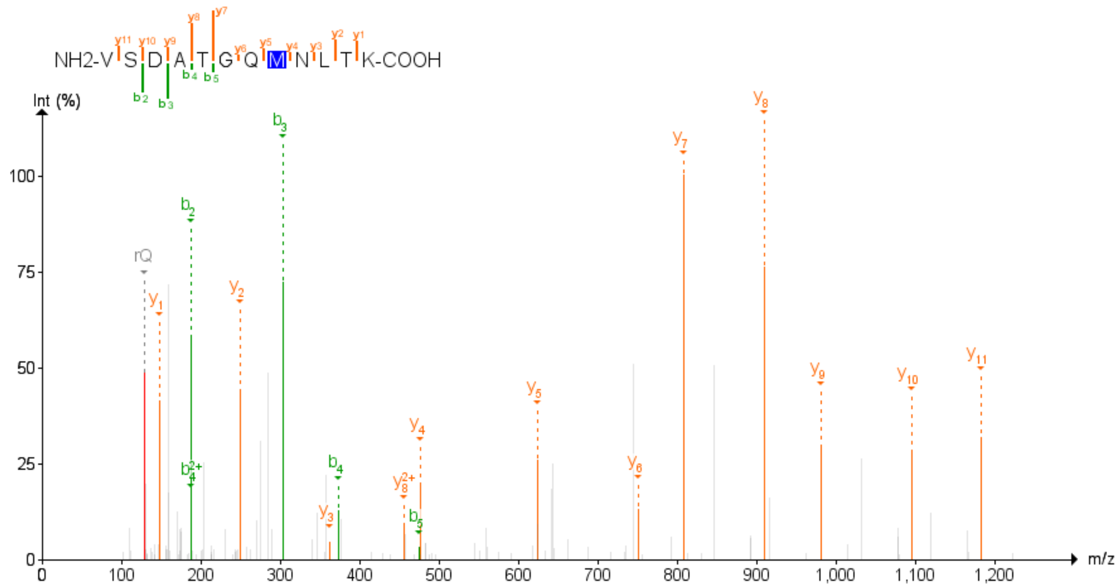

Figure 19: CAPG Third experiment

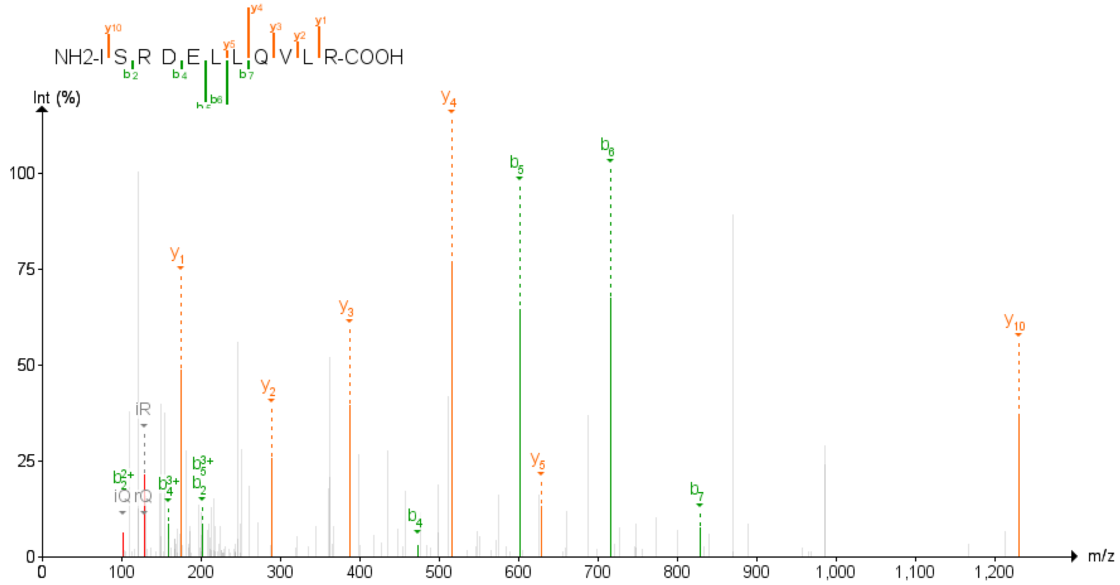

Figure 20: CHP1 Second experiment

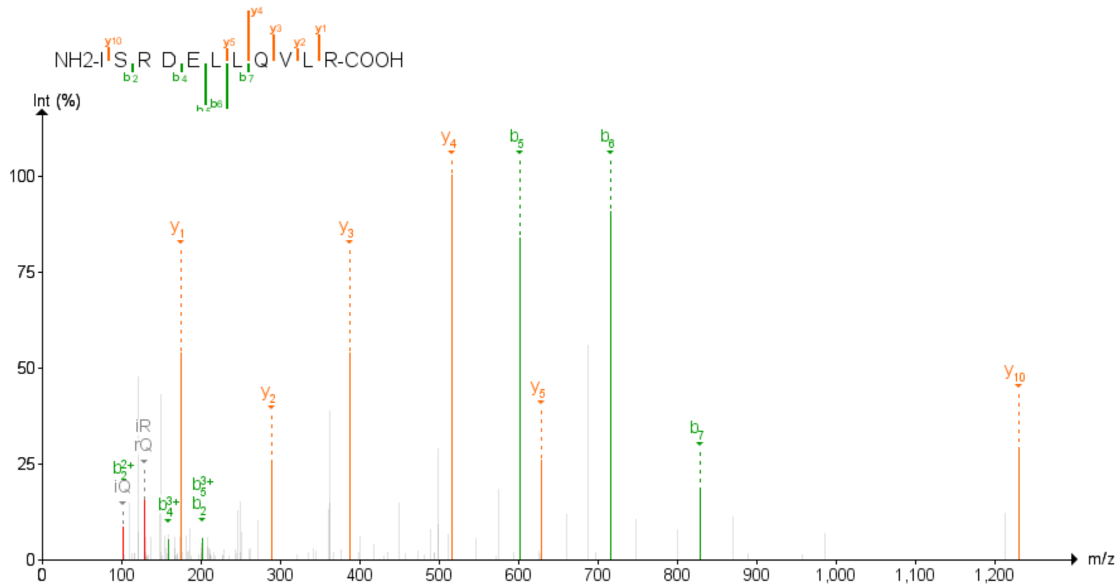

Figure 21: CHP1 Third experiment

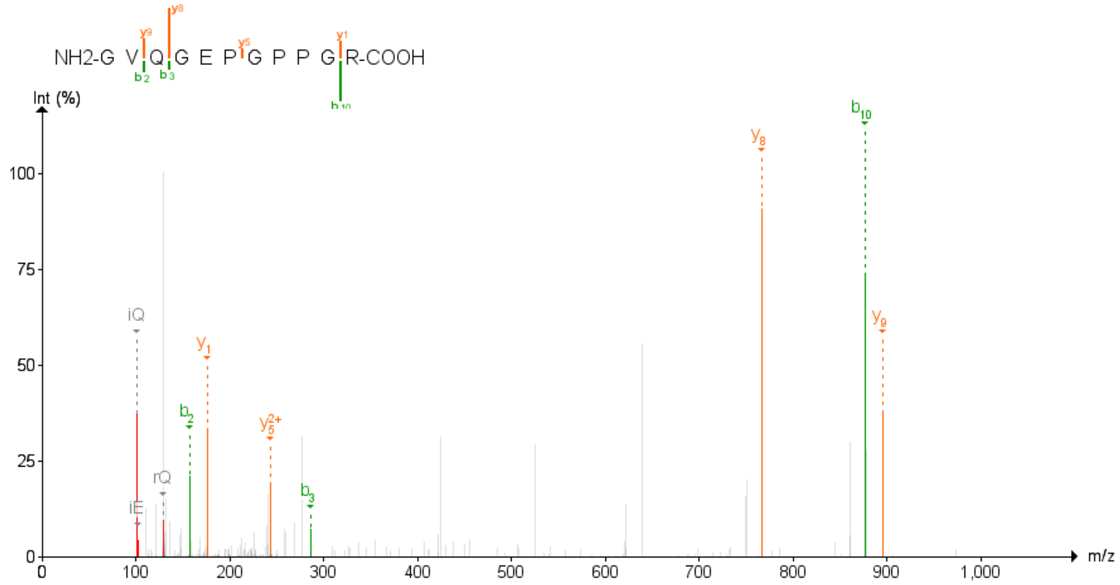

Figure 22: COL16A1 Second experiment

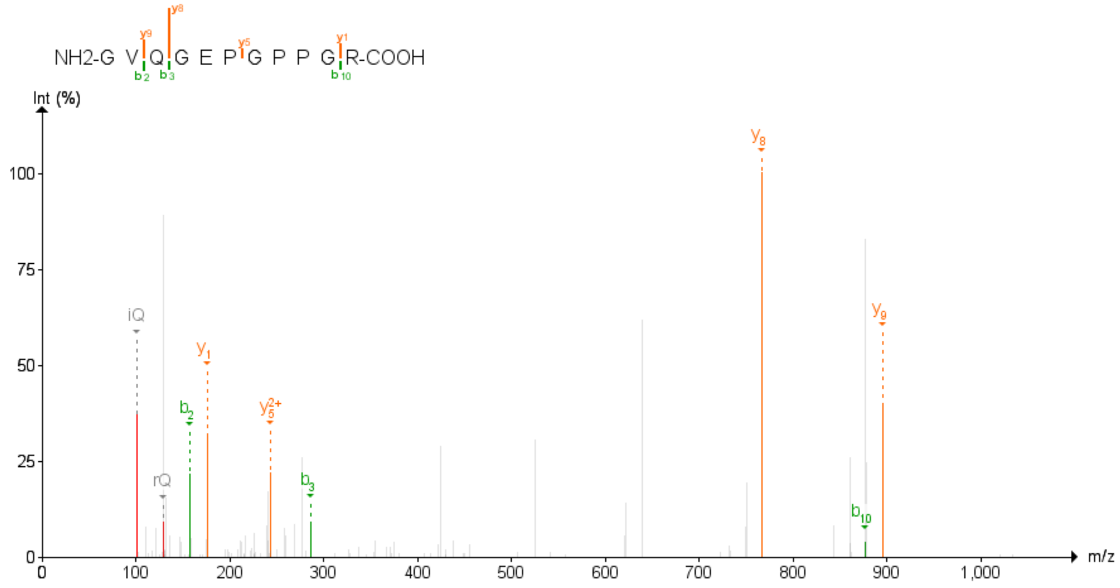

Figure 23: COL16A1 Third experiment

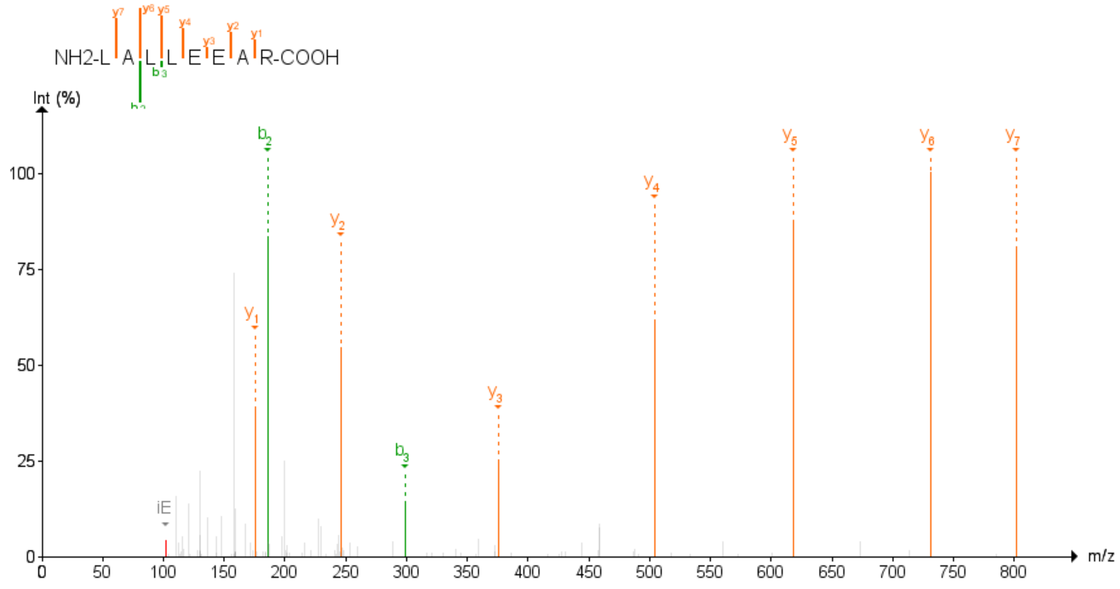

Figure 24: CROCC First experiment

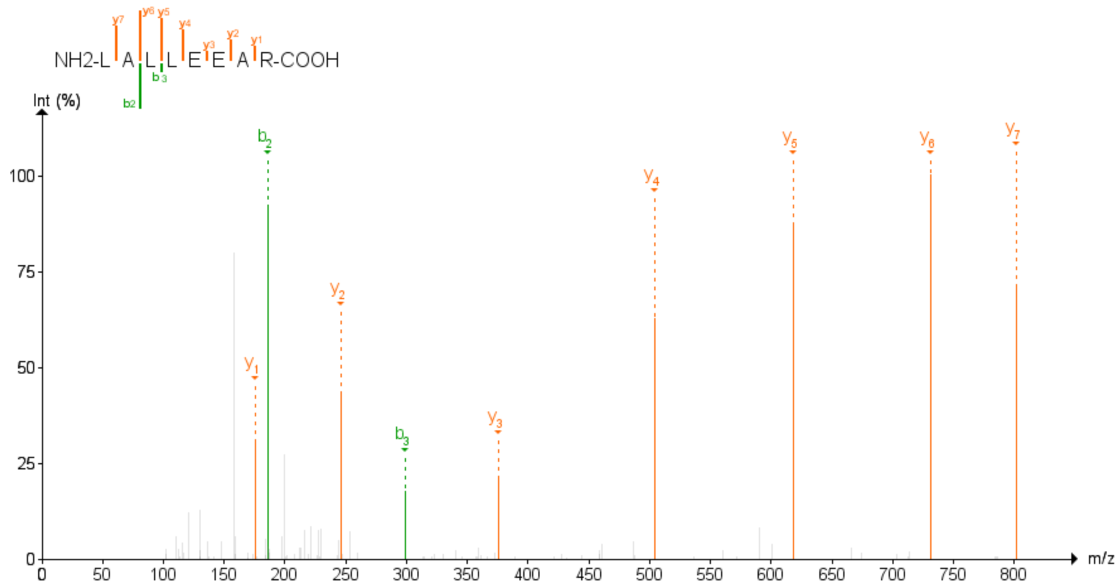

Figure 25: CROCC Third experiment

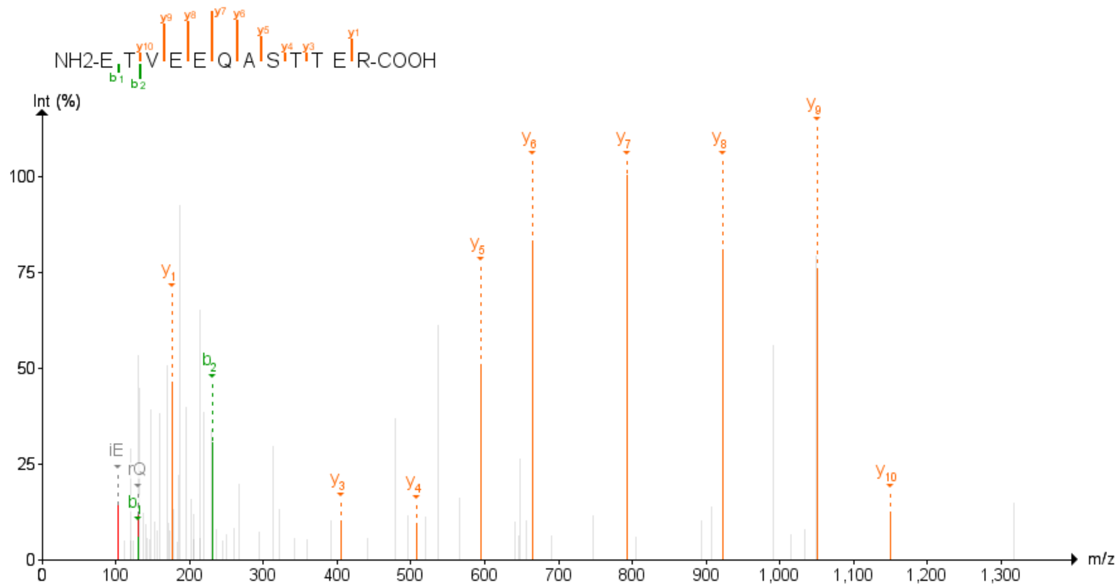

Figure 26: CUL4B First experiment

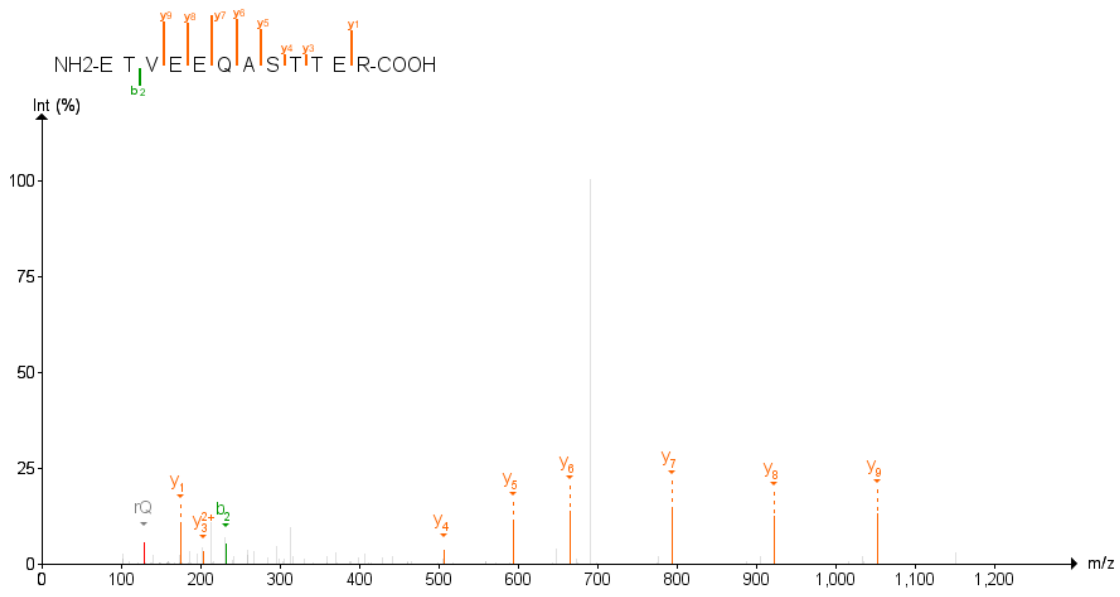

Figure 27: CUL4B Third experiment

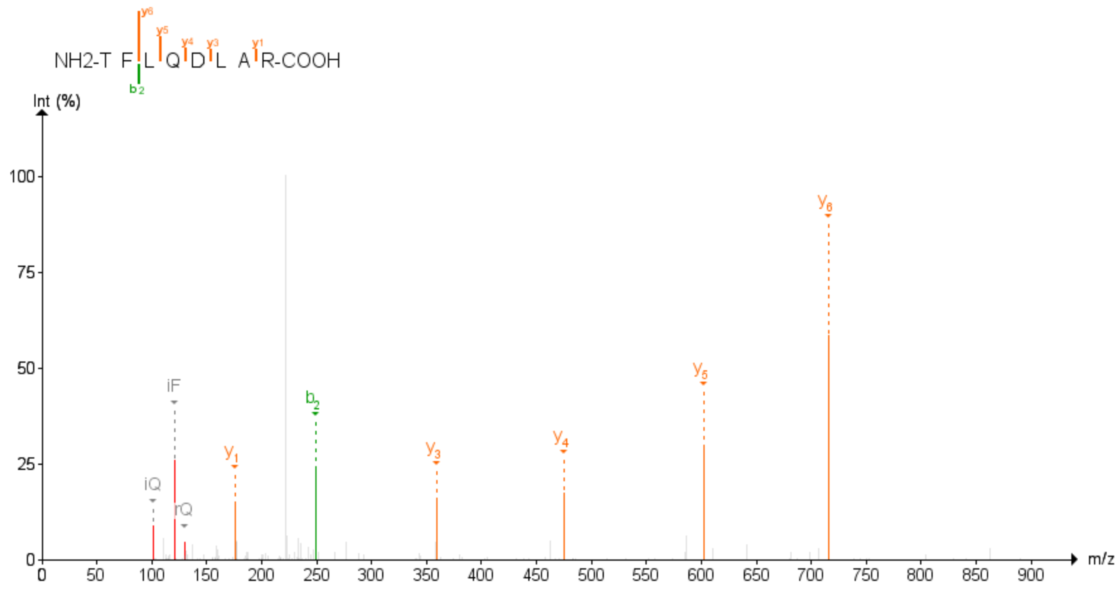

Figure 28: EI24 Second experiment

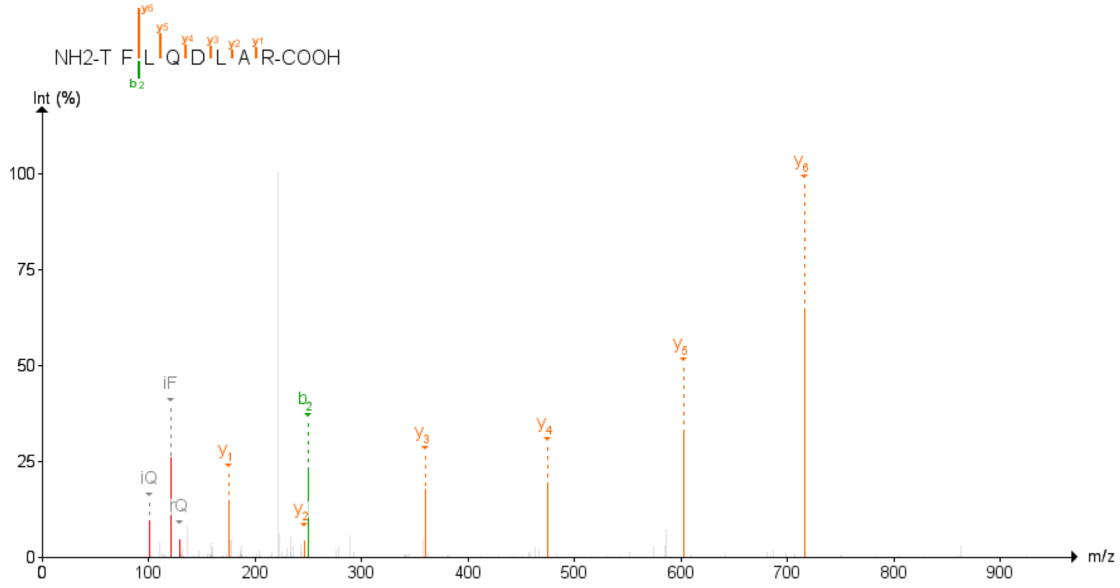

Figure 29: EI24 Third experiment

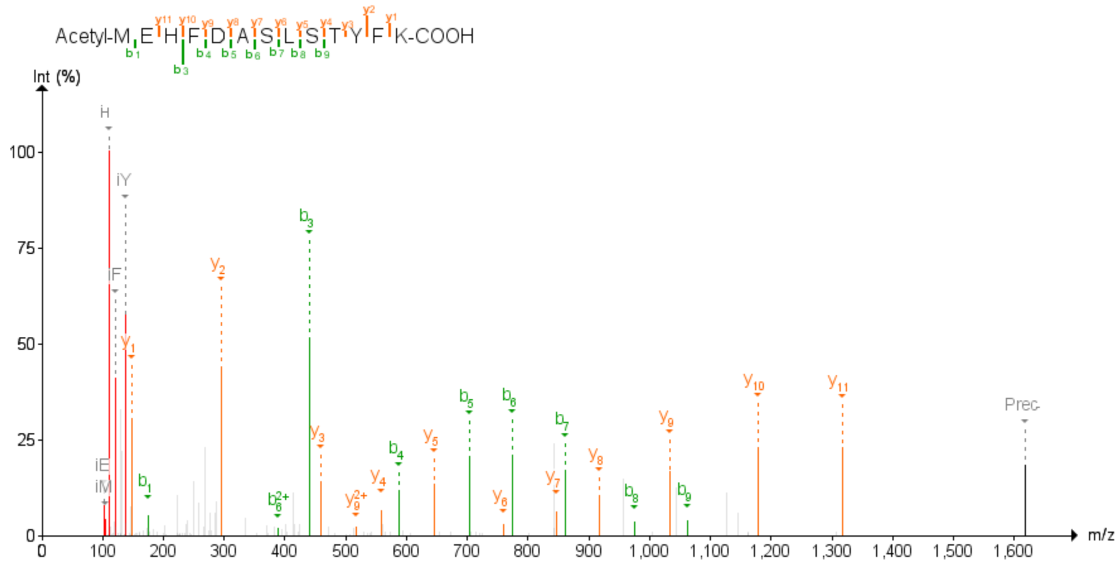

Figure 30: ELOVL5 First experiment

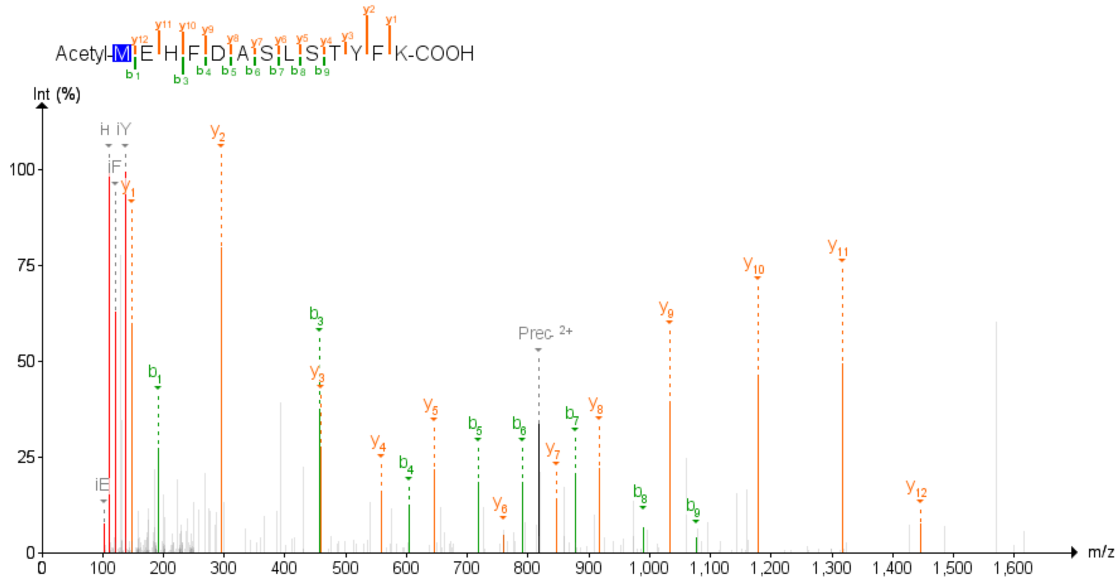

Figure 31: ELOVL5 Second experiment

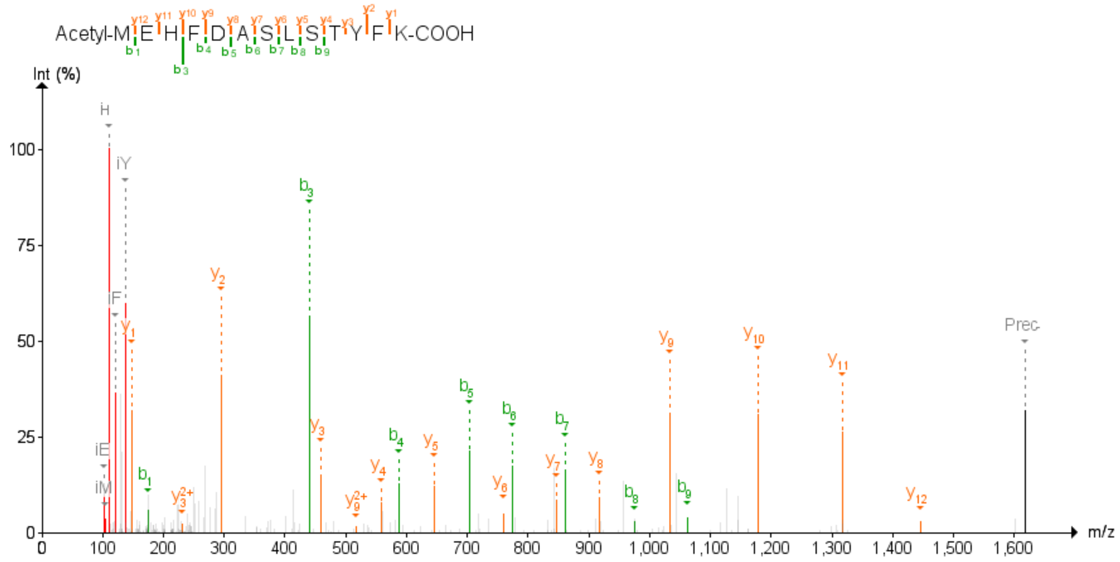

Figure 32: ELOVL5 Third experiment

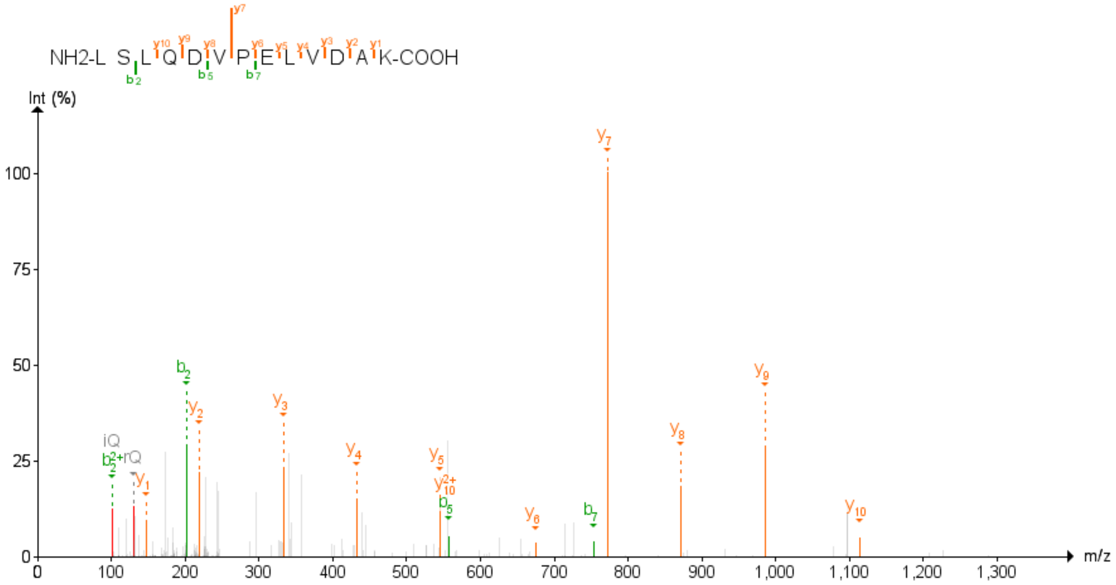

Figure 33: FAM101B First experiment

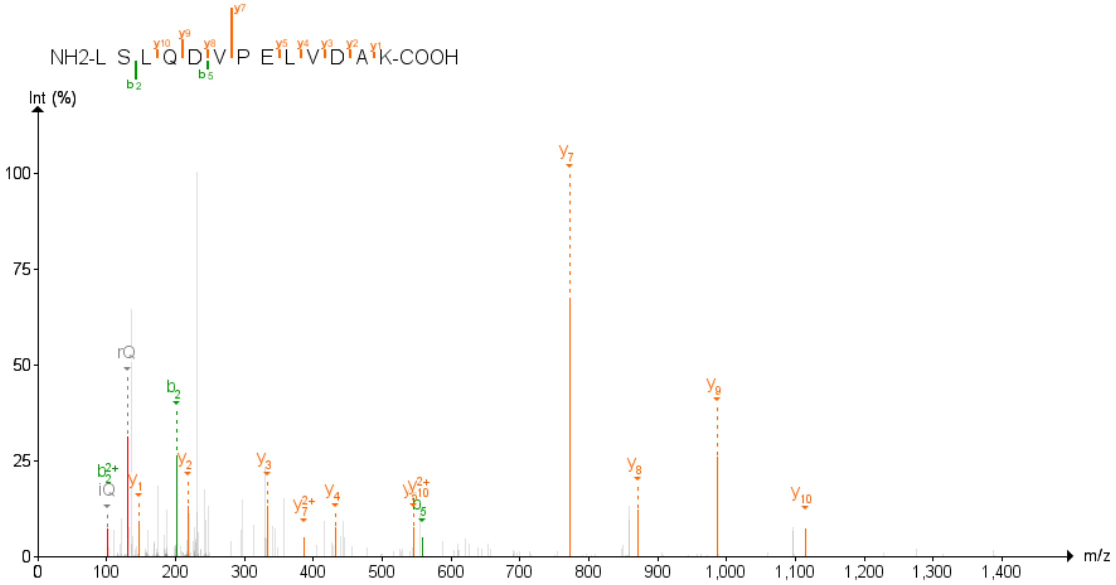

Figure 34: FAM101B Second experiment

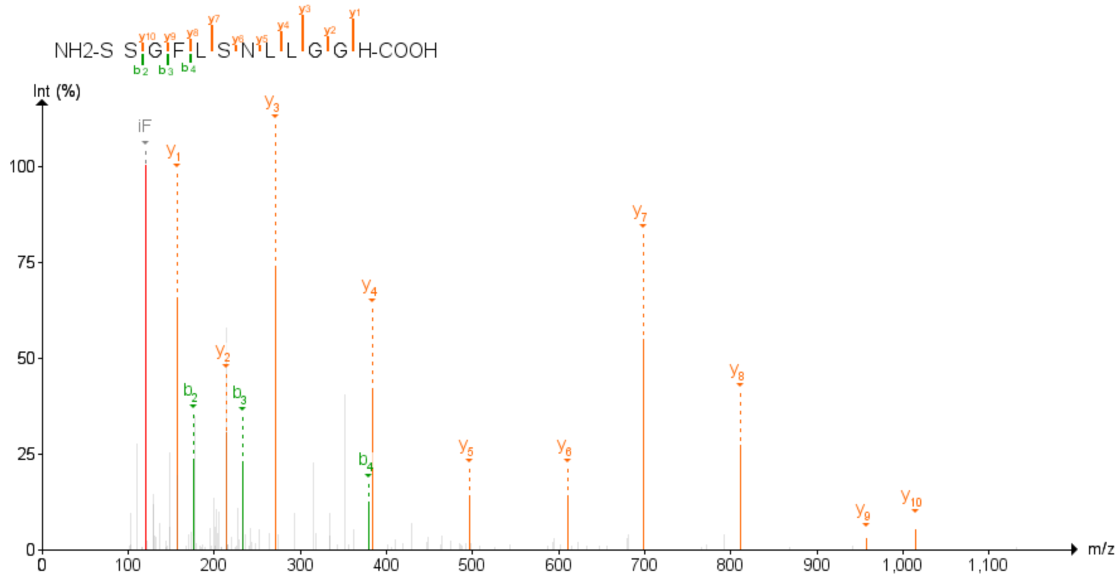

Figure 35: FAM134B First experiment

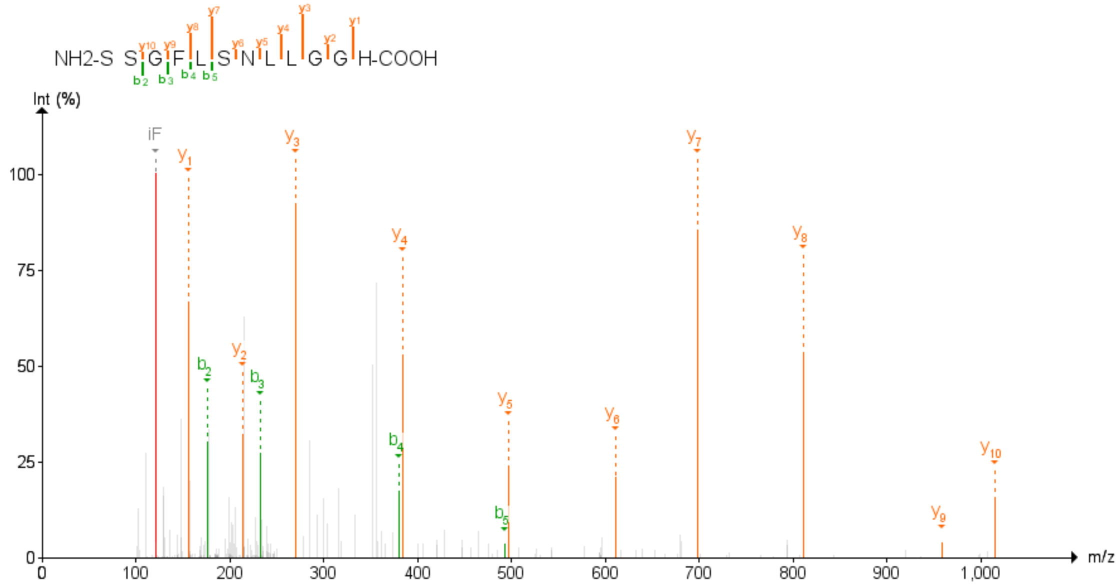

Figure 36: FAM134B Second experiment

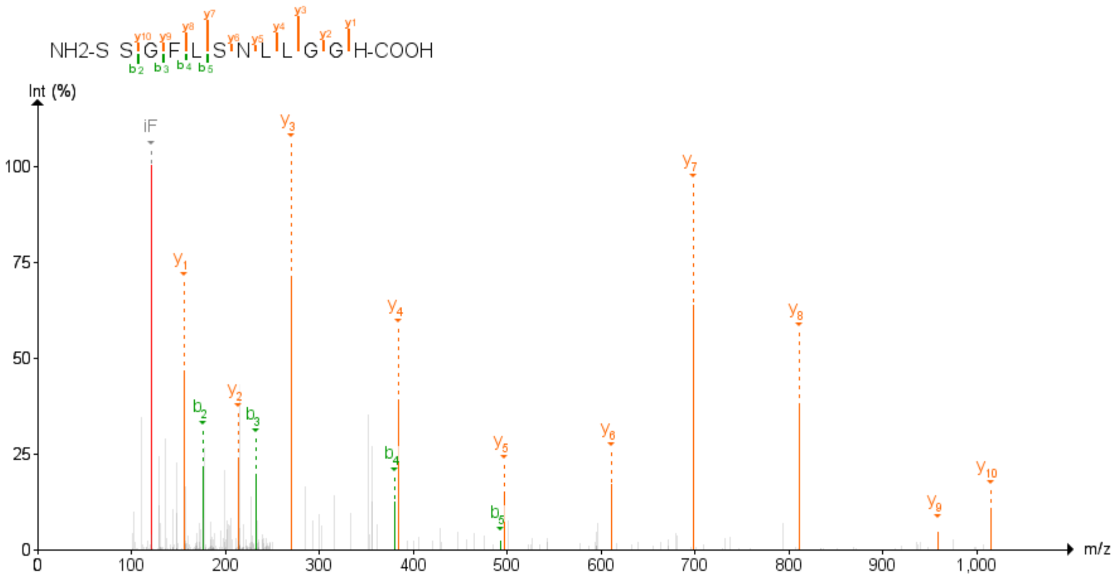

Figure 37: FAM134B Third experiment

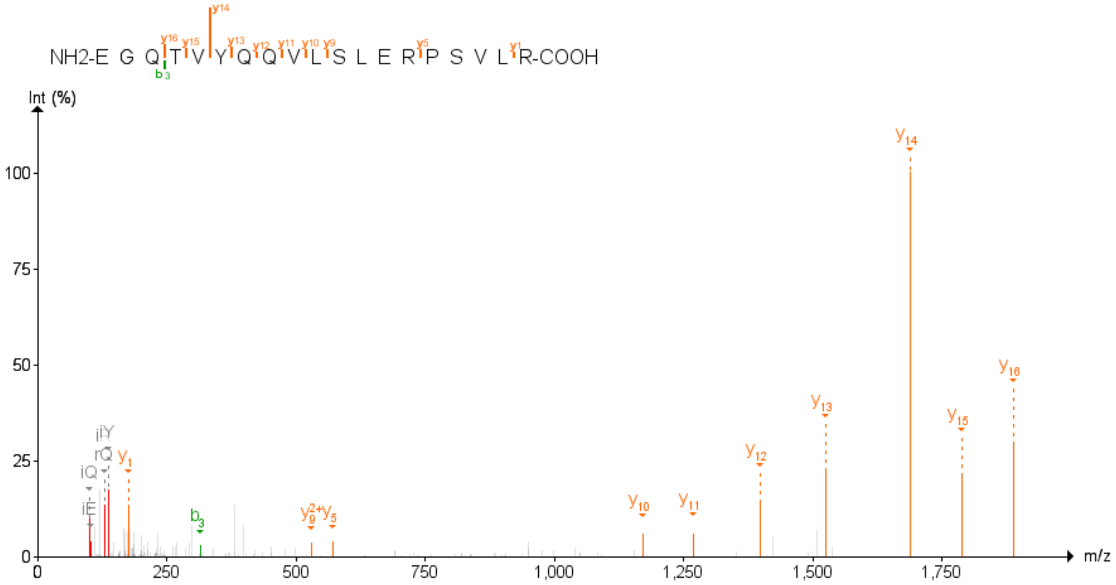

Figure 38: GBA2 First experiment

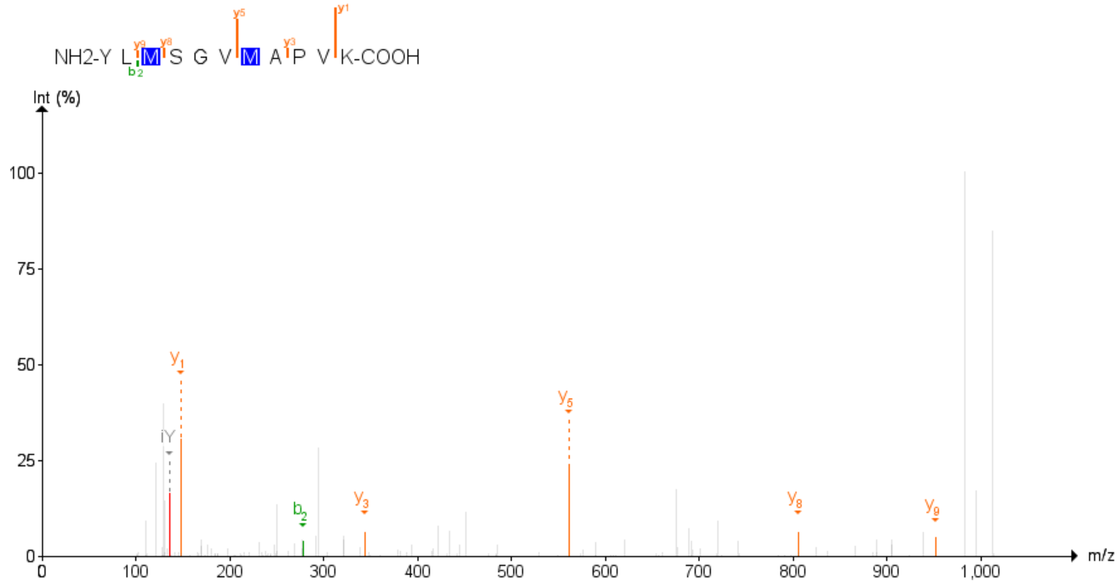

Figure 39: GBA2 Second experiment

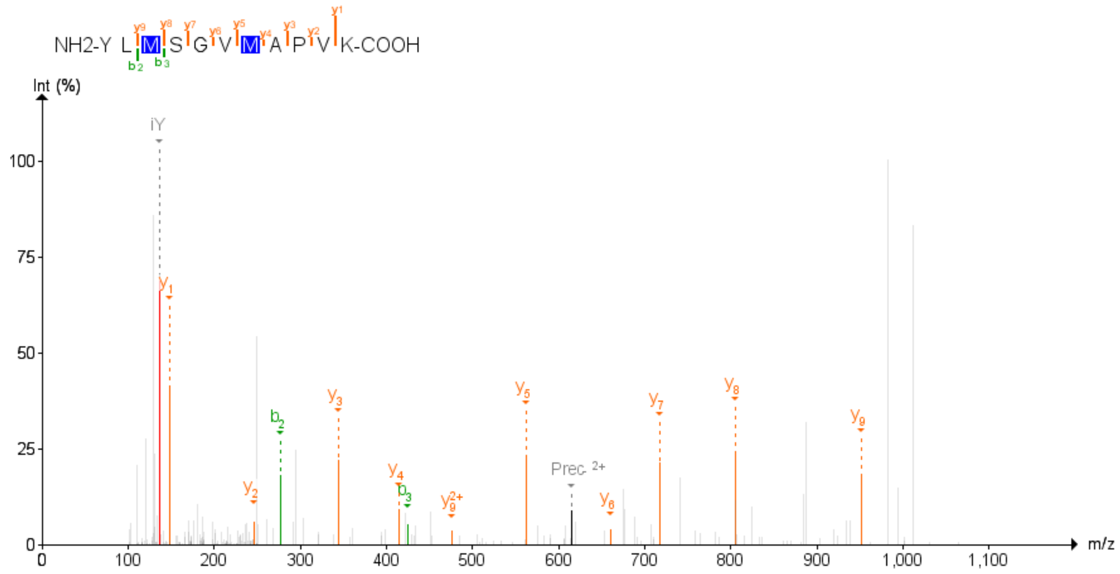

Figure 40: GBA2 Third experiment

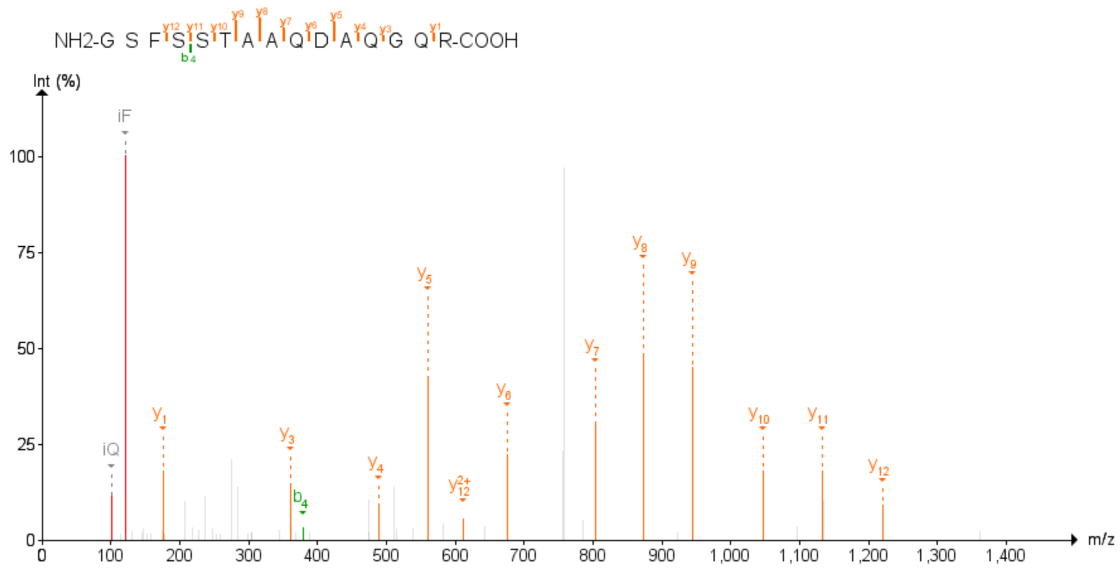

Figure 41: GPR180 First experiment

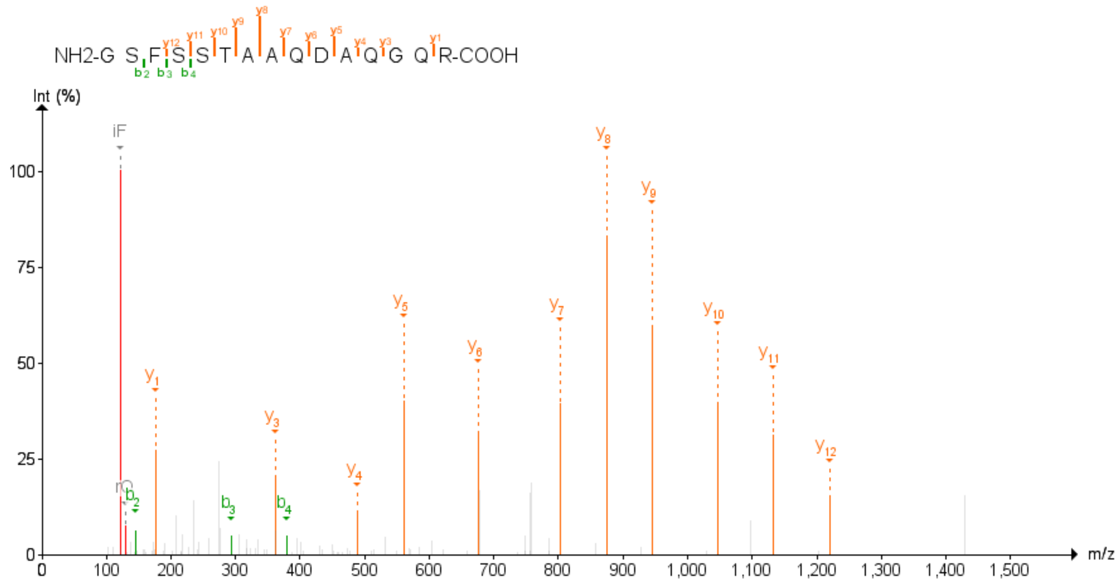

Figure 42: GPR180 Second experiment

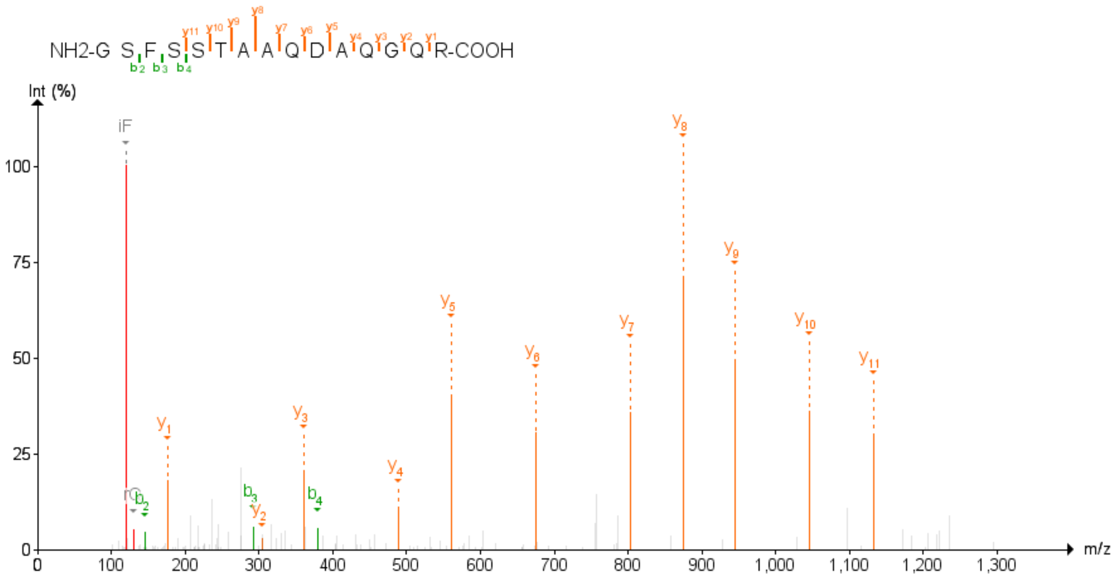

Figure 43: GPR180 Third experiment

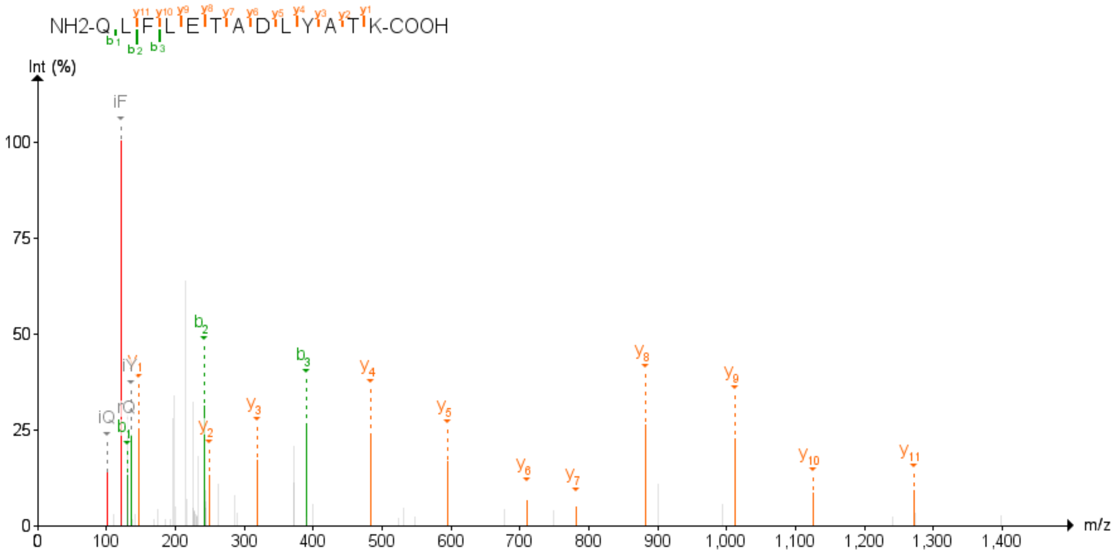

Figure 44: GPR89B;GPR89A First experiment

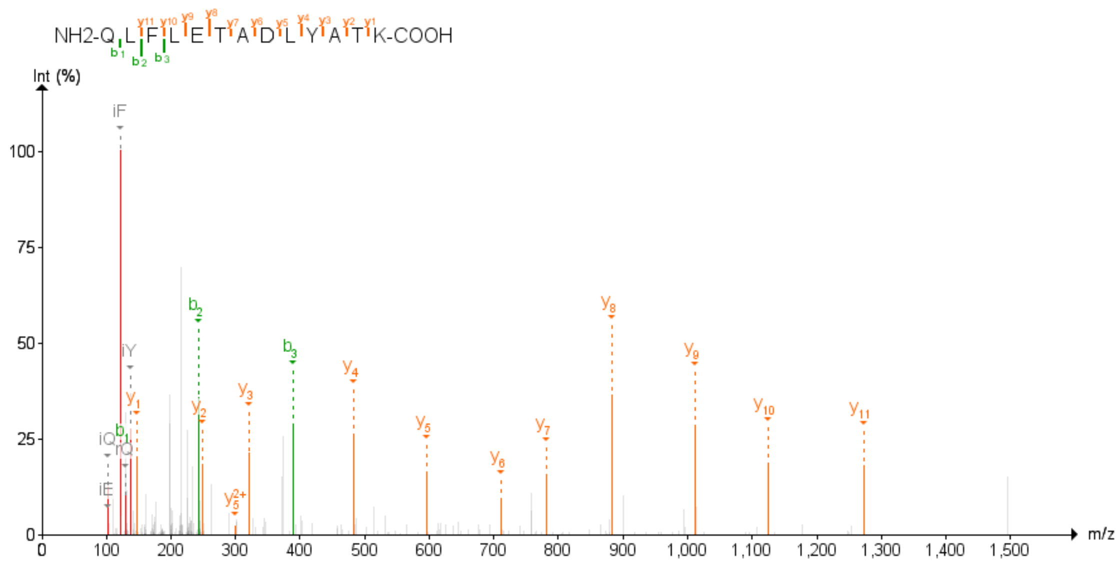

Figure 45: GPR89B;GPR89A Second experiment

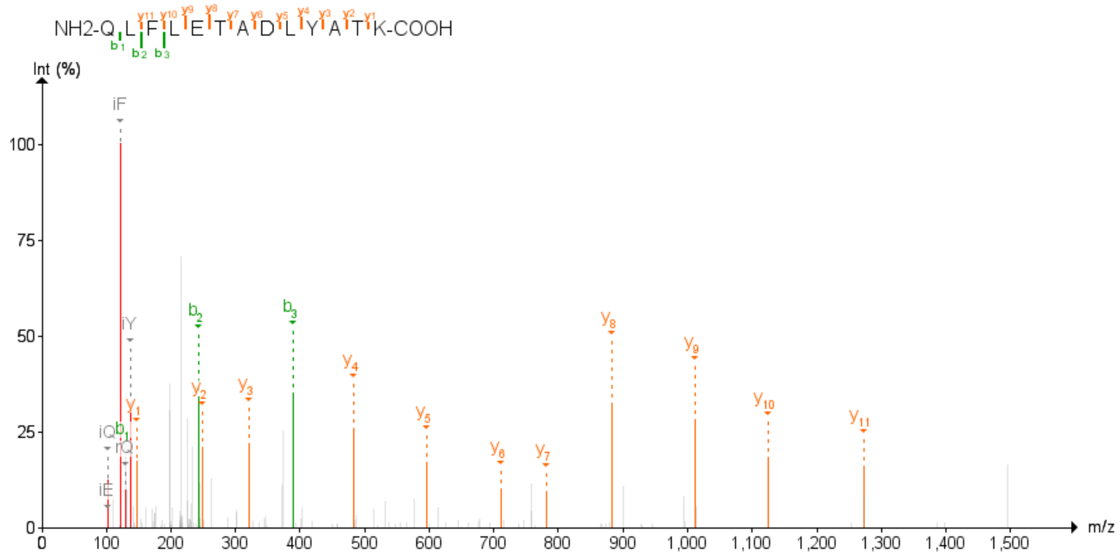

Figure 46: GPR89B;GPR89A Third experiment

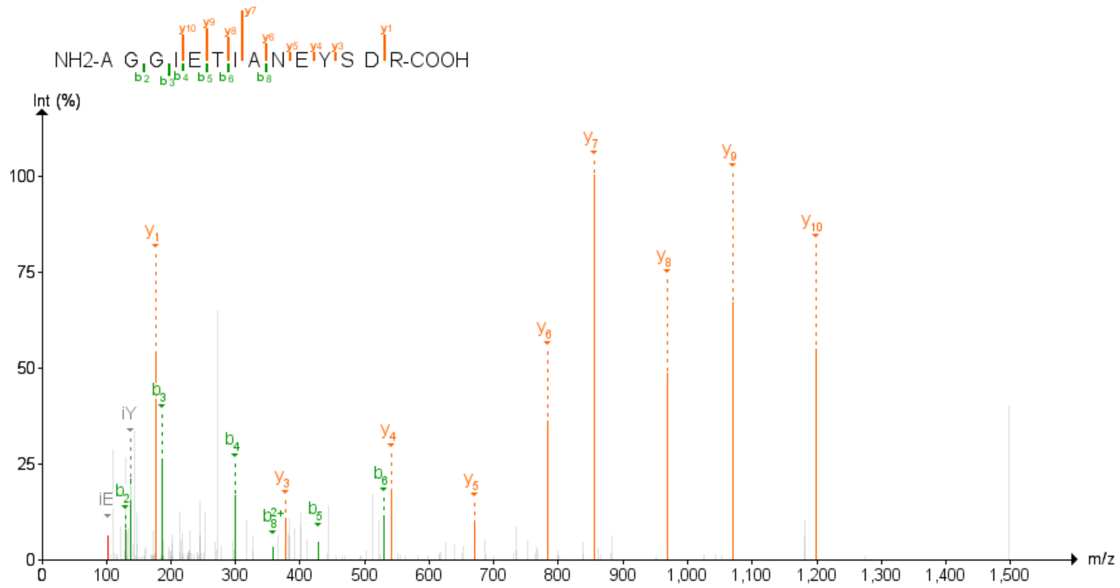

Figure 47: HSPA4 Second experiment

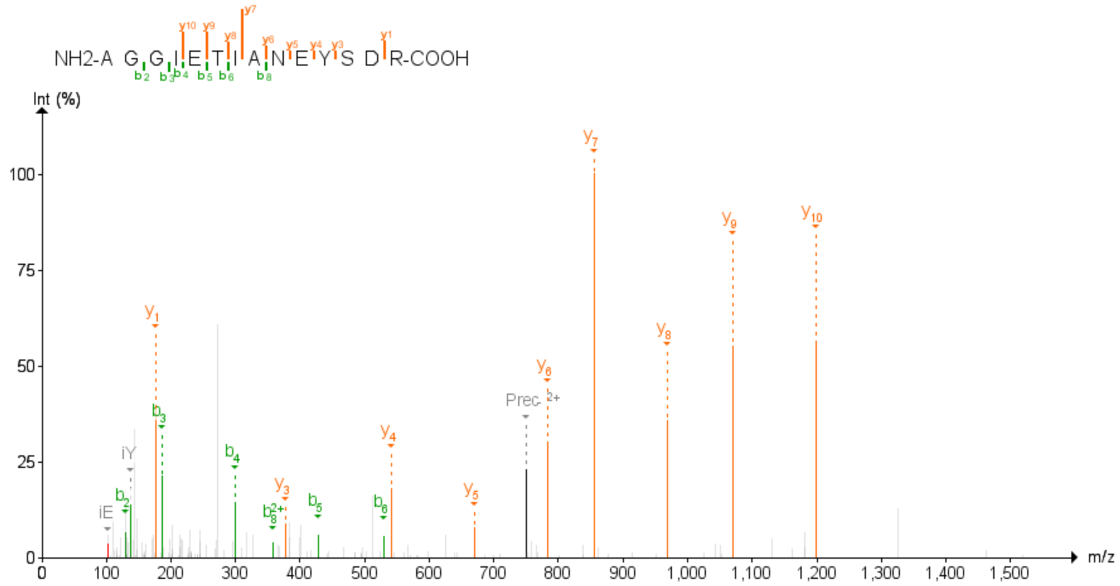

Figure 48: HSPA4 Third experiment

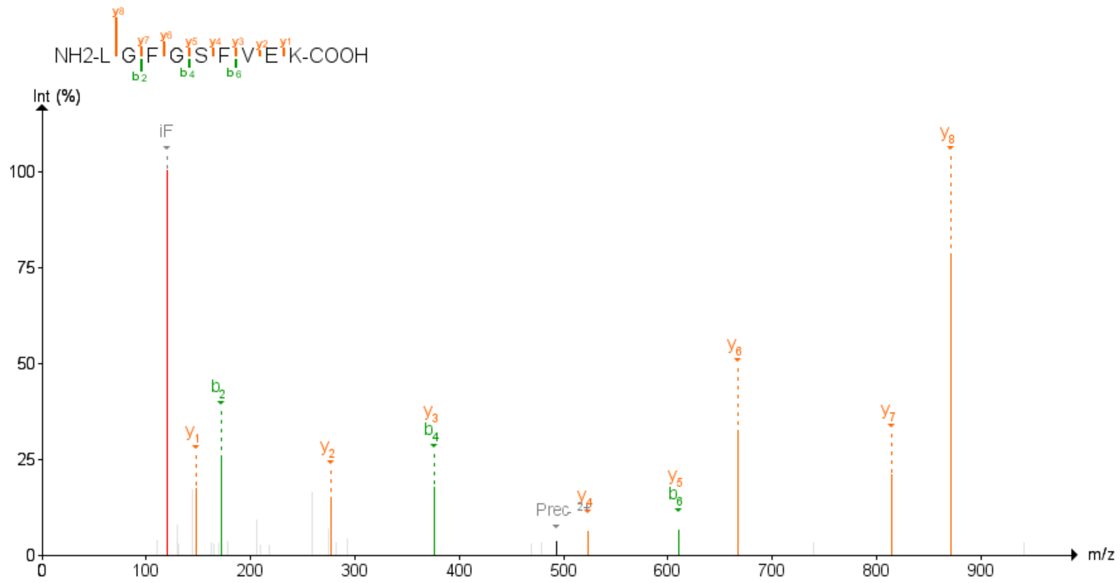

Figure 49: ITGB6 First experiment

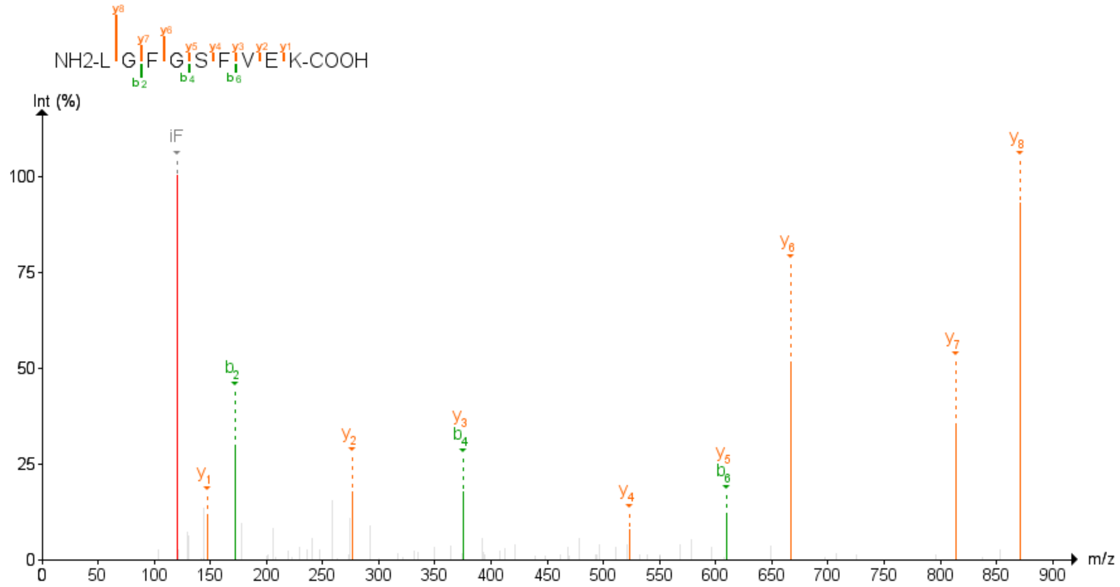

Figure 50: ITGB6 Second experiment

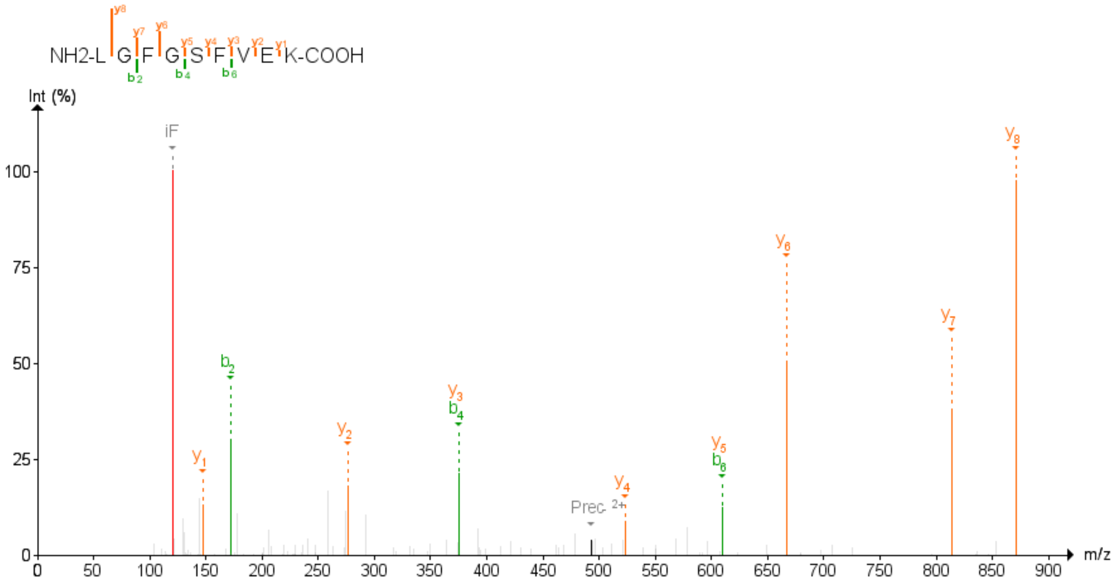

Figure 51: ITGB6 Third experiment

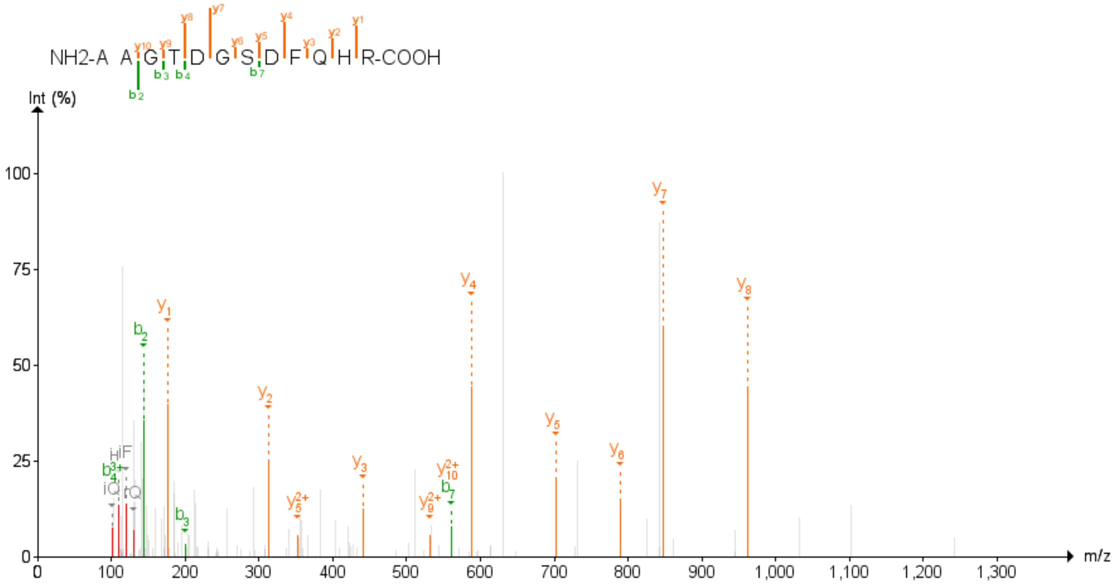

Figure 52: JAGN1 Second experiment

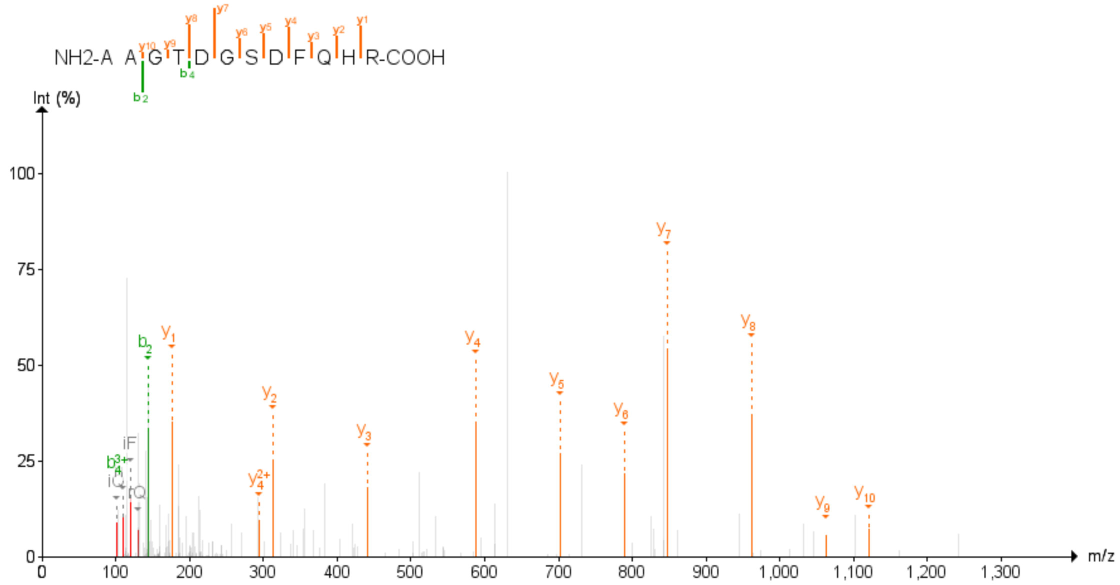

Figure 53: JAGN1 Third experiment

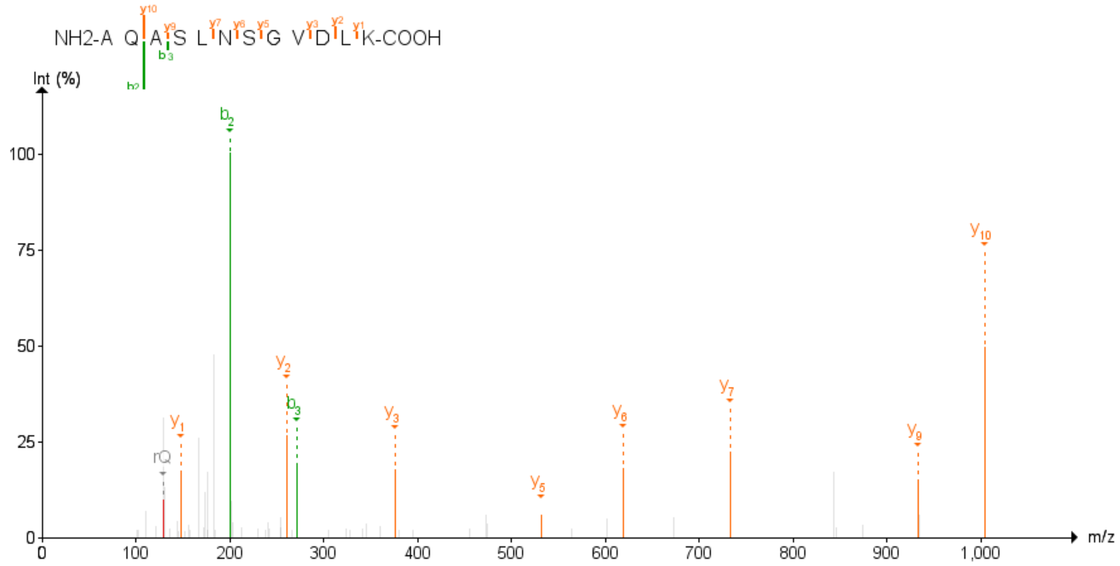

Figure 54: MOB3B First experiment

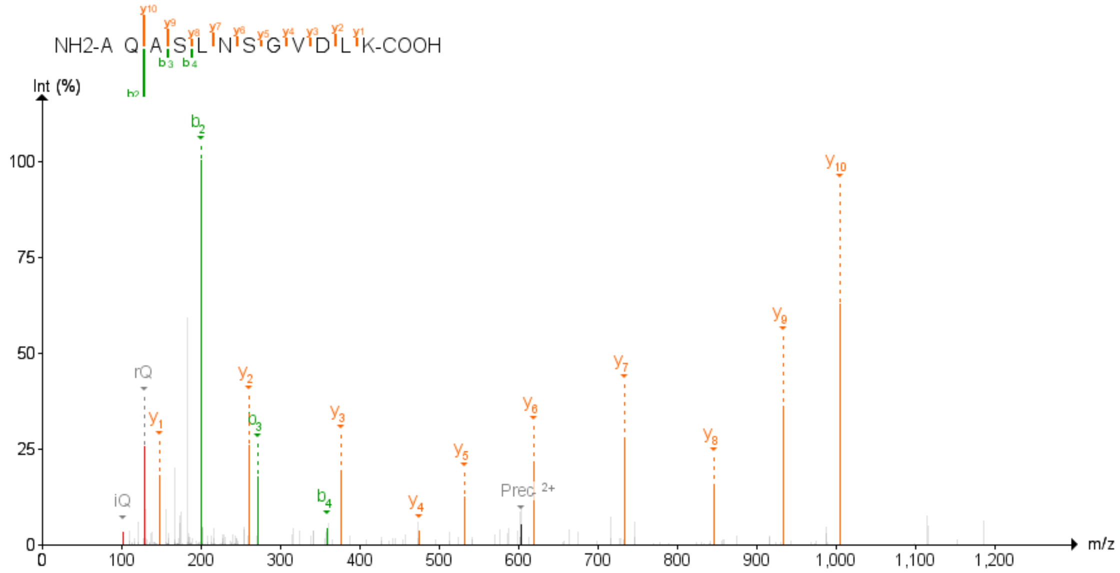

Figure 55: MOB3B Second experiment

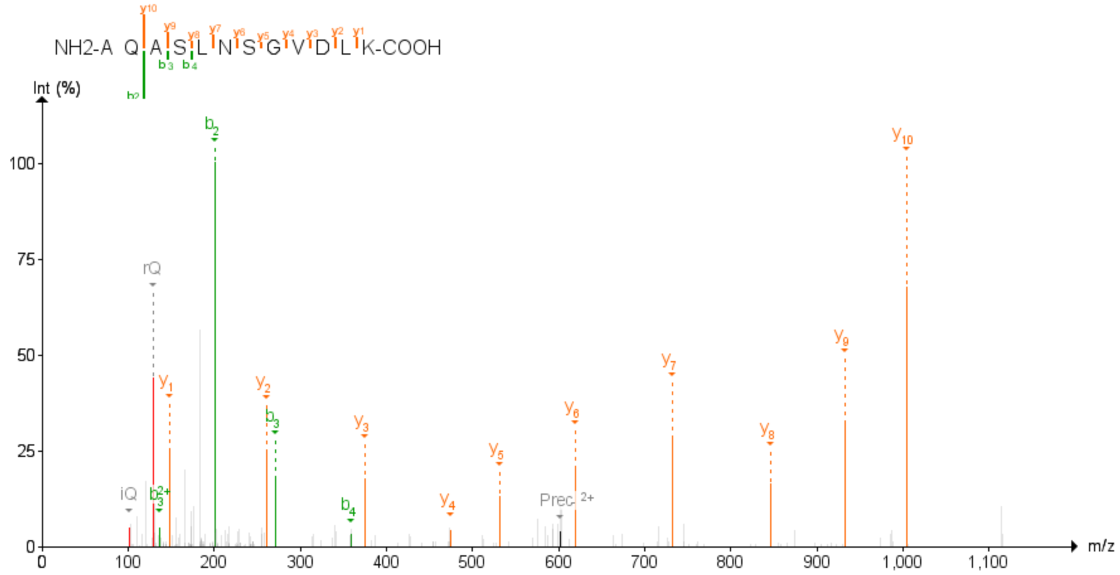

Figure 56: MOB3B Third experiment

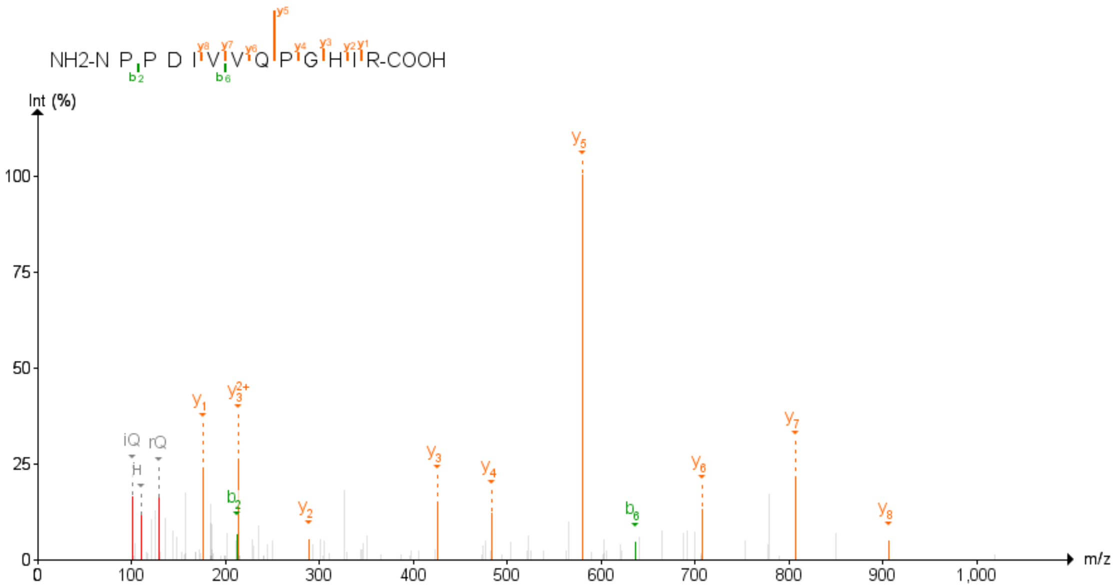

Figure 57: MPZL1 First experiment

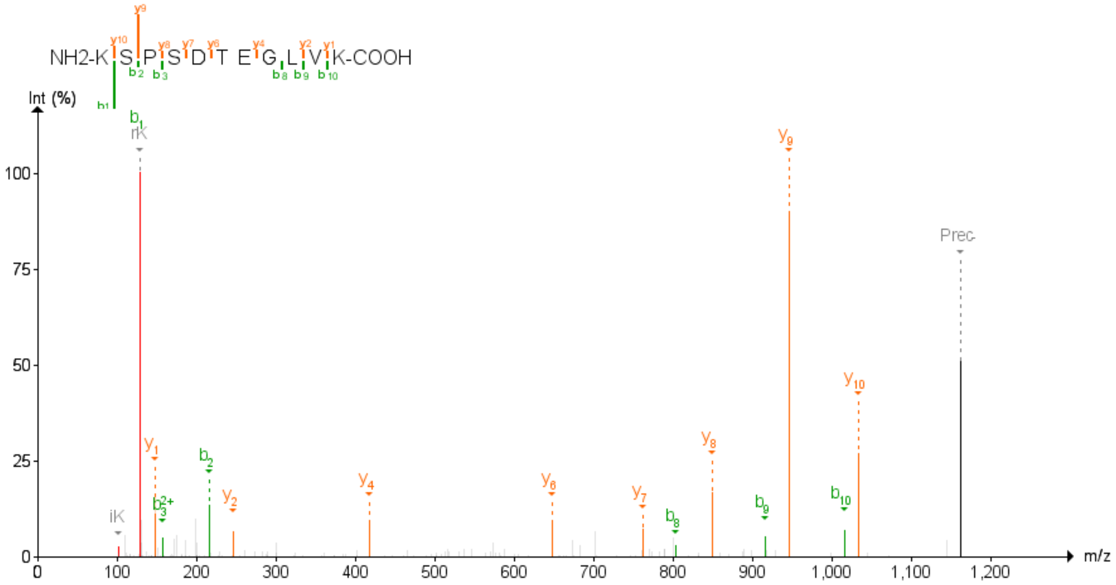

Figure 58: MPZL1 Second experiment

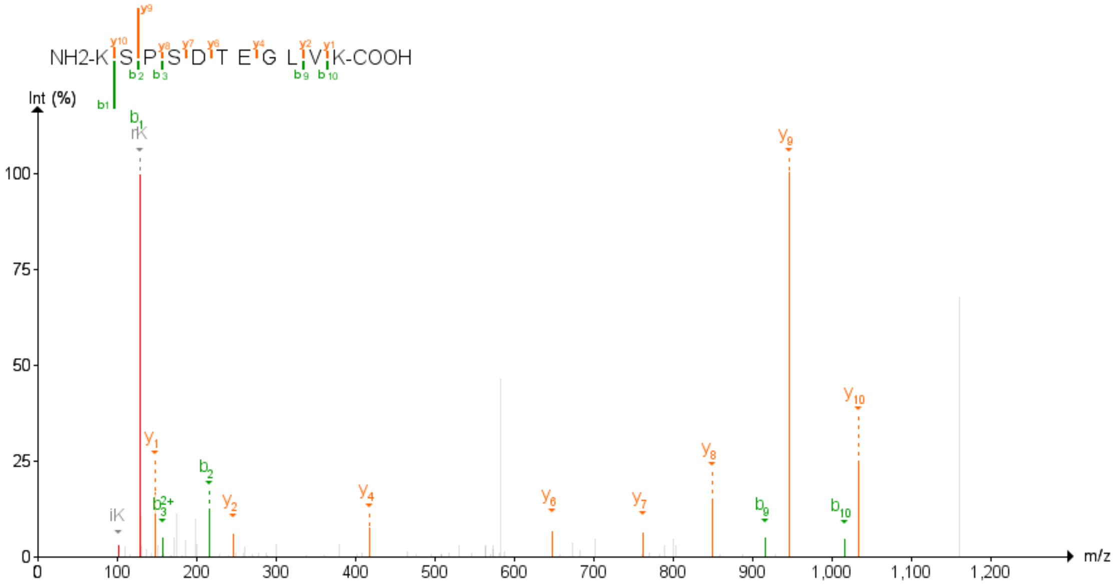

Figure 59: MPZL1 Third experiment

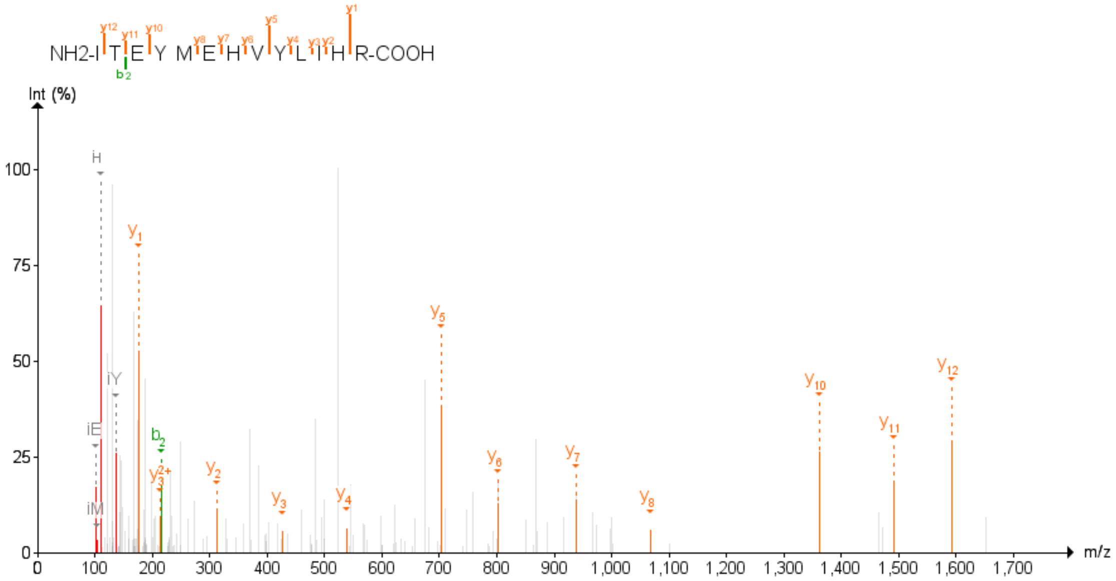

Figure 60: OSBPL2 First experiment

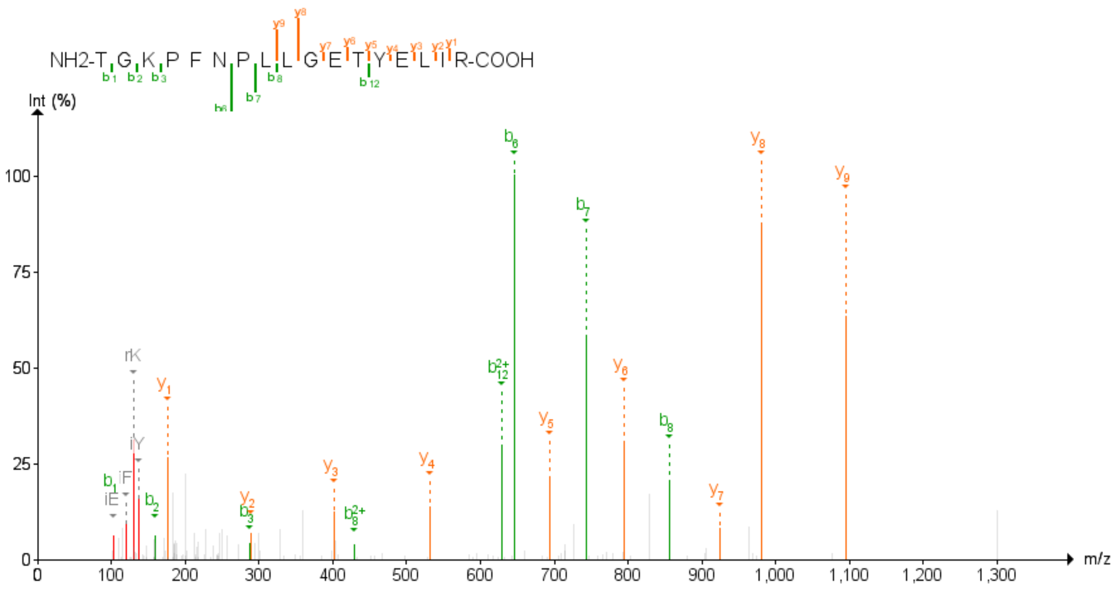

Figure 61: OSBPL2 Second experiment

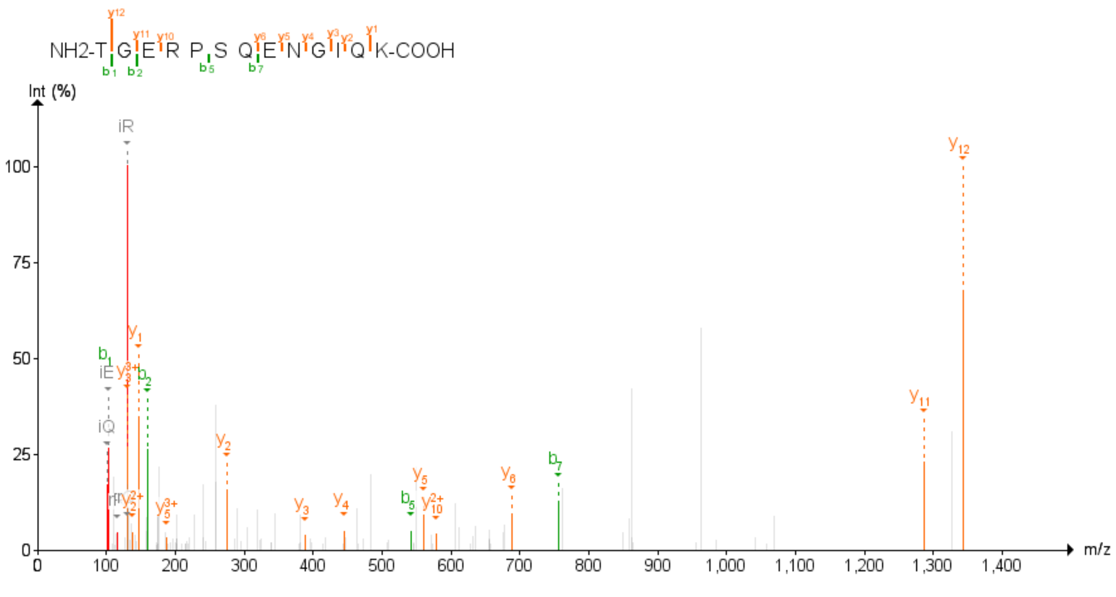

Figure 62: OSBPL2 Third experiment

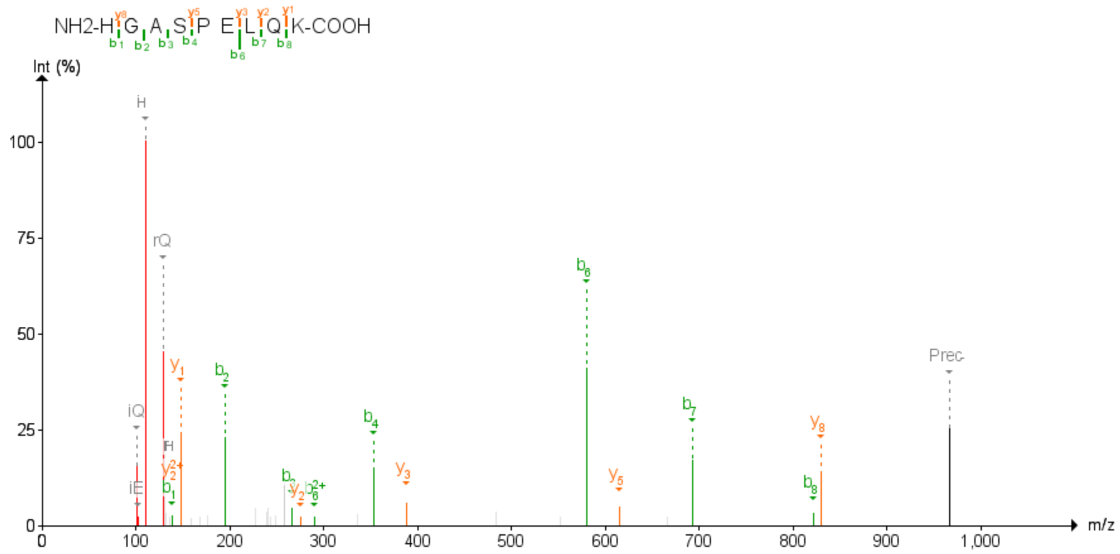

Figure 63: PEX11B First experiment

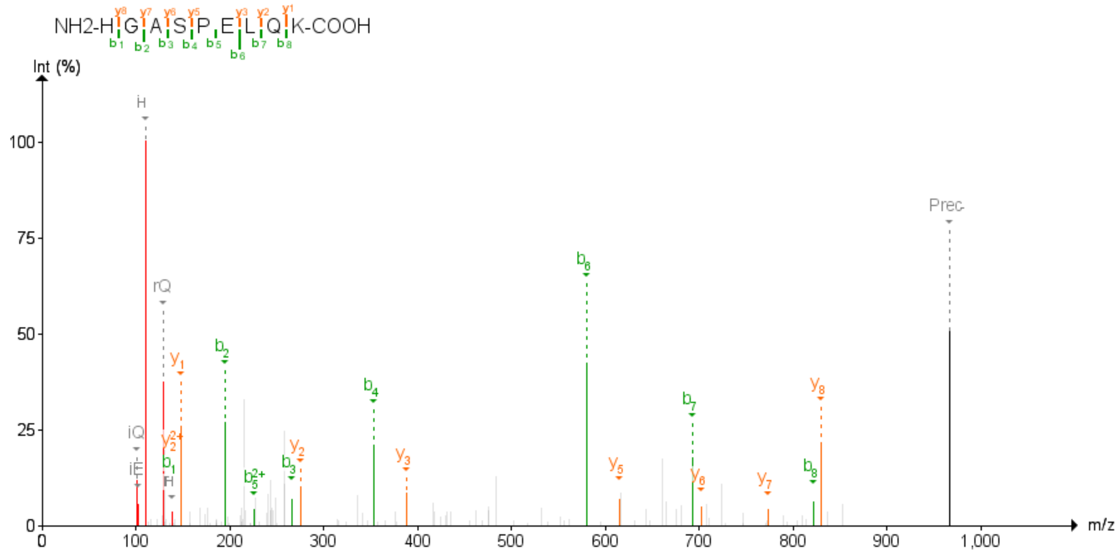

Figure 64: PEX11B Second experiment

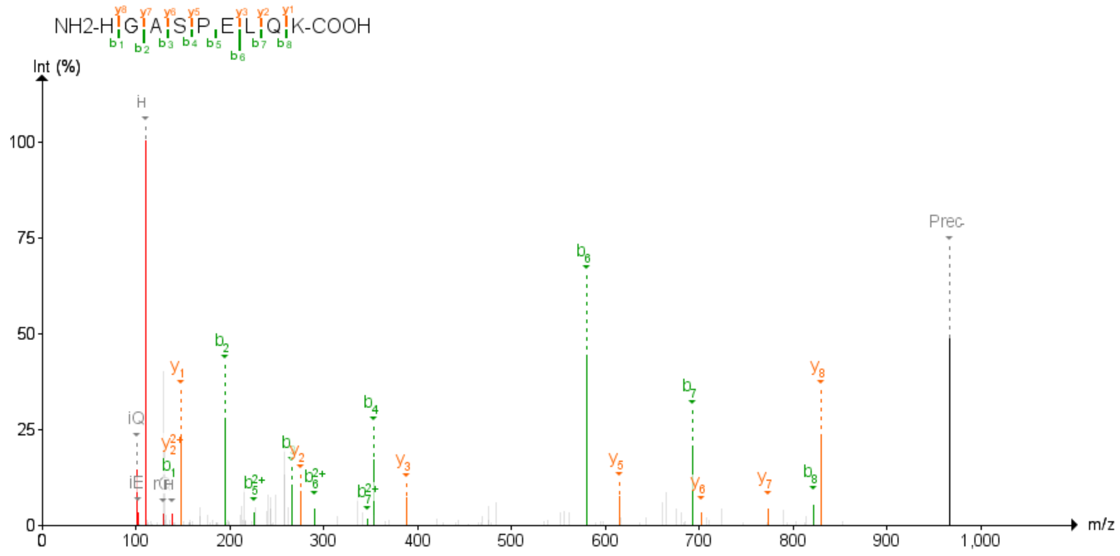

Figure 65: PEX11B Third experiment

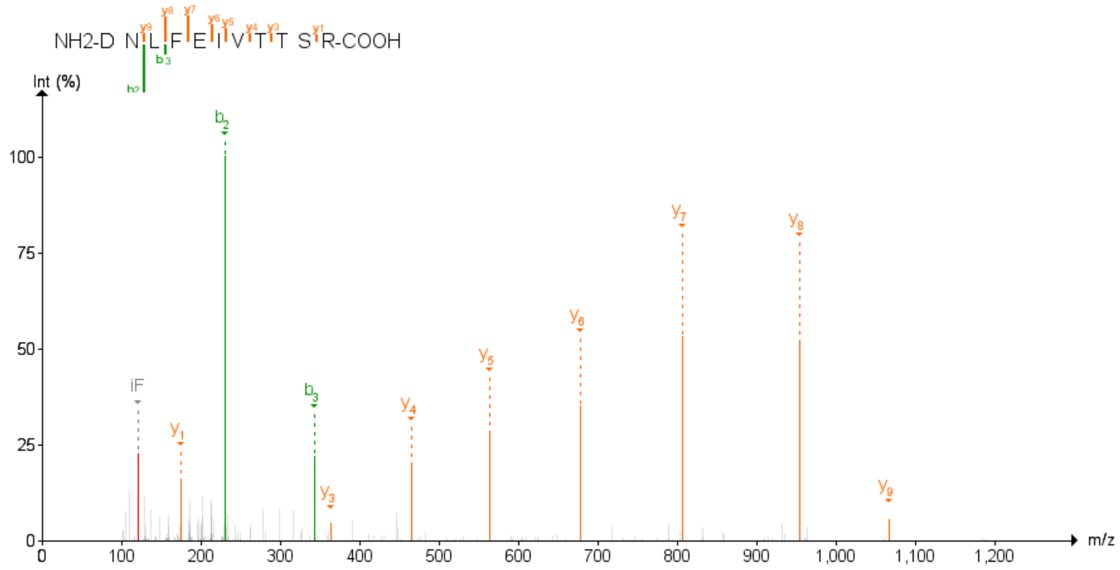

Figure 66: PLEKHA1 First experiment

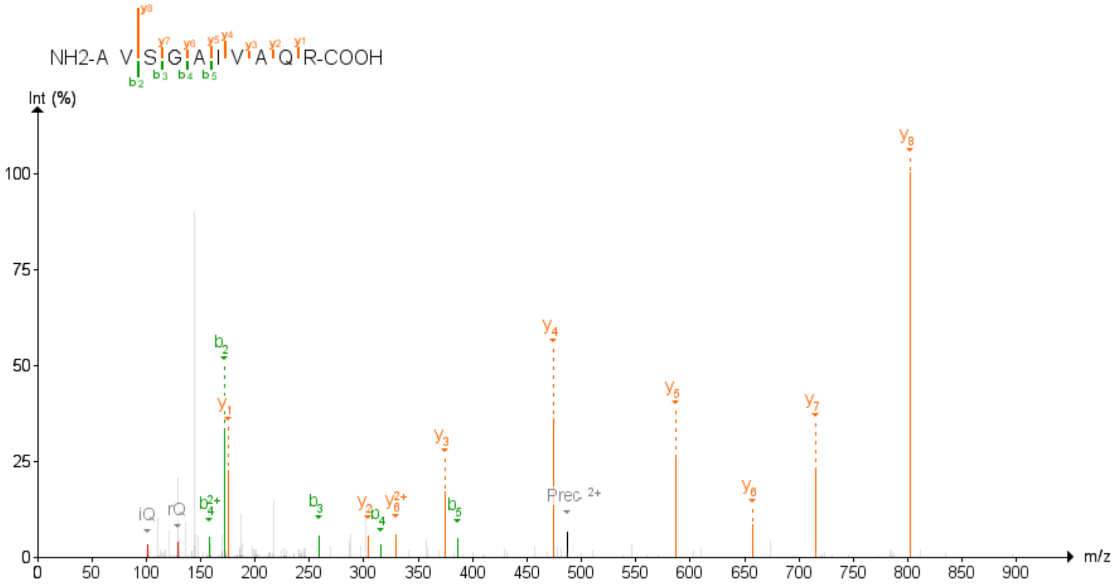

Figure 67: PLEKHA1 Second experiment

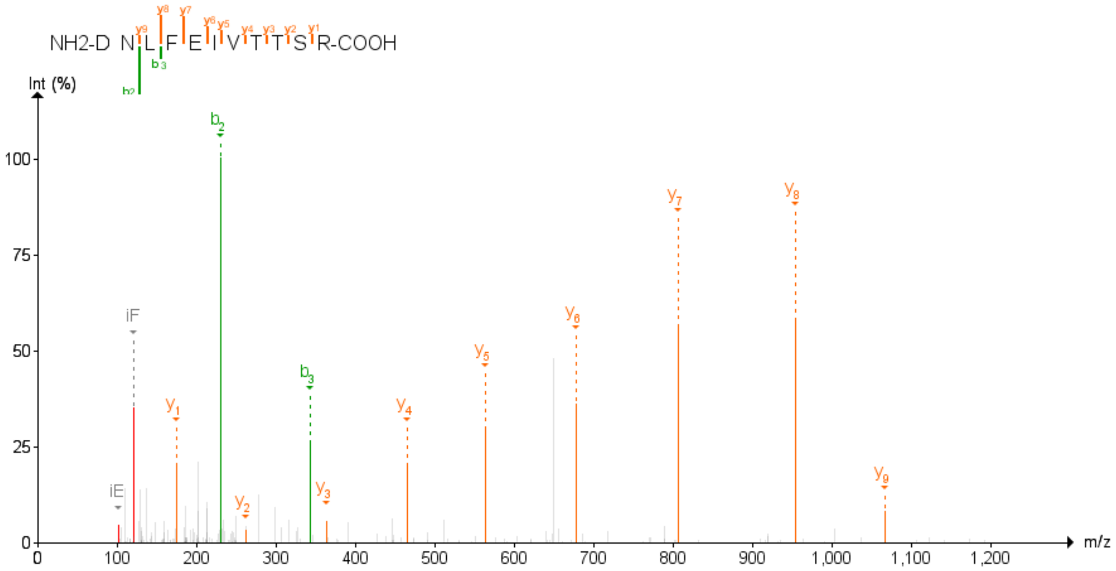

Figure 68: PLEKHA1 Third experiment

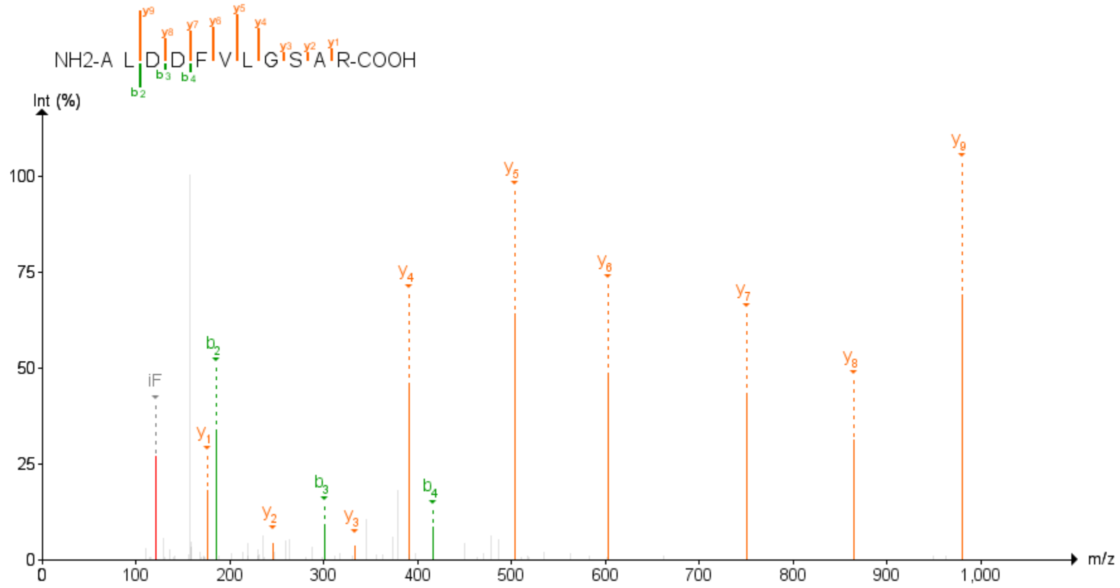

Figure 69: PRAF2 First experiment

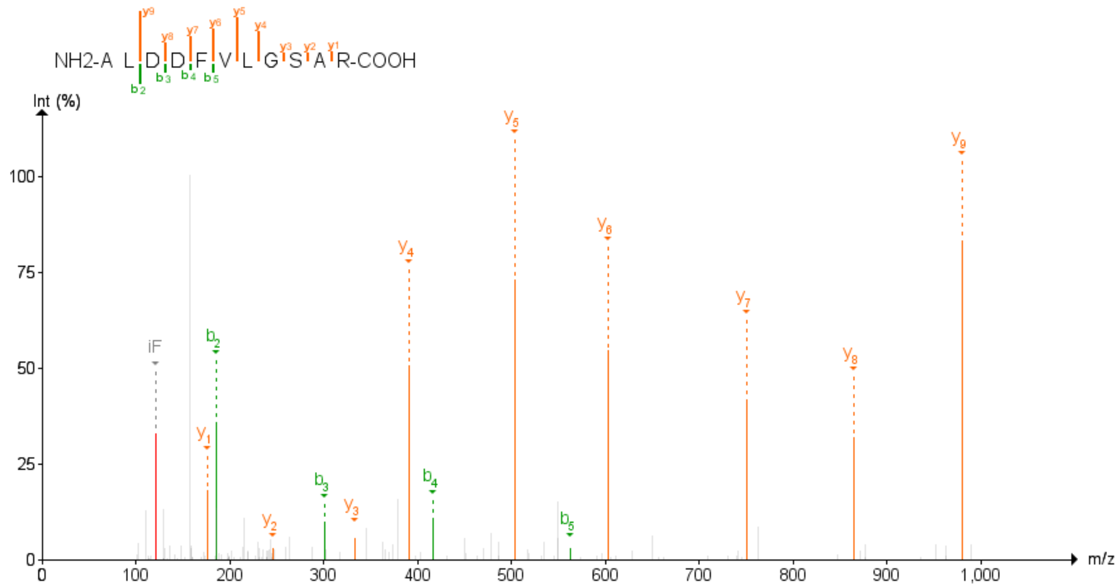

Figure 70: PRAF2 Second experiment

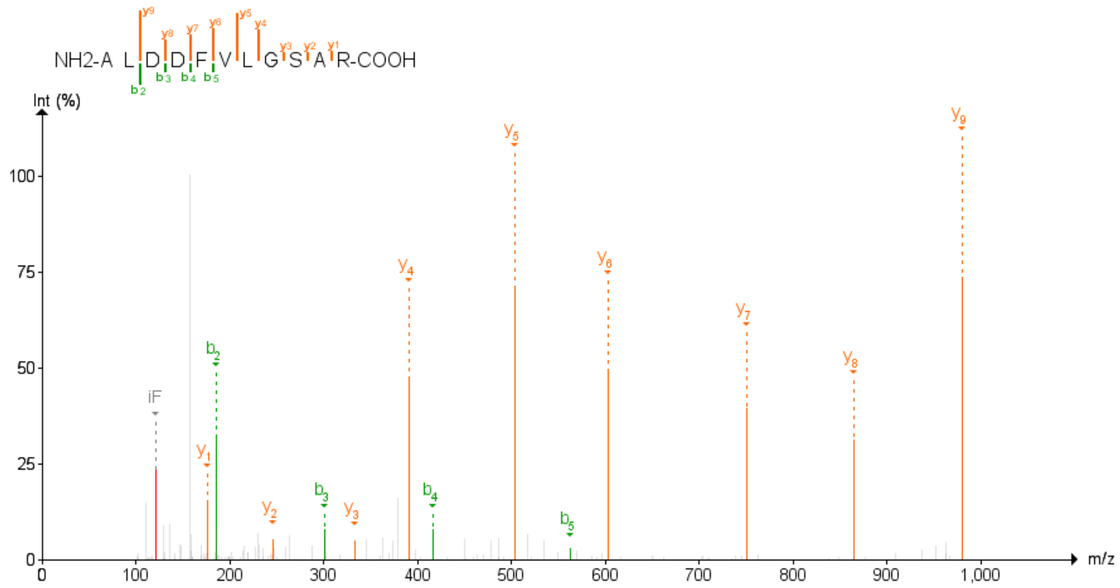

Figure 71: PRAF2 Third experiment

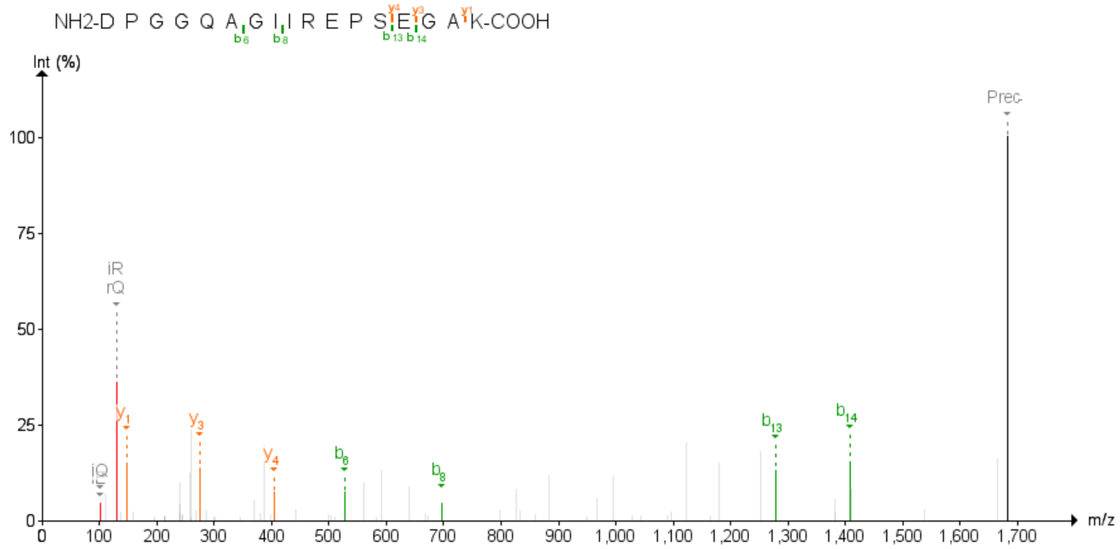

Figure 72: PRR14L First experiment

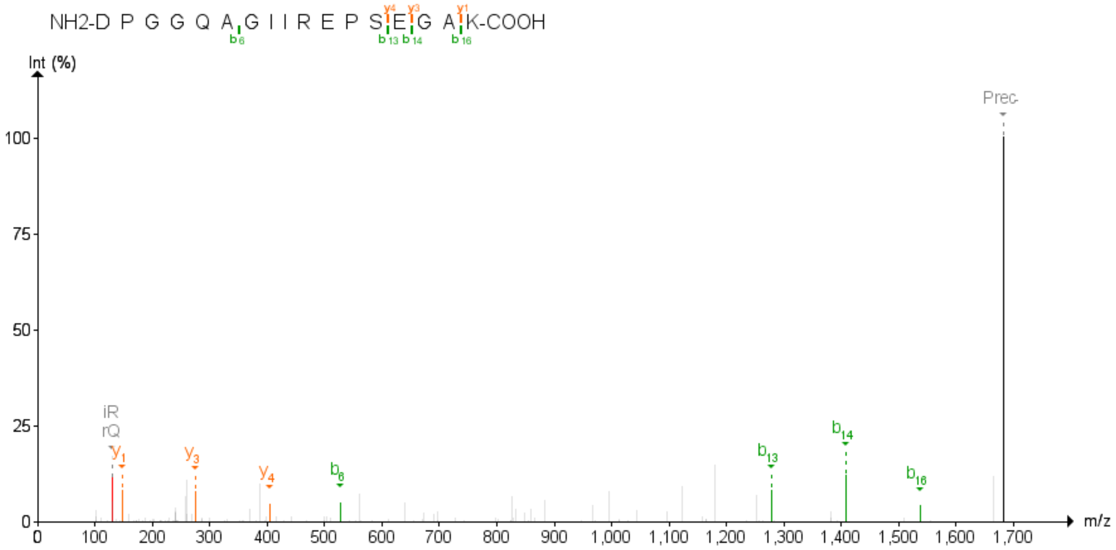

Figure 73: PRR14L First experiment

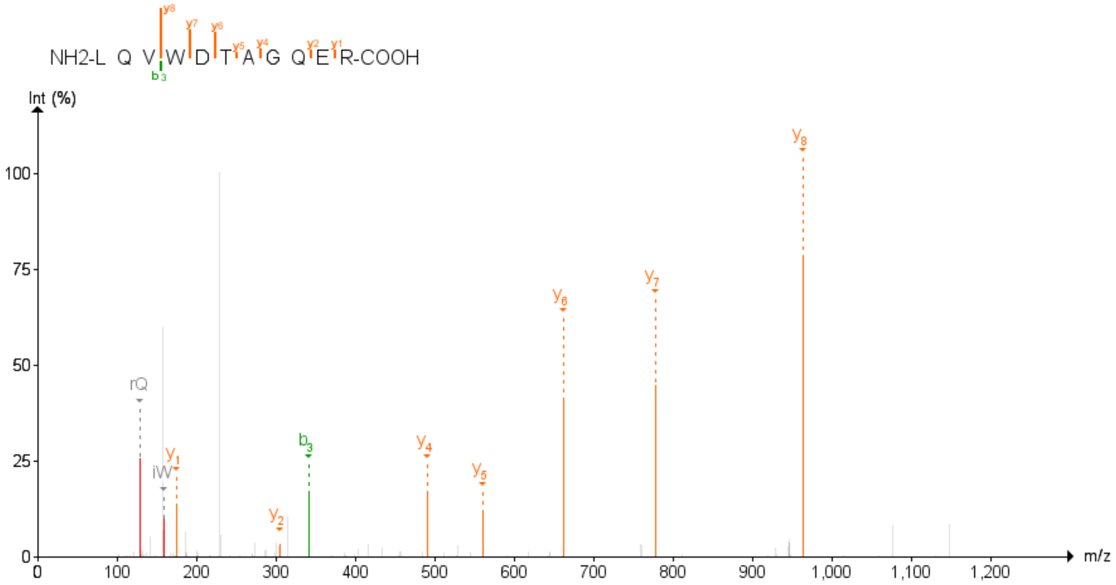

Figure 74: RAB13 Second experiment

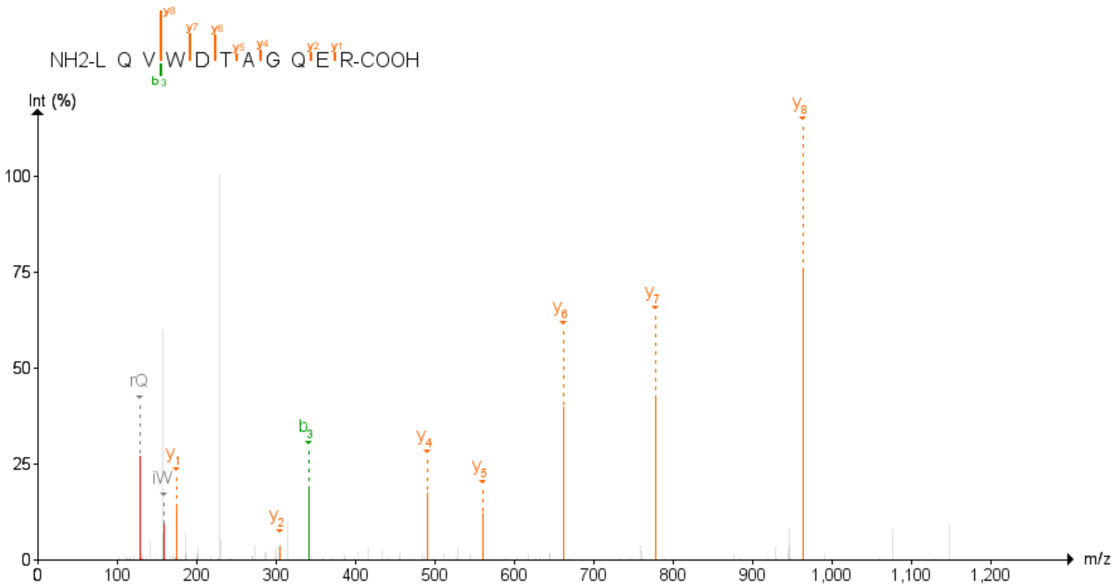

Figure 75: RAB13 Third experiment

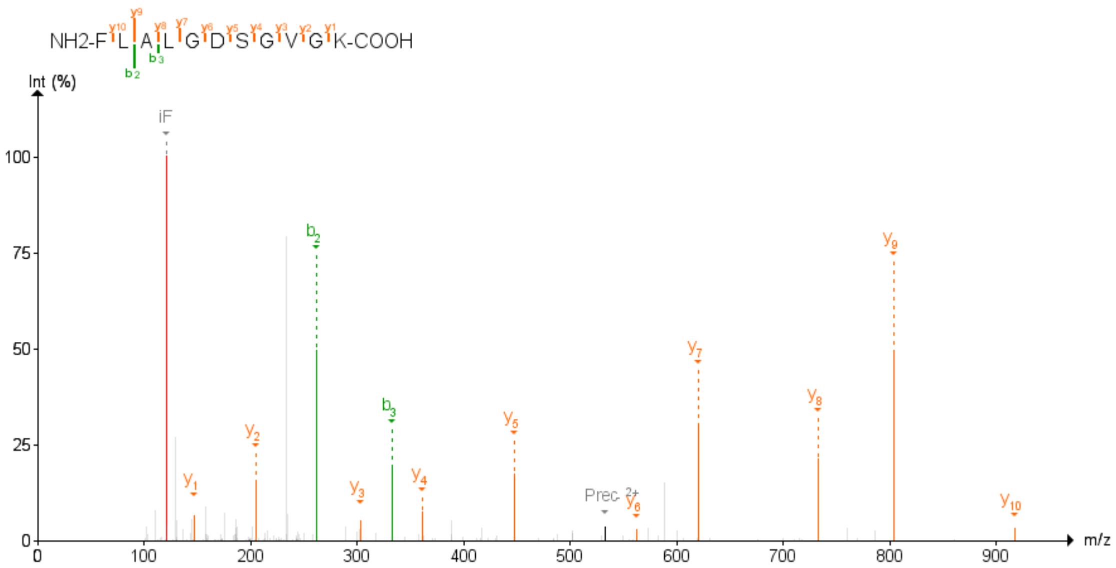

Figure 76: RAB27A Second experiment

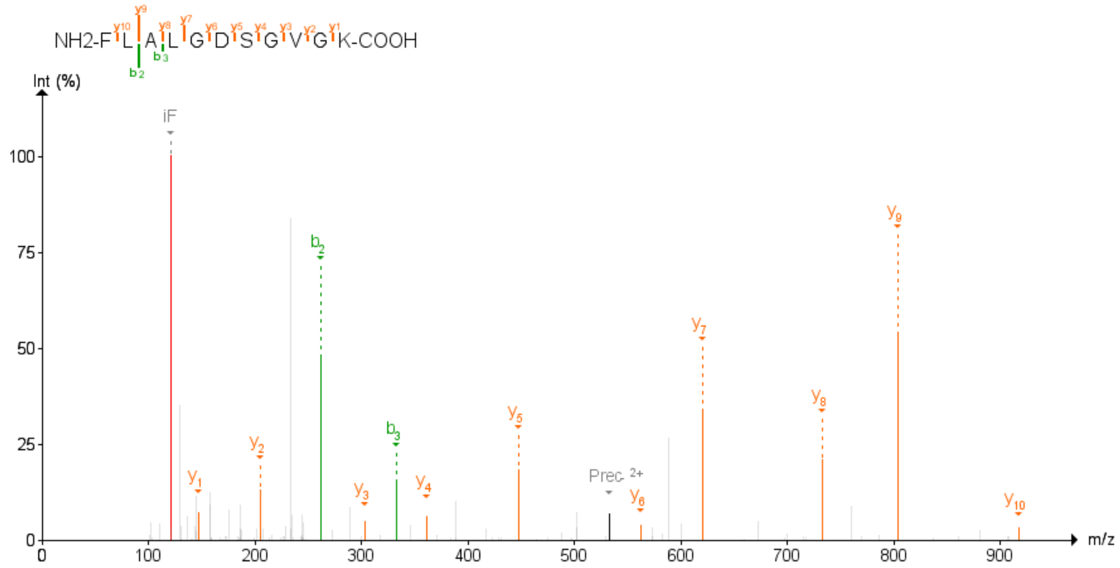

Figure 77: RAB27A Third experiment

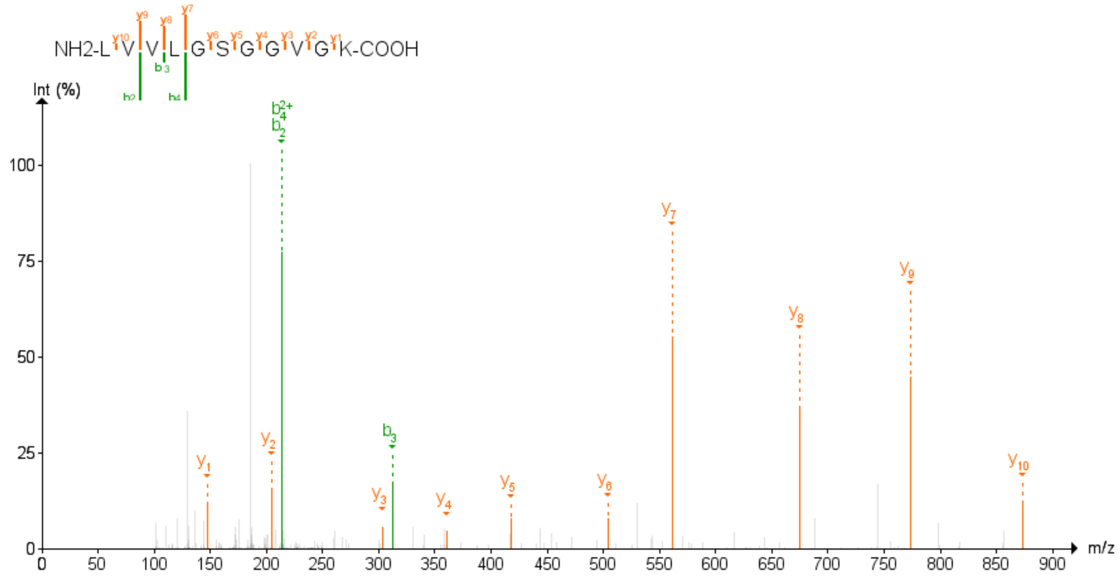

Figure 78: RAP1A;RAP1B Second experiment

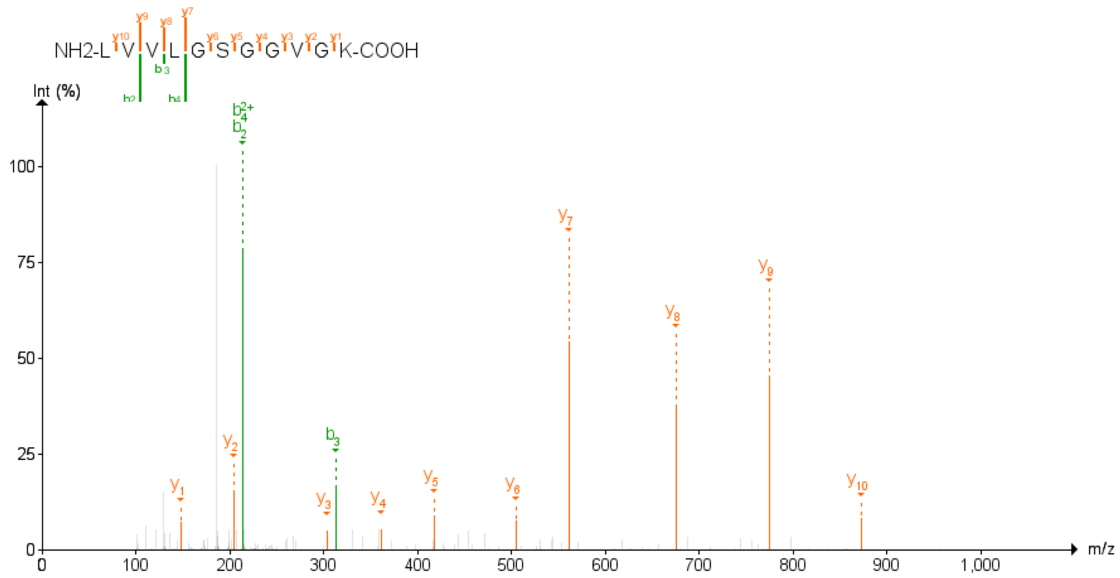

Figure 79: RAP1A;RAP1B Third experiment

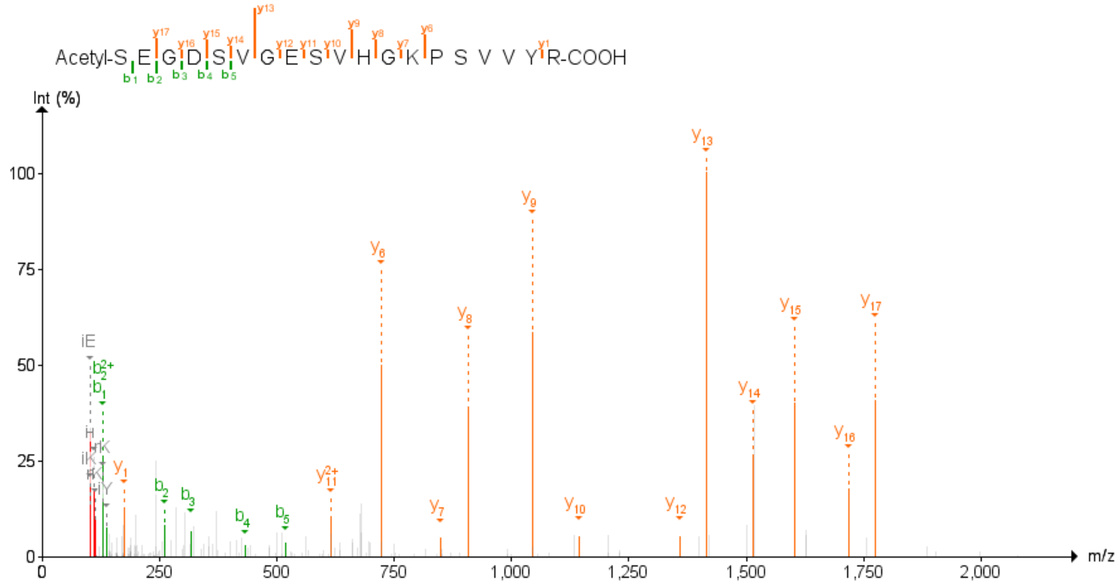

Figure 80: RER1 Second experiment

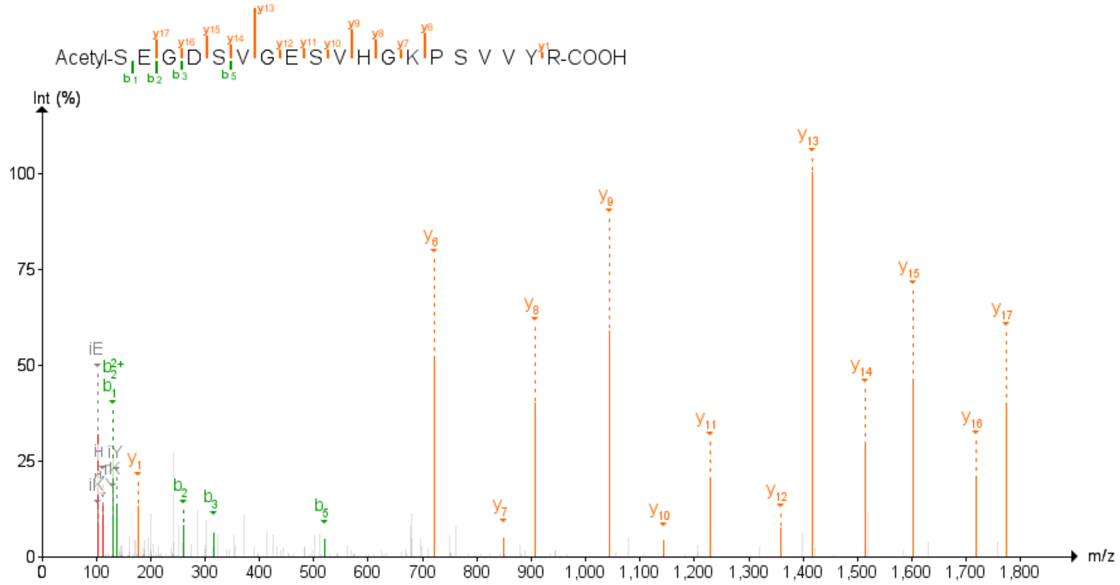

Figure 81: RER1 Third experiment

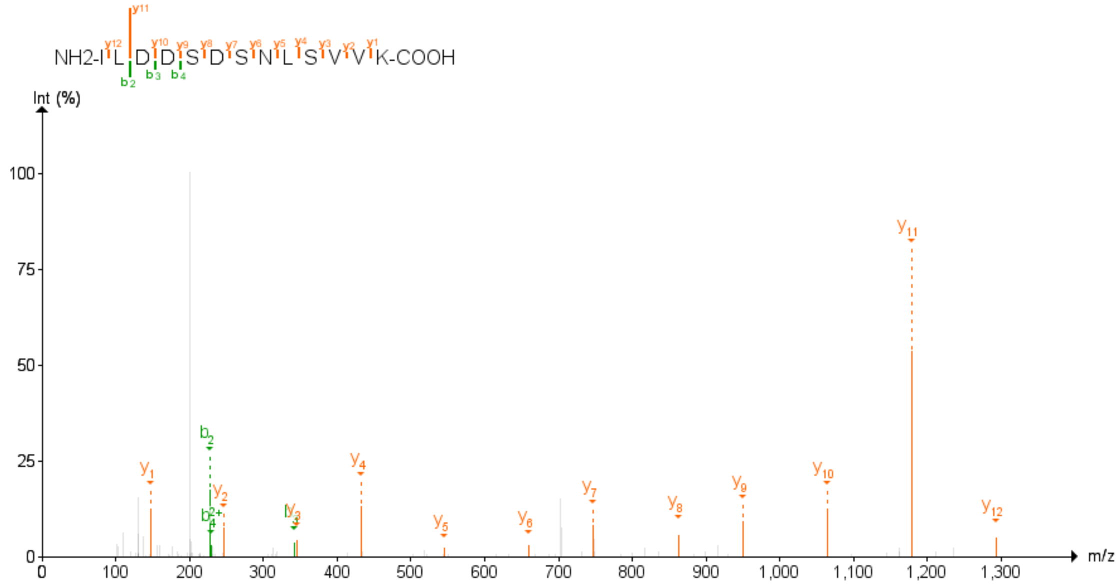

Figure 82: RGPD3;RGPD4 Second experiment

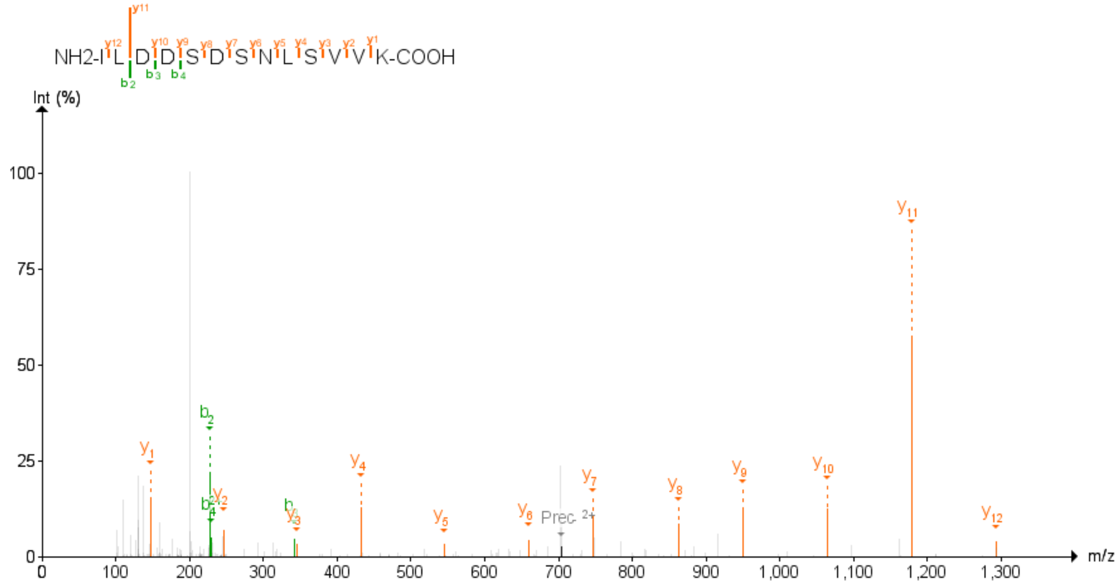

Figure 83: RGPD3;RGPD4 Third experiment

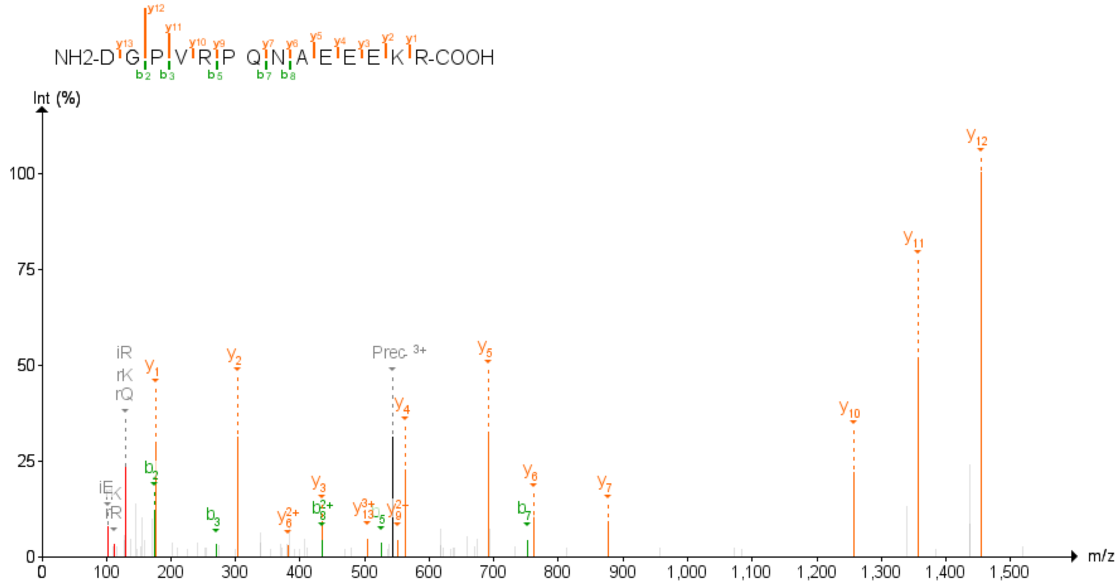

Figure 84: SLC39A7 First experiment

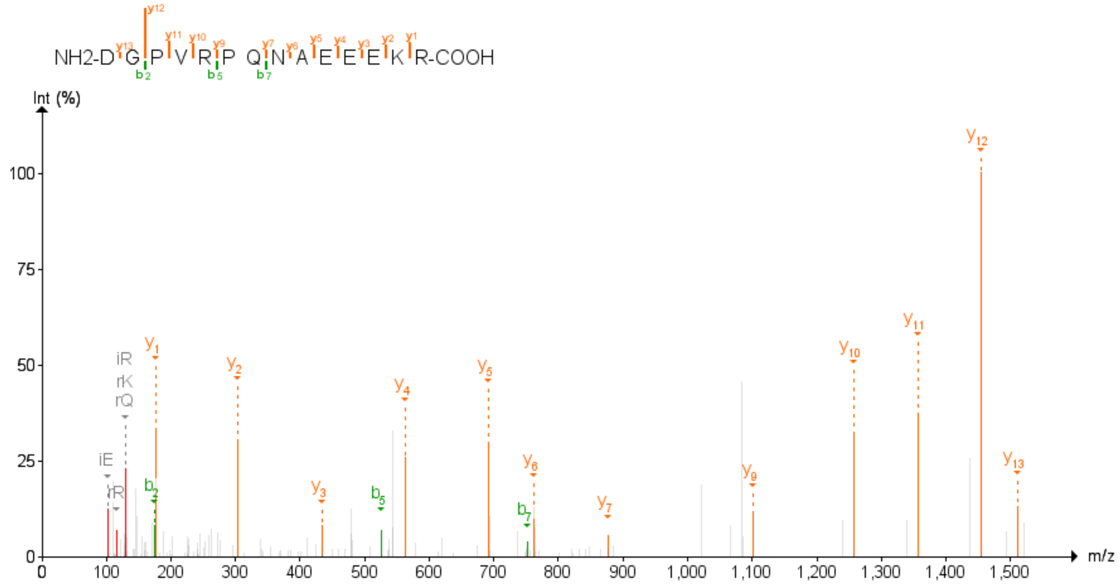

Figure 85: SLC39A7 Second experiment

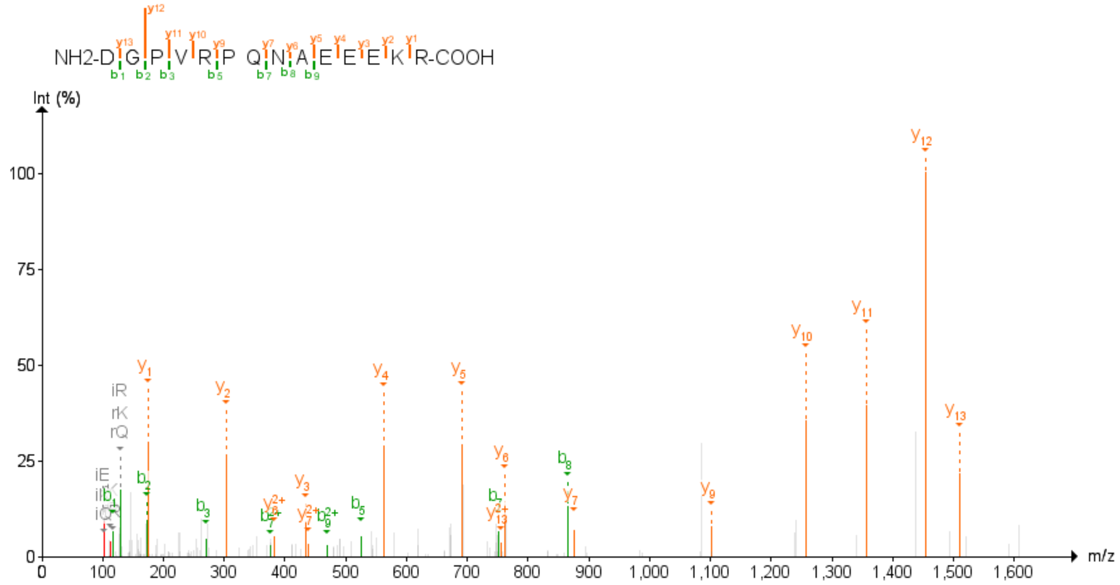

Figure 86: SLC39A7 Third experiment

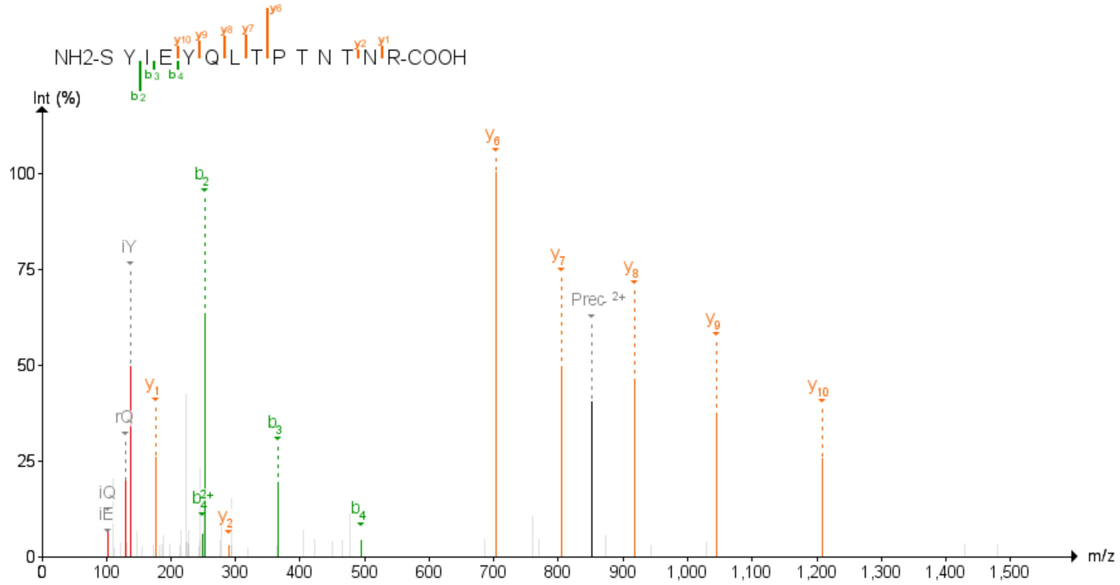

Figure 87: SNX9 First experiment

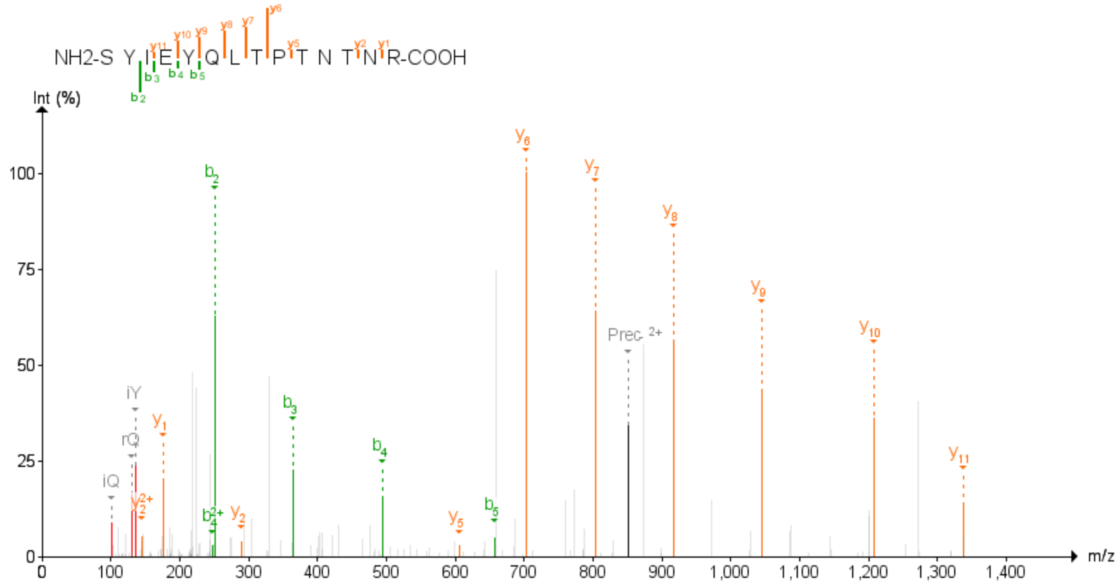

Figure 88: SNX9 Third experiment

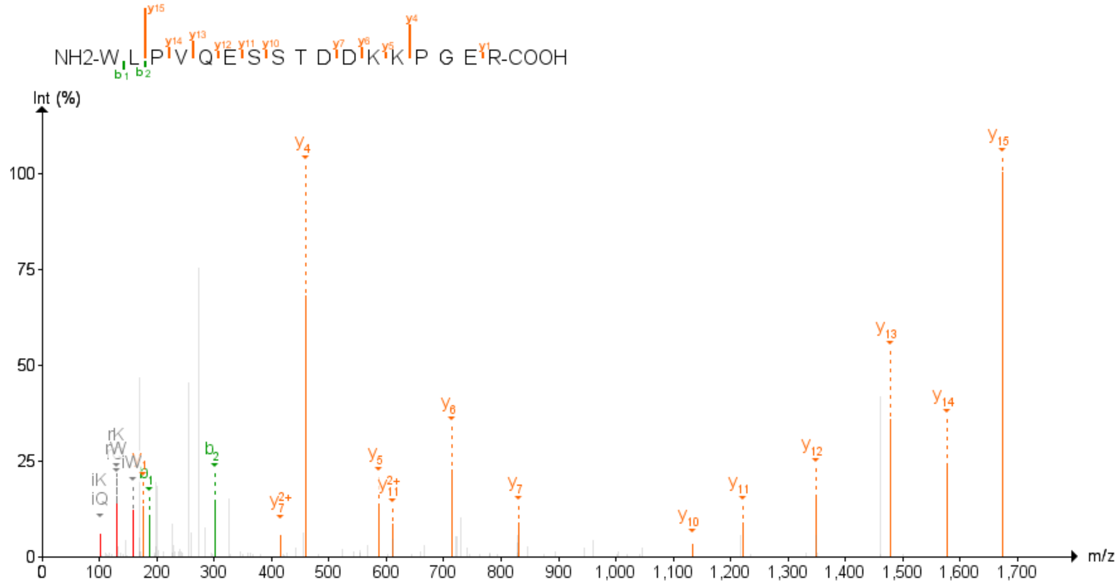

Figure 89: SPCS1 First experiment

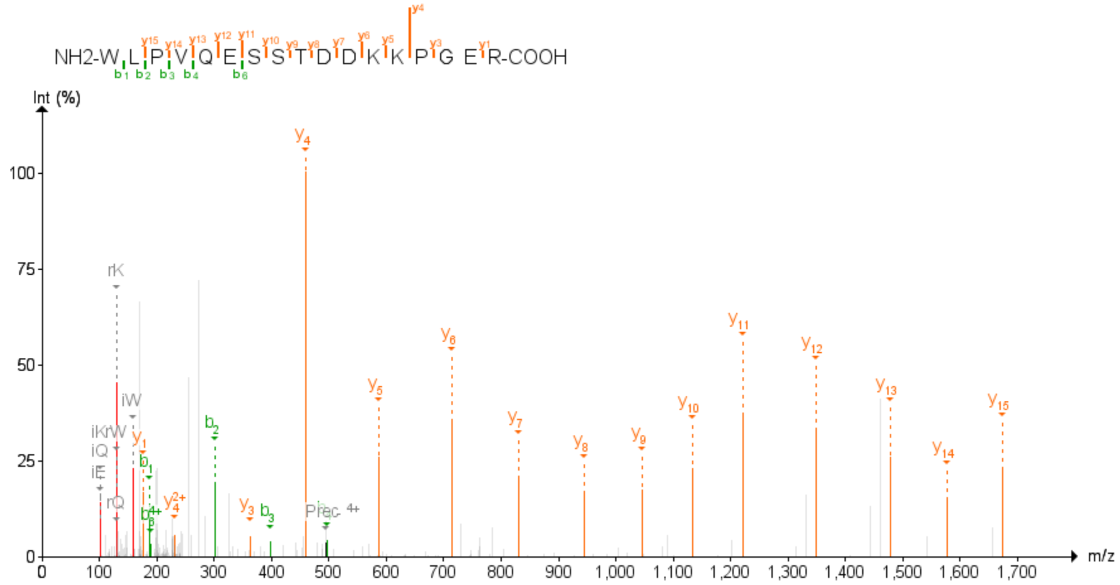

Figure 90: SPCS1 Second experiment

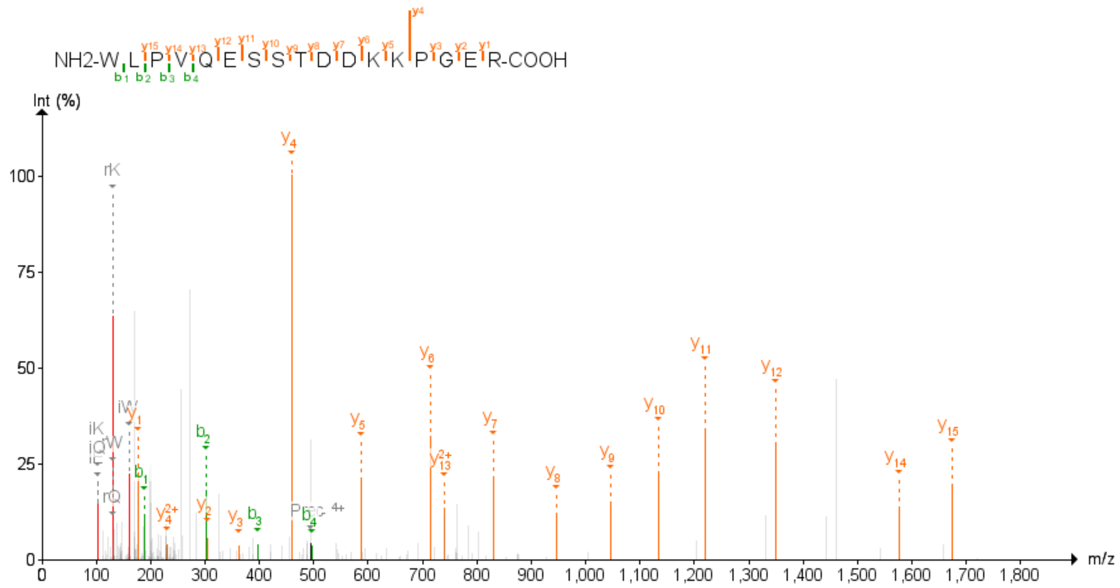

Figure 91: SPCS1 Third experiment

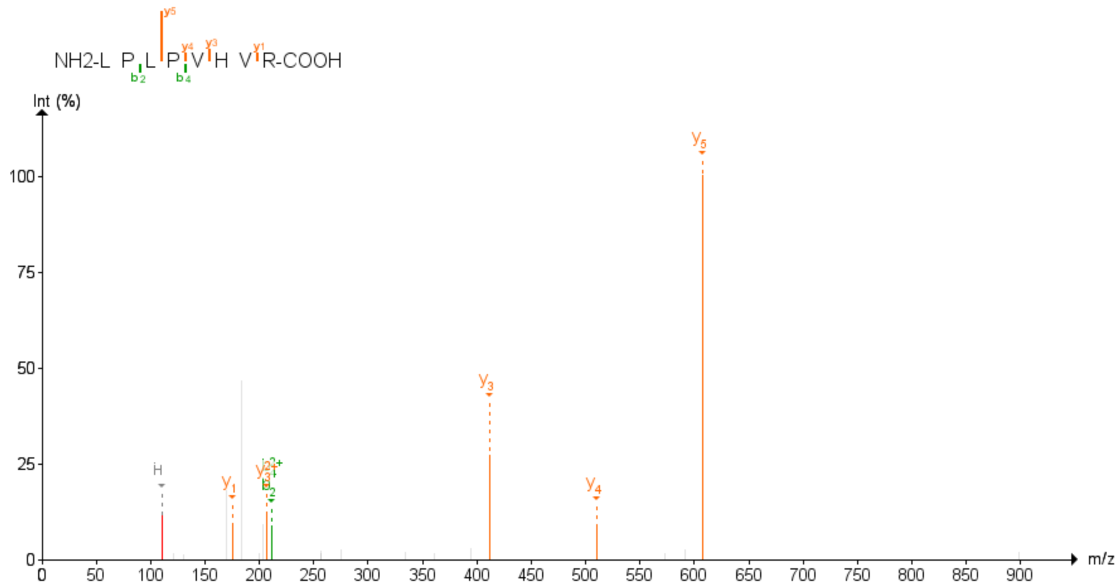

Figure 92: TLCD1 First experiment

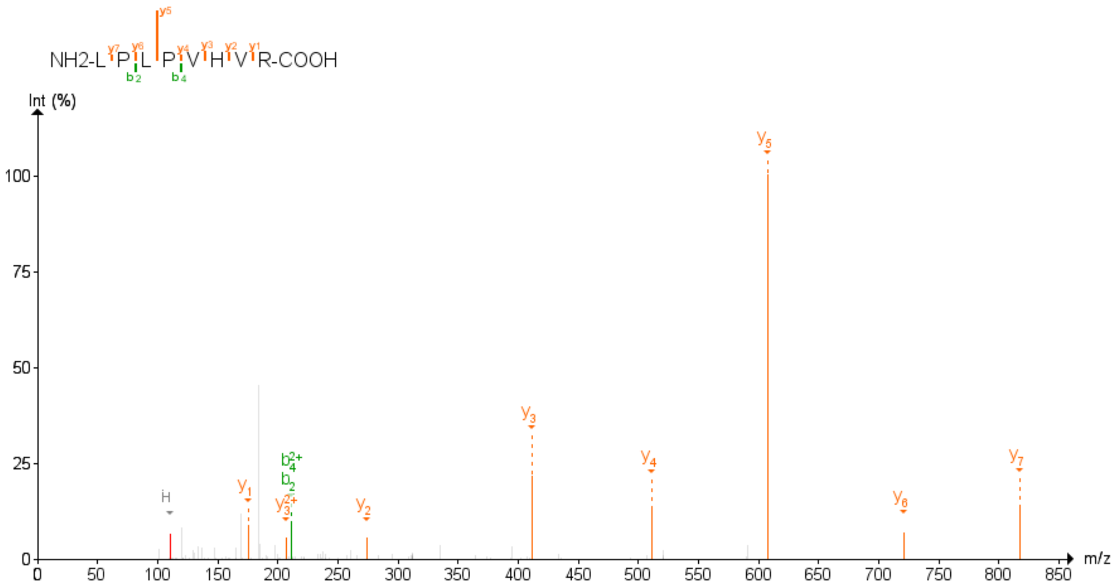

Figure 93: TLCD1 Third experiment

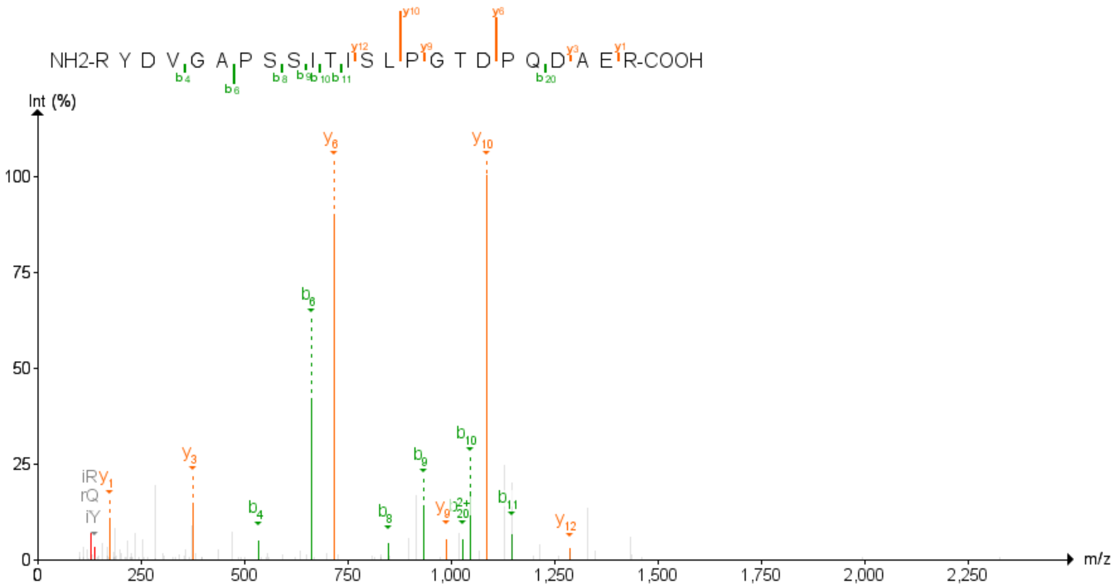

Figure 94: TMEM115 First experiment

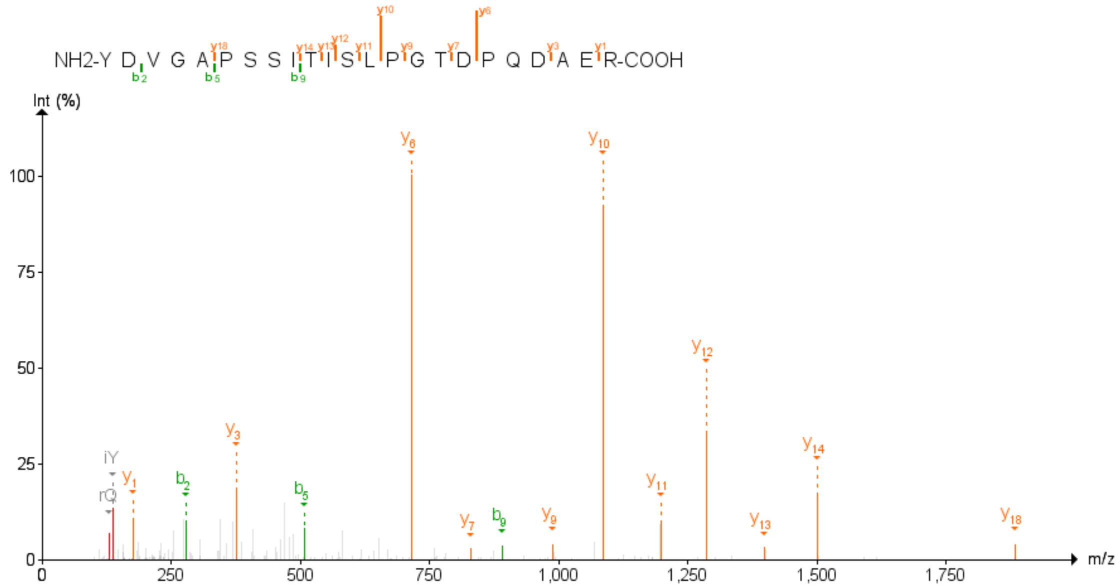

Figure 95: TMEM115 Second experiment

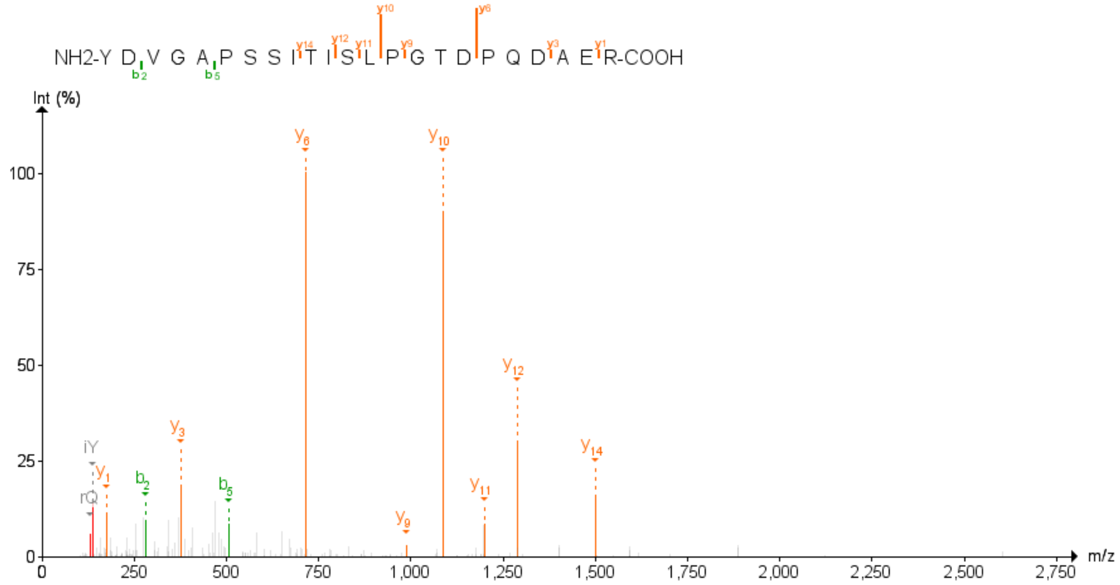

Figure 96: TMEM115 Third experiment

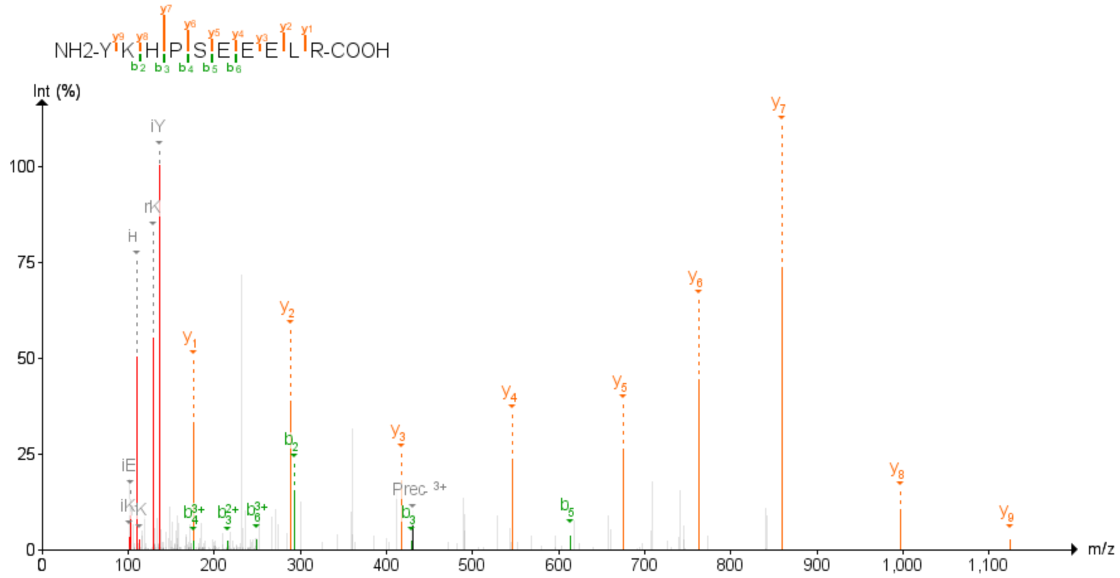

Figure 97: TMEM161A Second experiment

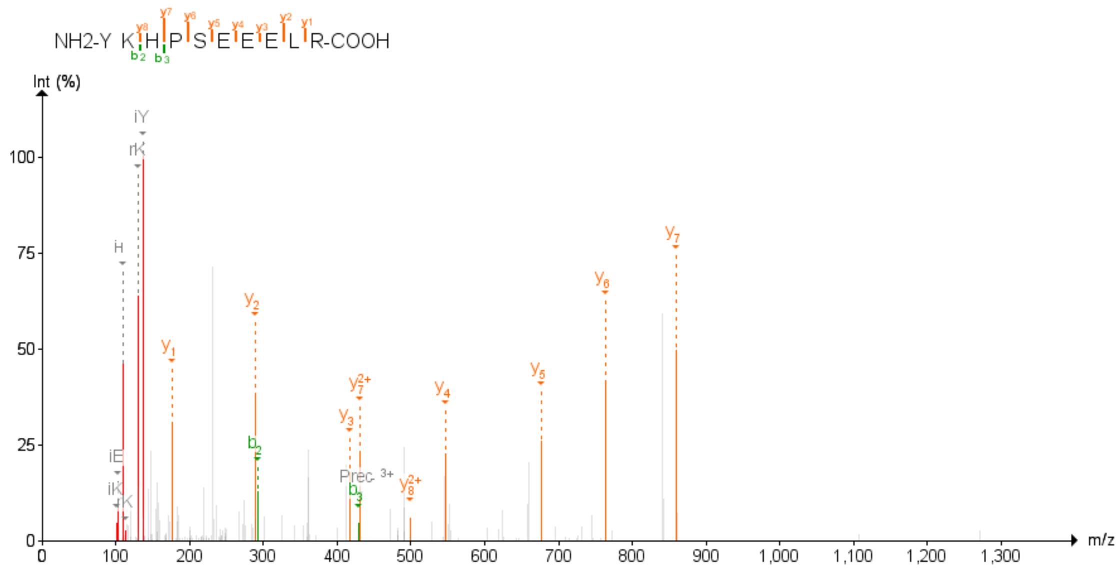

Figure 98: TMEM161A Third experiment

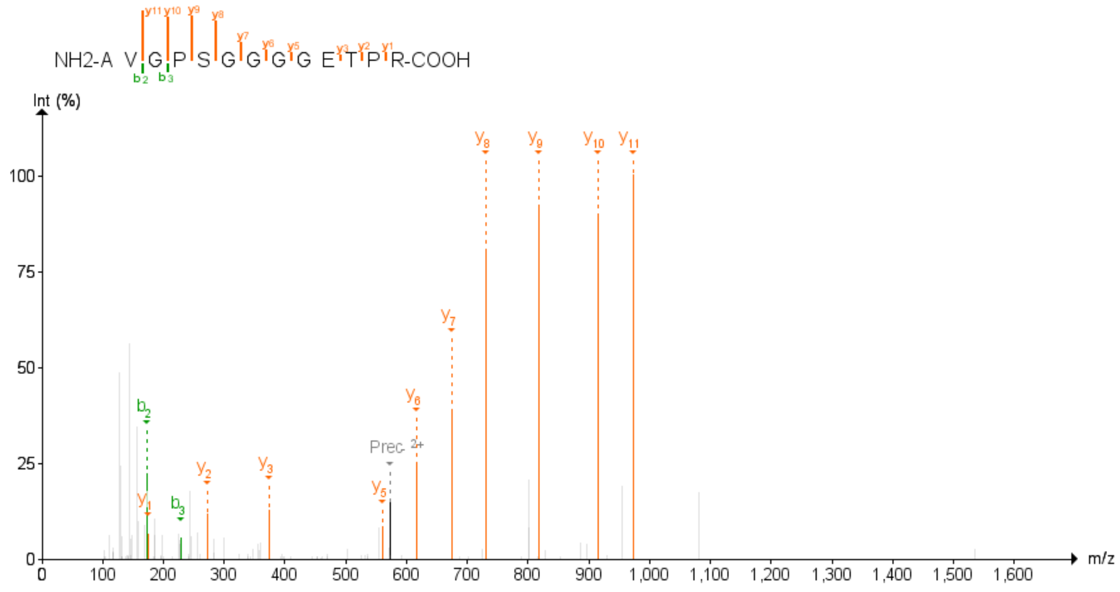

Figure 99: TMEM245 Second experiment

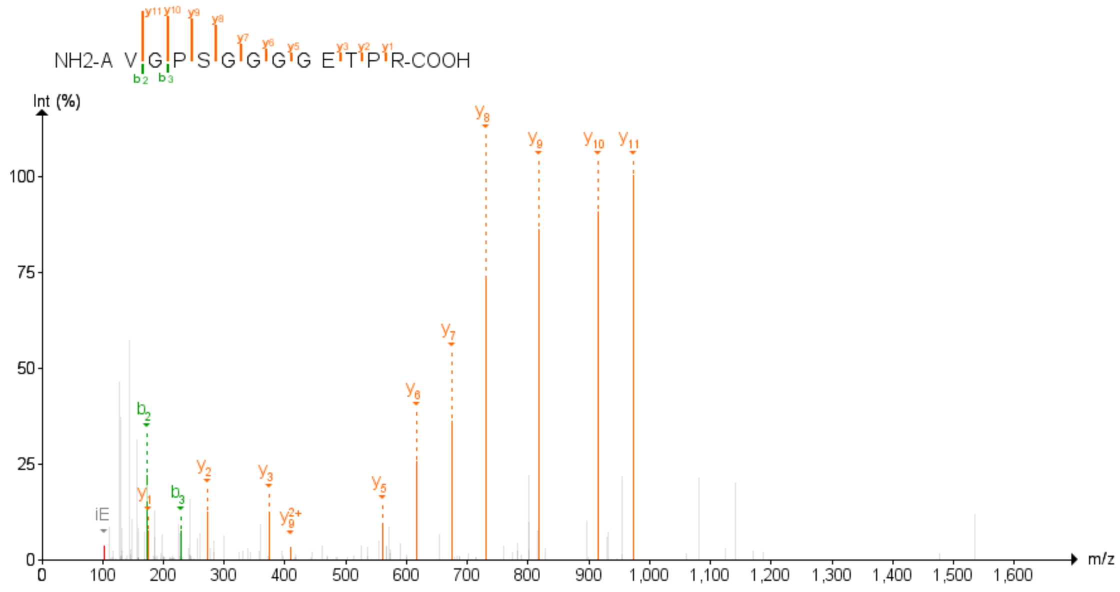

Figure 100: TMEM245 Third experiment

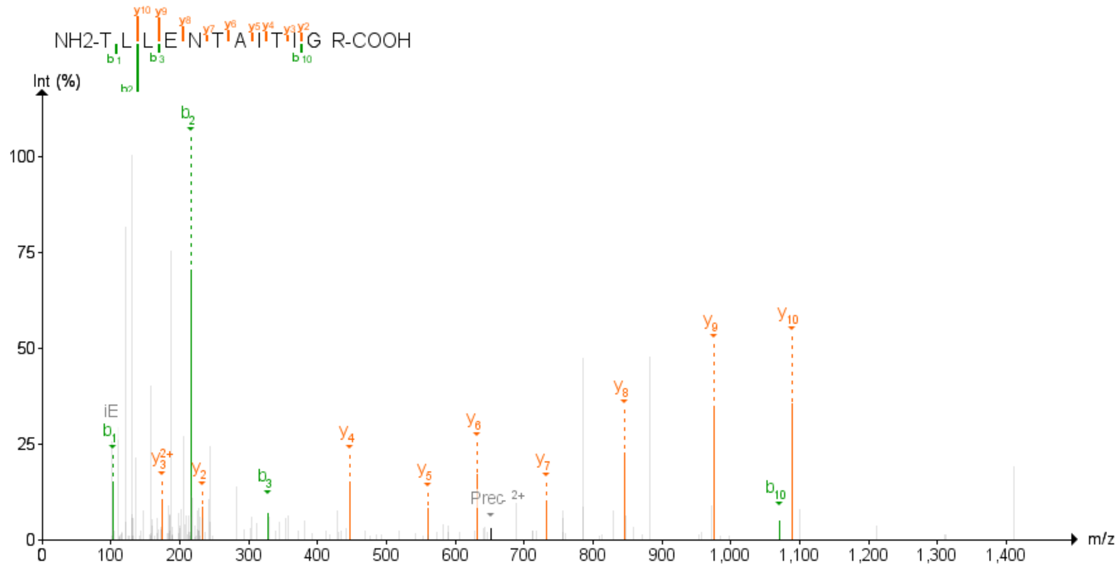

Figure 101: TNPO1 First experiment

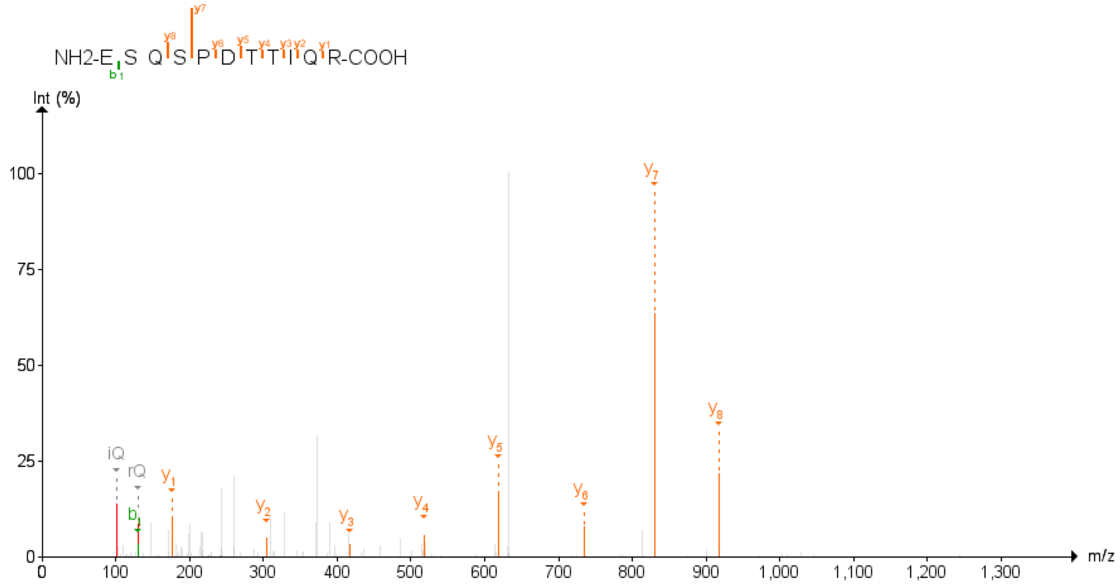

Figure 102: TNPO1 Second experiment

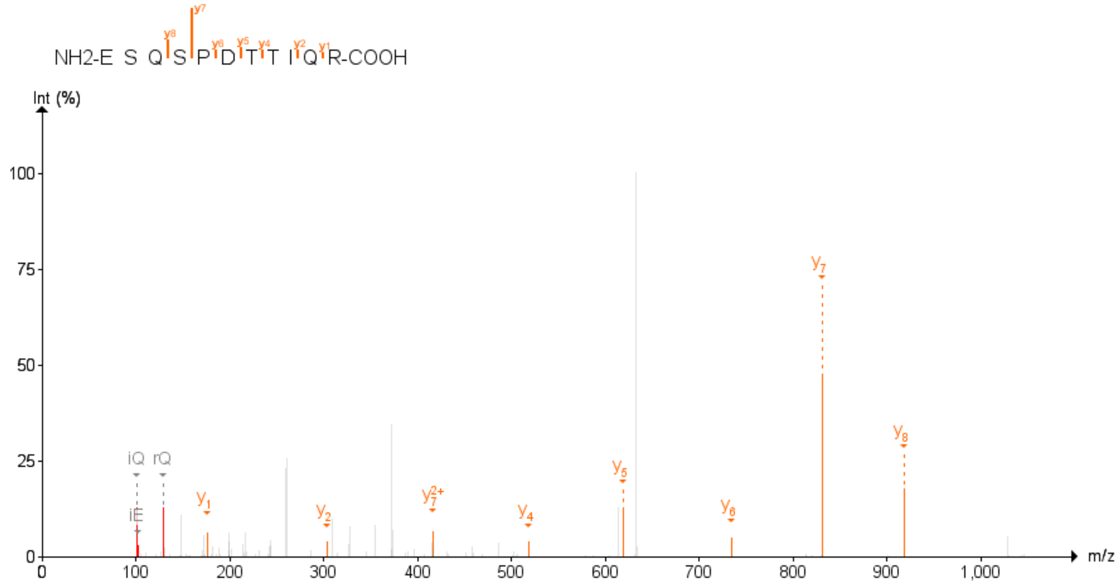

Figure 103: TNPO1 Third experiment

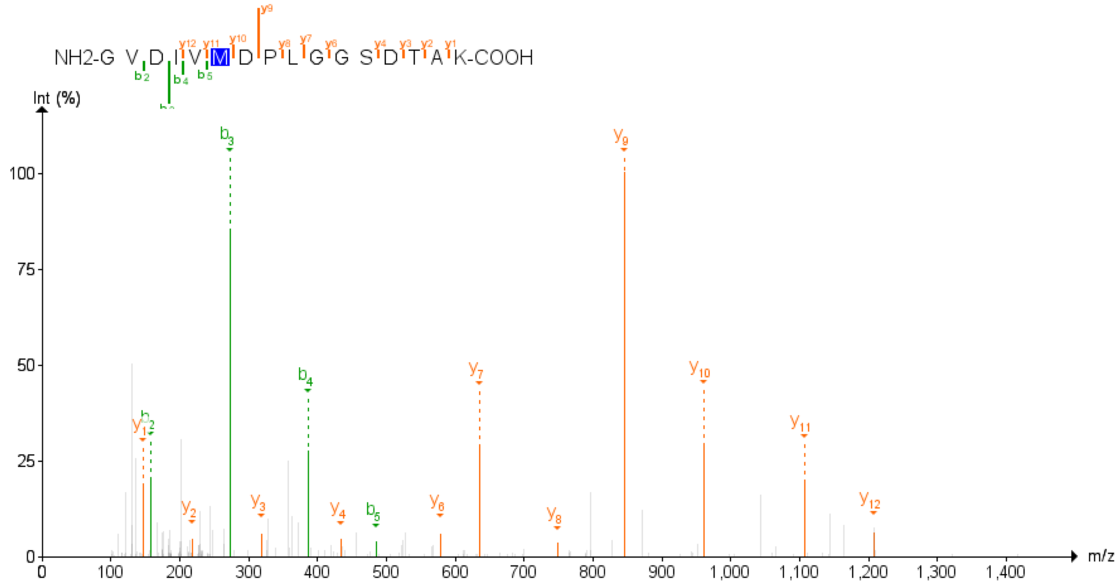

Figure 104: VAT1 Second experiment

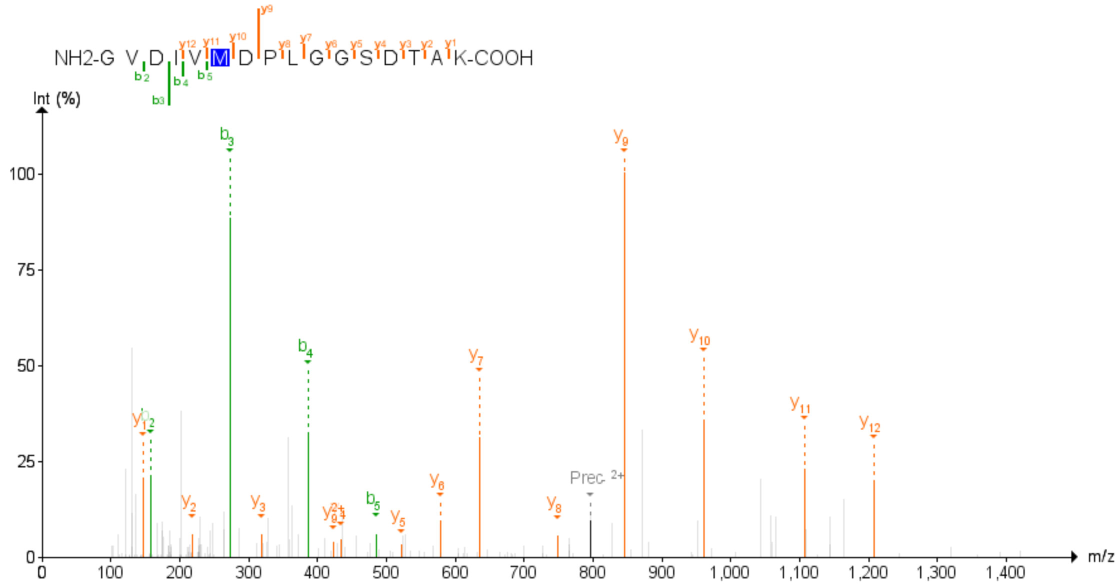

Figure 105: VAT1 Third experiment
